# Oxylipin metabolism is controlled by mitochondrial β-oxidation during bacterial inflammation

**DOI:** 10.1101/2020.08.17.252007

**Authors:** Mariya Misheva, Konstantinos Kotzamanis, Luke C Davies, Victoria J Tyrrell, Patricia R S Rodrigues, Gloria A Benavides, Christine Hinz, Robert C Murphy, Paul Kennedy, Philip R Taylor, Marcela Rosas, Simon A Jones, James E McLaren, Sumukh Deshpande, Robert Andrews, Nils Helge Schebb, Magdalena A Czubala, Mark Gurney, Maceler Aldrovandi, Sven W Meckelmann, Peter Ghazal, Victor Darley-Usmar, Daniel A White, Valerie B O’Donnell

**Affiliations:** Systems Immunity Research Institute and Division of Infection and Immunity, and School of Medicine, Cardiff University, CF14 4XN, UK; Department of Pathology, University of Alabama at Birmingham, Birmingham, AL 35294, USA; Department of Pharmacology, University of Colorado Denver, Aurora, CO 80045, USA; Cayman Chemical 1180 E Ellsworth Rd, Ann Arbor, MI 48108, United States; UK Dementia Research Institute at Cardiff, Cardiff University, CF14 4XN, UK; University of Wuppertal, Chair of Food Chemistry, Faculty of Mathematics and Natural Sciences, Gausstraße 20, 42119 Wuppertal, Germany

## Abstract

Oxylipins are potent biological mediators requiring strict control, but how they are removed en masse during infection and inflammation is unknown. Here we show that lipopolysaccharide (LPS) dynamically enhances oxylipin removal via mitochondrial β-oxidation. Specifically, genetic or pharmacological targeting of carnitine palmitoyl transferase 1 (CPT1), a mitochondrial importer of fatty acids, reveal that many oxylipins are removed by this protein during inflammation in vitro and in vivo. Using stable isotope-tracing lipidomics, we find secretion-reuptake recycling for 12-HETE and its intermediate metabolites. Meanwhile, oxylipin β-oxidation is uncoupled from oxidative phosphorylation, thus not contributing to energy generation. Testing for genetic control checkpoints, transcriptional interrogation of human neonatal sepsis finds upregulation of many genes involved in mitochondrial removal of long-chain fatty acyls, such as *ACSL1,3,4, ACADVL, CPT1B, CPT2 and HADHB*. Also, *ACSL1/Acsl1* upregulation is consistently observed following the treatment of human/murine macrophages with LPS and IFN-γ. Last, dampening oxylipin levels by β-oxidation is suggested to impact on their regulation of leukocyte functions. In summary, we propose mitochondrial β-oxidation as a regulatory metabolic checkpoint for oxylipins during inflammation.

## Introduction

Oxygenated polyunsaturated fatty acids (oxylipins) are essential bioactive lipid mediators during inflammation/infection. There are generated by cyclooxygenases (COX), lipoxygenases (LOX) or cytochrome P450s (CYP), expressed in a variety of cells and tissues [1–10]. They signal through activation of G protein-coupled receptors at sub nM concentrations [1, 5–16]. Oxylipin signaling requires deactivation, however our understanding of how this happens during infection is poor. Systemic pathways for individual oxylipins, including thromboxane, prostacyclin and hydroxyeicosatetraenoic acids (HETEs) were uncovered in healthy humans decades ago [17–20]. There, infusion of exogenous labelled lipids enabled determination of half-life and metabolites, some of which appear immediately, slowly disappearing over several minutes [21]. Urinary metabolite analysis became the gold standard for whole body oxylipin analysis. Separately, peroxisomal β-oxidation of individual oxylipins was explored using liver microsomes. There, partial β-oxidation revealed truncated products termed dinors (minus 2 carbons) and tetranors (minus 4 carbons). These were identified as stable intermediates in tissue, plasma and urine [21].

Aside from peroxisomes, mitochondria contain fully competent β-oxidation machinery, used for the first steps of fatty acid (FA)-dependent energy metabolism, FA oxidation (FAO) [22, 23]. Here, FA are converted to acetyl-CoA which enters the tricarboxylic cycle (TCA) providing substrates for oxidative phosphorylation (OxPhos) [24]. Although mediated by distinct enzymes to peroxisomes, mitochondrial β-oxidation also involves sequential removal of 2-carbon fragments from the carboxyl terminus. More recently, there has been significant interest in how mitochondrial FAO supports innate and adaptive immunity in T cells and macrophages[25, 26]. Saturated FA such as palmitate or stearate are considered the main β-oxidation substrates, however oxylipins generated abundantly during inflammation have not been considered [19]. We recently showed that endogenously-generated platelet oxylipins are removed by carnitine palmitoyltransferase-1 (CPT1), the mitochondrial import protein for FAs [27]. However, it is not known whether mitochondria also remove oxylipins in other leukocytes, nor how this might be regulated during inflammation/infection, if this impacts lipid signaling, and if it supplies acetyl-CoA to OxPhos. These are important questions, since large amounts of diverse oxylipins are generated during inflammation/infection, with downstream autocrine and paracrine signaling being a major contributor to the overall inflammatory response. Currently there is significant interest in how mitochondria contribute to inflammation, and their role in immunometabolism[26, 28, 29]. One view is that in lipopolysaccharide (LPS) treated-macrophages (e.g. M1 phenotype) mitochondrial OxPhos is shut down in favor of increased glycolysis, along with increased synthesis of FA [30]. How this process impacts the bioavailability oxylipins and their inflammatory bioactivity is unknown.

To address these questions, we examine the role of mitochondria in removing oxylipins in macrophages during inflammatory activation *in vitro* and *in vivo*, and how modulation of physiological oxylipin levels impacts leukocyte signaling relevant to an acute inflammatory challenge (macrophages, neutrophils, T cells). Genetic and lipidomic approaches characterize a metabolic β-oxidation network for oxylipins and their metabolites, during inflammation. The study identifies a mechanism for dampening oxylipin signaling that could be more broadly relevant in inflammation and infection.

## Results

### Inhibition of CPT1 prevents secretion of diverse peritoneal oxylipins in vivo

Mitochondrial β-oxidation of native long chain FA requires uptake of their FA-CoA derivatives *via* the outer membrane transporter, CPT1, of which there are three isoforms, a,b,c. However, whether oxylipin species can be imported via CPT1 in macrophages is unknown. To test this *in vivo*, the pan CPT1 inhibitor etomoxir was injected intraperitoneally (i.p.) into wild type C57BL/6 mice alone or with the bacterial lipid LPS (1 μg), then peritoneal lavage harvested after 6 hrs, and analyzed using LC/MS/MS. A large number of oxylipins were detected in cell-free supernatant with several showing expected elevations in response to LPS (Figure 1 A,B, Supplementary Figure 1 A). These are broadly categorized into lipids generated by 12/15-LOX, COX-1/2 or CYP450, although some are from more than one pathway (Supplementary Data File 1)). Many specialist pro-resolving mediators (SPM) including resolvins or protectins were not conclusively detected. For maresin1 (Mar1, 7*R*,14*S*-diHDOHE), very small peaks were seen that fell below our LOQ for the assay. When injecting larger amounts of sample and broadening the retention time window, two peaks were seen with one having the same retention time as the authentic standard (Supplementary Figure 1 B). It wasn’t possible to generate a convincing MS/MS spectrum from tissues that matched the standard due to the low levels present. The detection of two isomers suggests diastereomers of 7,14-di-HDOHE, and the peaks may also contain co-eluting stereoisomers. This is suggestive of non-enzymatic origin and this peak is more appropriately named 7,14-diHDOHE.

**Figure 1.**
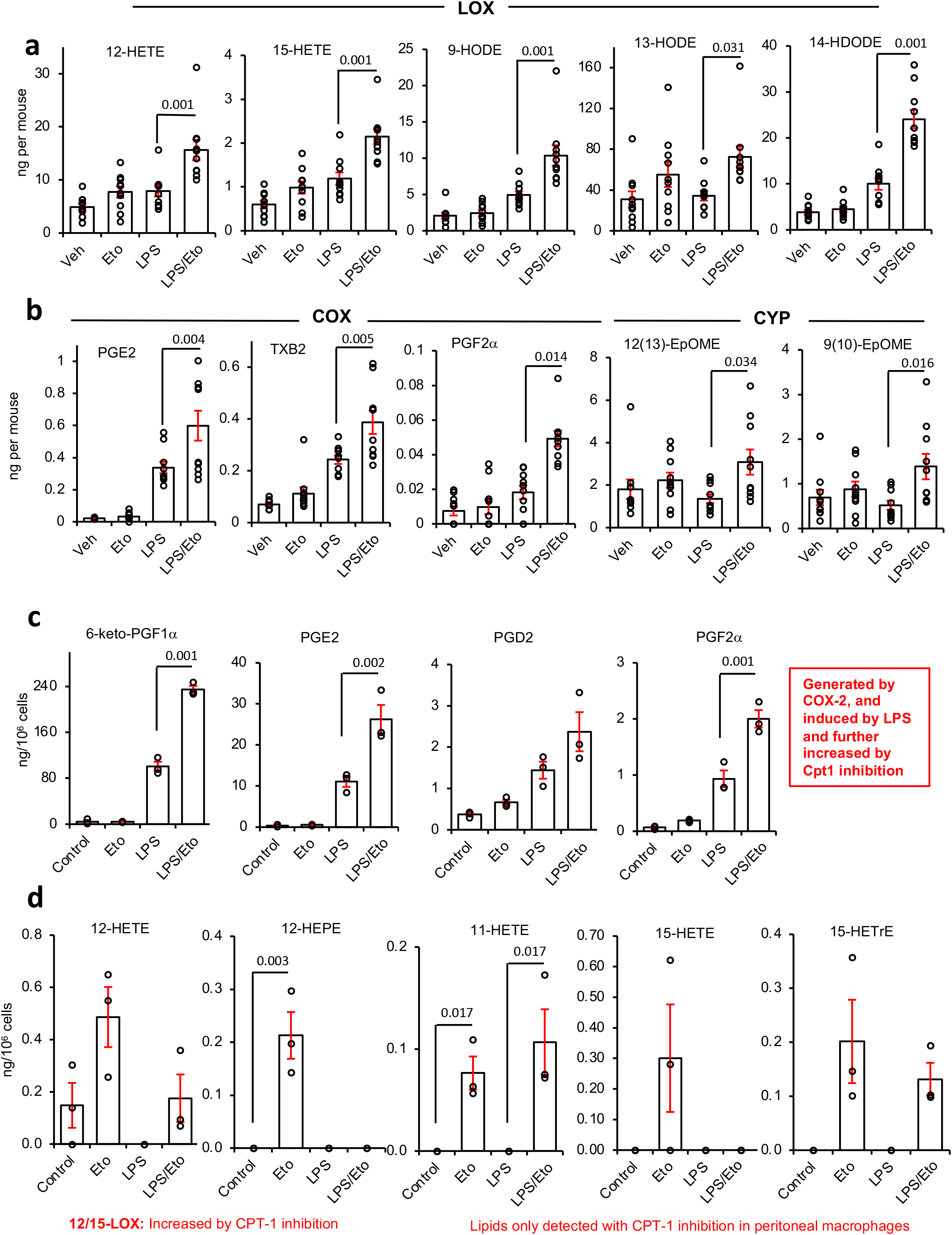
Inhibition of CPT1 increases oxylipin levels significantly during inflammation *in vivo* and in peritoneal macrophages. *Panels A, B. Plots for individual lipids showing that inhibition of CPT1 enhances oxylipin levels, with a bigger impact during inflammatory challenge with LPS*. Wild type mice (female, 7-9 weeks) were injected i.p. with vehicle (PBS), etomoxir (100 μg) or LPS (1 μg). After 6 hrs, lavage was harvested and lipids extracted using SPE then analyzed using LC/MS/MS as outlined in Methods. A series of example lipids from 12/15-LOX, COX and CYP450 are shown (n = 10). *Panels C-D Plots for individual lipids showing impact of etomoxir +/- LPS on oxylipin release by peritoneal macrophages in vitro.* Peritoneal macrophages were isolated as described in Methods, then cultured in serum-free medium in the presence of etomoxir (25 μM) with/without LPS (100 ng/ml). After 24 hrs, supernatant was harvested and lipids extracted using SPE, then analyzed using LC/MS/MS. A series of lipids based on enzymatic origin are shown, n = 3 per sample. For all panels, data is mean +/- SEM, one way ANOVA with Tukey post hoc test, stats are shown for effect of etomoxir only, where significant. Where no bar is shown no significant difference was seen.

Etomoxir stimulated small increases in several oxylipins, however when included with LPS, a far stronger impact was seen, and levels of many lipids were effectively doubled (Figure 1 A,B). This suggests that CPT1 in peritoneal mitochondria actively transports bioactive oxylipins during health, but in infection, its role is significantly enhanced.

### CPT1 regulates metabolism of endogenously-generated oxylipins by peritoneal macrophages

*In vivo*, peritoneal oxylipins could be metabolized by mitochondria in resident macrophages, B cells and/or peritoneal membrane (mesothelium). Naïve resident macrophages, which express COX-1 and 12/15-LOX (*Alox15*), are likely to be the primary basal source [31]. Thus, naïve peritoneal macrophages from wild type mice were assessed for mitochondrial removal of endogenous oxylipins. We focused on extracellular forms, since most are secreted to activate G protein coupled receptors (GPCRs) extracellularly. Serum free media was used to prevent contamination with blood derived oxylipins. LPS (24 hrs) stimulated prostaglandin (PG) secretion by naïve resident peritoneal macrophages, consistent with COX-2 induction (Figure 1 C, Supplementary Figure 1 C). Although 12/15-LOX is highly expressed in naïve peritoneal macrophages, its loss in culture leads to relatively low generation of monohydroxy FA (12-HETE and 12/15-HEPEs) (Figure 1 D, Supplementary Figure 1 D) [32]. Overall, oxylipin species generated by peritoneal macrophages were similar to those in lavage, although the total number/diversity was less with fewer CYP-derived species and no SPMs detected (Supplementary Figure 1 C).

Etomoxir significantly increased detected levels of several mono-hydroxy lipids and COX-2 products (Figure 1 C,D). This indicates that both their formation and removal is increased by inflammation, similar to *in vivo* (Figure 1 A,B). 12-HETE, 12-HEPE and 15-HEPE increased with etomoxir but were suppressed by LPS, likely due to inflammation-associated loss of 12/15-LOX (Figure 1 D, Supplementary Figure 1D). Importantly, *in vivo* and *in vitro*, COX-2 and 12/15-LOX products were consistently increased by CPT1 inhibition.

Several mono- and di-hydroxy oxylipins were only detected when CPT1 was blocked, indicating that their mitochondrial metabolism exceeds their generation (Figure 1 D, Supplementary Figure 1D). Thus, CPT1 prevents their secretion. In contrast, during LPS challenge, the eicosapentaenoic acid (EPA) product 17,18-diHETE, generated by soluble epoxide hydrolase oxidation of CYP-derived 17,18-EET, was suppressed by etomoxir (Supplementary Figure 1 E). Overall, these data indicate that peritoneal macrophage oxylipin secretion (12/15-LOX or COX-2 derived) is counterbalanced by CPT1-mediated uptake into mitochondria. However, CYP-derived lipids show the opposite, where CPT1 inhibition dampens levels.

### Bone marrow derived macrophages consume diverse oxylipins from serum

Macrophages are exposed to oxylipins from other immune or stromal cells during inflammation. Serum contains numerous oxylipins from 12-LOX and COX-1, primarily mono-hydroxy isoforms and thromboxane, generated by white cells and platelets. Serum contains few prostaglandins (PGs). Serum forms during innate immune responses, thus oxylipin metabolism may occur local or systemic sites. To examine this, we tested primary murine bone marrow-derived cells differentiated to macrophages, then treated with M-CSF, alone or along with LPS/IFNγ or IL-4 (henceforth referred to as M0, M1 and M2) respectively as *in vitro* of models for macrophage inflammation [33]. All macrophage phenotypes consumed significant amounts of exogenous mono-hydroxy oxylipins from 10 % serum-containing medium, including 12-HETE/12-HEPE (12-LOX, generated by platelets), 5-HETE/5-HEPE (5-LOX, generated by neutrophils) and 9-/13-HODEs (Supplementary Figure 2 A). 11-HETE/11-HEPE and 15-HETE/15-HEPE were removed by M0(M-CSF) and M2(IL-4) cells, however they appeared higher or unchanged for M1(LPS/IFNγ), since they were simultaneously generated by COX-2, induced by LPS/IFN (Supplementary Figure 2 A). Furthermore, M1(LPS/IFNγ) cells secreted large amounts of PGs and thromboxane due to COX-2, causing a net increase in their extracellular levels. (Supplementary Figure 2 A). Thus, the overall pattern was consumption of serum oxylipins by all macrophage populations, coupled with simultaneous generation of COX-2 PGs by M1(LPS/IFNγ). Last, low dose etomoxir (25 M) significantly increased PGs released by M1(LPS/IFNγ) cells (Supplementary Figure 2 B). Overall, this demonstrates that mitochondrial uptake of PGs suppresses their secretion from classic inflammatory BMDM cells (M1(LPS/IFNγ)), in the same manner as seen with naïve peritoneal macrophages and *in vivo* with LPS peritonitis, while all BMDM cell types consume significant amounts of HETEs and HEPEs.

### Dynamic control of oxylipins by CPT1 during inflammatory challenge in RAW macrophages

Next, we used RAW cells, a model for macrophage inflammatory responses *in vitro*, amenable to genetic modification. These don’t express 12/15-LOX, therefore, to examine the impact of CPT1 on its products, cells stably overexpressing *Alox15* (RAW*Alox15*) were generated. Basally, neither RAW^mock^ nor RAW*Alox15* cells secreted many oxylipins, with only 9/-HODE, two PGs, and three di-hydroxy products of CYP released in low amounts (Supplementary Figure 3 A). LPS (24 hrs, 100 ng/ml) stimulated robust secretion of many oxylipins, including several PGs (COX-2) 9/13-HODEs, and 11-and 15-HETEs, HETrEs and HEPEs (likely from COX-2) (Figure 2 A, Supplementary Figure 3 A,B, 4 A-C). The primary 12/15-LOX products 12-HETE and 14-HDOHE were only detected in RAW*Alox15* cells, and their generation was increased by LPS (Figure 2 A).

**Figure 2.**
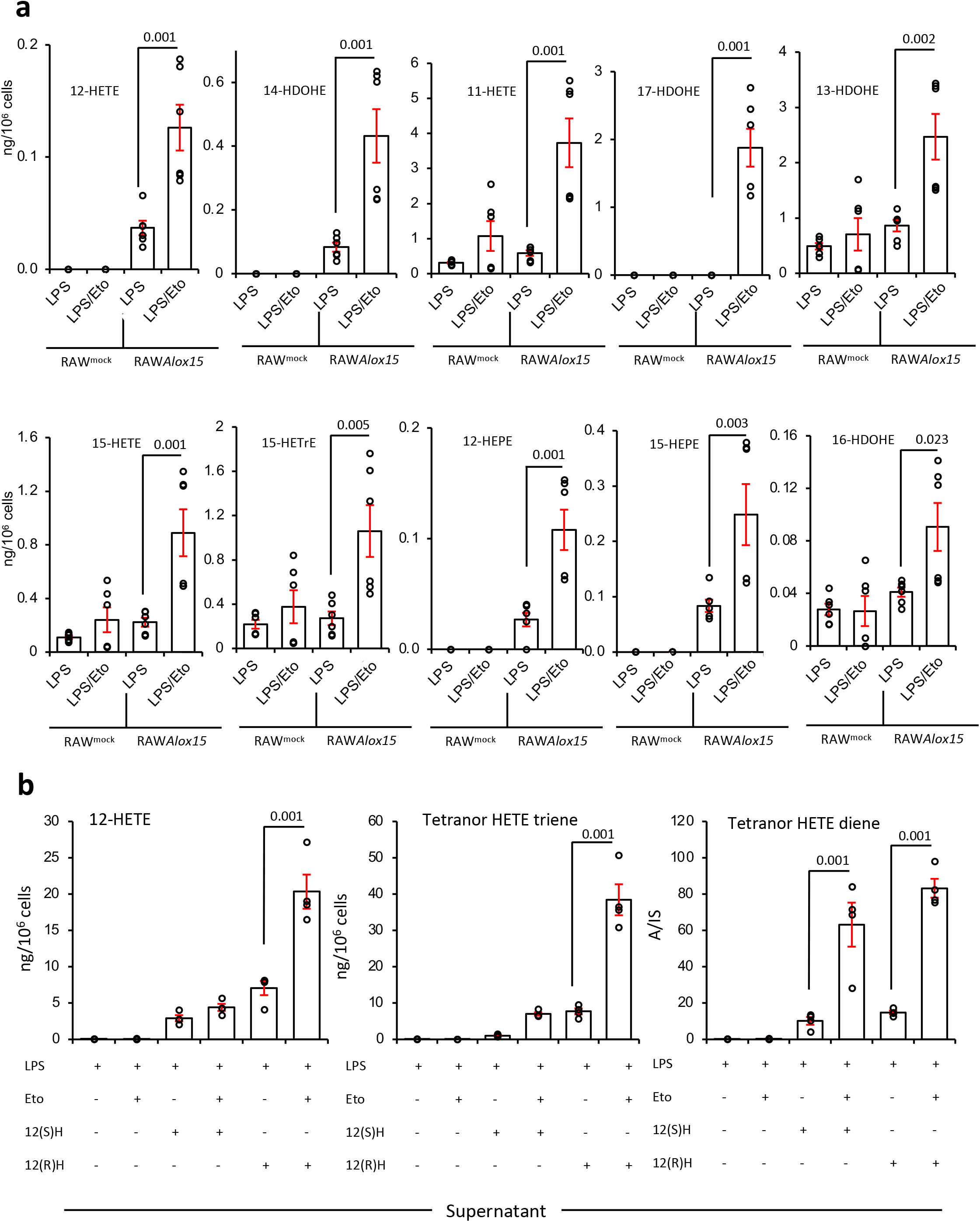
Inhibition of CPT1 increases secretion of 12/15-LOX-derived eicosanoids from RAW cells overexpressing *Alox15*, while RAW cells rapidly consume exogenous 12-HETE, secreting two tetranor metabolites which are themselves metabolized by mitochondrial β-oxidation. *Panel A. Blockade of CPT1 leads to significant elevation of 12/15-LOX-derived oxylipins in RAW cells*. RAW^mock^ and RAW*Alox12* cells were cultured with etomoxir (25 μM) and/or LPS (100 ng/ml) for 24 hr before harvest and SPE extraction of supernatant for oxylipin analysis using LC/MS/MS. Bar charts showing how oxylipins from 12/15-LOX are impacted by etomoxir (n = 6, separate wells of cells, mean +/- SEM). *Panel B. CPT1 inhibition prevents metabolism of 12(S) or 12(R)HETE and their tetranor triene and diene metabolites*. RAW cells were supplemented with 1.5μg 12(S) or 12(R)-HETE/10^6^ cells for 3 hrs with/without etomoxir (25 μM), then supernatants were analyzed for levels of 12-HETE and its triene and diene tetranor products using LC/MS/MS (n = 3 (LPS alone) or 4 (all other samples), mean +/- SEM, separate wells of cells). For all panels, comparisons are with/without etomoxir, one way ANOVA with Tukey post hoc test, stats are shown for effect of etomoxir only, where significant.

CPT1 inhibition had little impact on basal oxylipin secretion (Supplementary Figure 3 A). However, it significantly elevated LPS-dependent generation of mono-hydroxy FA (HETEs, HDOHEs, HETrEs and HODEs) from RAW*Alox15* cells (Figure 2 A). These data indicate that secreted oxylipins normally represent only a fraction of the overall amounts made by 12/15-LOX. Thus, LPS increases CPT1-mediated removal of 12/15-LOX products, similar to *in vivo* and peritoneal macrophages. Many primary 12/15-LOX products (12-HETE, 14-HDOHE) were seen, while others (9-HOTrE, 9-HODE) may be biproducts of secondary propagation. In etomoxir/LPS-treated RAW*Alox*15 (but not RAW cells), a lipid suggestive of resolvinD5 (RvD5, 7S,17S-diHDOHE) was detected, however of the two peaks, only one matched the retention time of the authentic standard (Supplementary Figure 4 A,B). The lipid was very low abundance, and it was not possible to generate a reliable MS/MS spectrum from cell supernatants. As there were two peaks, they likely represent diastereomers of 7,17-di-HDOHE, and the individual peaks may also contain co-eluting stereoisomers. Thus, we labelled this 7,17-diHDOHE and suggest it arises from non-enzymatic oxidation. No SPMs were detected in any other samples from RAW or RAW*Alox15* cells.

Several COX-2 derived PGs and 14,15-diHETE (both CYP) were significantly decreased by etomoxir in RAW cells (Supplementary Figure 4 C). This mirrored the impact of CPT1 inhibition on peritoneal macrophage 17,18-diHETE. In summary, while LOX products were consistently elevated by etomoxir in RAW or all primary macrophages, for RAW cells alone, both COX and CYP-derived lipids were suppressed.

### Exogenous 12-HETE is metabolized to diene and triene tetranor metabolites via mitochondrial and non-mitochondrial β-oxidation in RAW cells

To determine the fate of oxylipins removed by CPT1, we examined metabolism of exogenous 12-HETE by RAW macrophages. 12(S)-HETE was rapidly removed following LPS-stimulation, coinciding with formation and subsequent metabolism of two tetranor 12-HETEs (Supplementary Figure 4 D-F). These were confirmed by MS/MS as a triene, 8-hydroxy-4Z,6E,10Z-hexadecatrienoic acid (comparison with authentic standard), and a diene, proposed as 8-hydroxy-6,10-hexadecadienoic acid (through comparison with [34]) (Supplementary Figure 5 A-C). Both will be 8(S) since they originate from 12(S)-HETE. As confirmation, 12(S)-HETE-d8 was added to RAW cells (3 hrs), and a deuterated diene was detected (Supplementary Figure 5 D,E). The MS/MS fragmentation of these lipids is shown (Supplementary Figures 12-14). For the tetranor diene, fragmentation matched expected gas phase chemistry of hydroxylated FAs, where cleavage occurs next to the hydroxyl group (Supplementary Schemes 13,14). For the triene, a daughter ion at *m/z* 165.1 was seen, with high resolution MS/MS of the standard confirming this as C_11_H_17_O (Supplementary Figure 6 A, Supplementary Scheme 12). NMR for the triene is shown in Supplementary Figure 7. Almost all triene and diene was detected extracellularly, and over 4-8 hrs, both disappeared from supernatant and cell pellets (Supplementary Figure 4 E,F). This suggests that macrophages generate and secrete primary and secondary metabolites of HETEs, but then re-internalize them for further metabolism. We next incubated triene tetranor 12(S)-HETE with RAW macrophages, and after 3 hrs, diene was detected extracellularly (Supplementary Figure 6 B). This indicates that diene forms via saturation of the triene, confirming the structure as 8(S)-hydroxy-6E,10Z-hexadecadienoic acid.

Next the impact of CPT1 inhibition on metabolism of exogenous 12(S)-HETE or 12(R)HETE was tested. Both HETEs were rapidly removed by LPS-stimulated RAW cells, with around 0.15% or 0.4% remaining after 3 hrs, for the S and R forms respectively (Figure 2 B, left panel). Cellular and supernatant 12(S)-HETE and 12(R)HETE were both increased by CPT1 inhibition, with 12(R)HETE most strongly impacted. Based on the impact of etomoxir, the primary removal appeared to be non-mitochondrial (Figure 2 B). 12-HETE was converted to tetranor trienes and dienes, which were detected mainly outside the cells (Figure 2 B, Supplementary Figure 8). These were around 5-8-fold increased by CPT1 blockade, confirming that they are also dynamically metabolized by mitochondria in LPS-stimulated macrophages (Figure 2 B, Supplementary Figure 8). In the case of the diene, both S and R enantiomers were similarly elevated by etomoxir treatment, while for the triene, the 12(R)HETE was most strongly impacted.

Since most removal appeared to be non-mitochondrial, we analysed for esterification of 12(S)-HETE into phospholipid pools via Land’s cycle, as previously described [35]. First, to determine which molecular species were formed, a precursor scanning LC/MS/MS analysis was undertaken, scanning for negative ion precursors of *m/z* 319.2, the carboxylate anion of HETEs. Cells were supplemented with 12(S)-HETE for 3 hrs, then harvested and lipid extracts were analysed. Several precursor ions were found, between *m/z* 738-810 which were absent in RAW cells not supplemented with 12-HETE (Figure 3 A,B). Their structures were subsequently confirmed using MS/MS, showing they originate from a series of expected HETE-containing PE or PC lipids using MS/MS scanning. Next, these lipids were quantified using LC/MS/MS. Substantial amounts of 12-HETE-PE/PC were generated during the 3 hr incubation with 12(S)-HETE (1.4 μg/10^6^ cells), totaling ca. 88% of the added oxylipin. When cells were concurrently stimulated with LPS, the proportion of HETE incorporated reduced to ca. 73%, with significant reductions in levels of three lipids and trends towards a reduction in all (Figure 3 C). This identifies esterification of 12-HETE into PLs to be the dominant route of removal in RAW cells, accounting for considerably more than β-oxidation and is in line with previous work by Pawlowski et al [35]. The partial dampening of this phenomenon by LPS may be a result of phospholipase activities being upregulated during inflammation. Further studies are needed to determine the impact of inflammation on all competing pathways of oxylipin removal in order to build up a more comprehensive picture of which are the dominant events under defined conditions relevant to specific situations such as inflammation, wounding, coagulation, etc.

**Figure 3.**
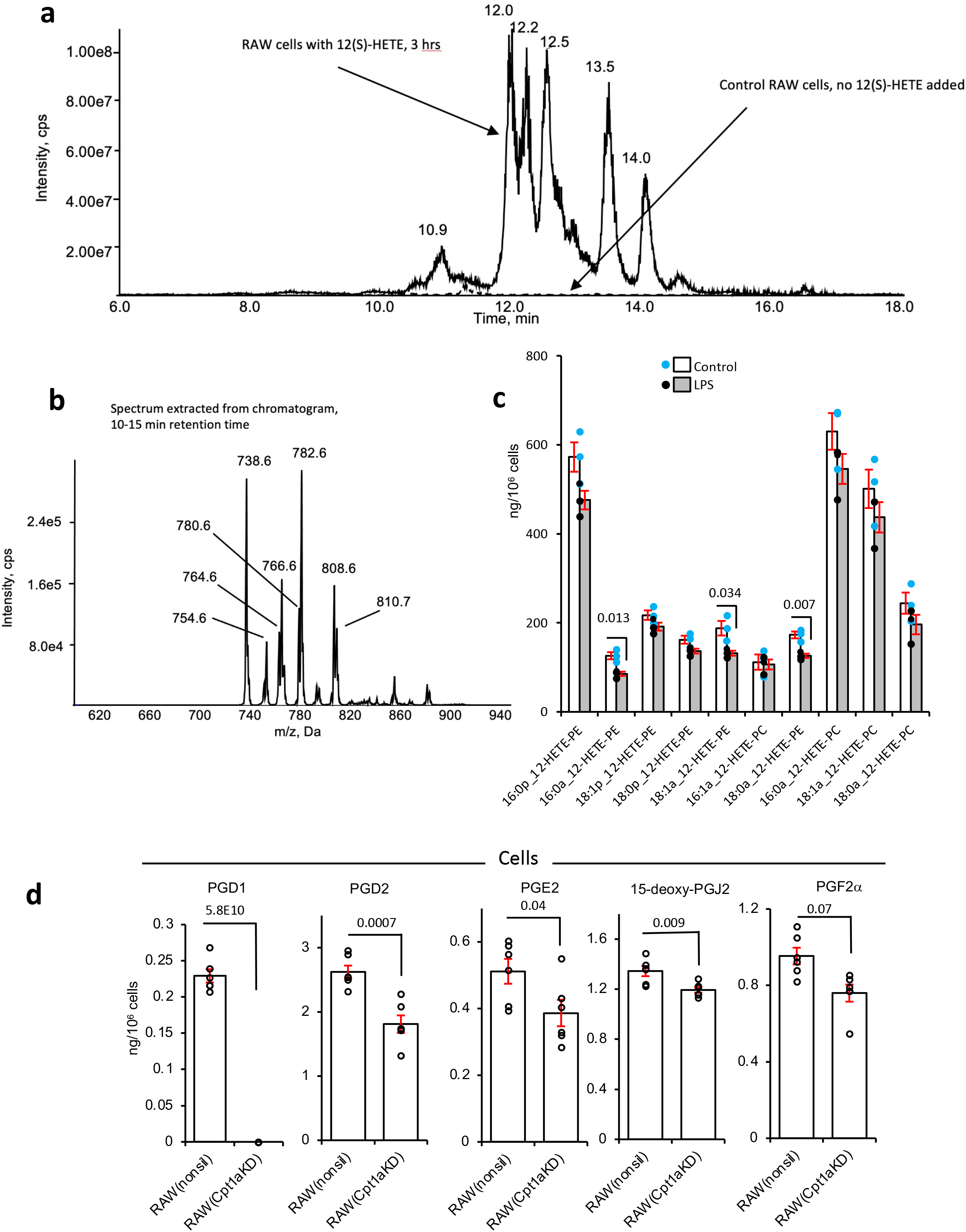
Identification of phospholipid-esterified 12-HETE in RAW cells, and CPT1 knockdown reduces PG levels in RAW cells. *Panels A-C.* Cells (10^6^) were incubated for 3 hours with 12(S)-HETE +/- LPS before lipids were extracted and analysed as in Methods. Precursor LC/MS/MS was undertaken as described in Methods, scanning for ions that fragment to generate HETE. *Panel A.* Chromatogram showing elution of precursors that generate product ions with *m/z* 319.2. *Panel B.* MS spectrum from 10-15 min showing PE and PC species that contain 12-HETE. *Panel C*. Quantification of 12-HETE PE and PC species that are formed following incubation of RAW cells with 12(S)-HETE. (n = 3, mean +/- SEM, separate wells of cells) compared with/without LPS, Student’s T-test, two-tailed. Where no bar is shown no significant difference was seen. *Panel D. Cpt1a knockdown dampens cellular levels of PGs in RAW cells*. RAW cells expressing either the non-silencing (RAW^nonsil^) or Cpt1a knockdown siRNA (RAW*Cpt1a^KD^*) were treated with LPS (100 ng/ml) for 24 hr and cell levels of PGs measured using LC/MS/MS as in Methods. (n = 6, mean +/- SEM, separate wells of cells). Student’s T-test, two-tailed. Where no bar is shown no significant difference was seen.

### CPT1a gene silencing confirms dysregulation of oxylipin secretion in RAW cells

Considerably higher amounts of etomoxir than we used (∼10-fold higher) can have off-target effects due to an impact on adenine nucleotide translocase [36, 37]. To rule this out, we targeted CPT1a using a lentiviral approach. Silencing to around 14 % control levels was achieved in RAW264.7 macrophages, stably expressing shRNA that targets *Cpt1a (*RAW*Cpt1a^KD^*), but not those expressing a non-silencing control (RAW^nonsil^) (Supplementary Figure 9 A-C). CPT1 maintains mitochondrial health since it imports FAs to maintain cardiolipin and phospholipid pools inside mitochondria [38]. Thus, CPT1-deficient cells do not proliferate normally and may alter their bioenergetic and lipid metabolism profiles. The balance between mitochondrial and peroxisomal β-oxidation and their ability to support elongase activities required to generate polyunsaturated FA (PUFA) for oxylipin synthesis may have adapted. Similar to low dose etomoxir, the overall impact on endogenous LPS-stimulated PG synthesis in RAW cells was suppression of supernatant and cell levels (Figures 3D, 4). Furthermore, the CYP450 product (17,18-diHETE) was suppressed, as also seen for etomoxir with naïve peritoneal macrophages and RAW cells. Endogenous generation of several monohydroxy eicosanoids was suppressed by silencing of *Cpt1a* (Figure 4). This data further confirms the dual role of CPT1 in both generation and removal of oxylipins depending on their enzymatic source and cell type.

**Figure 4.**
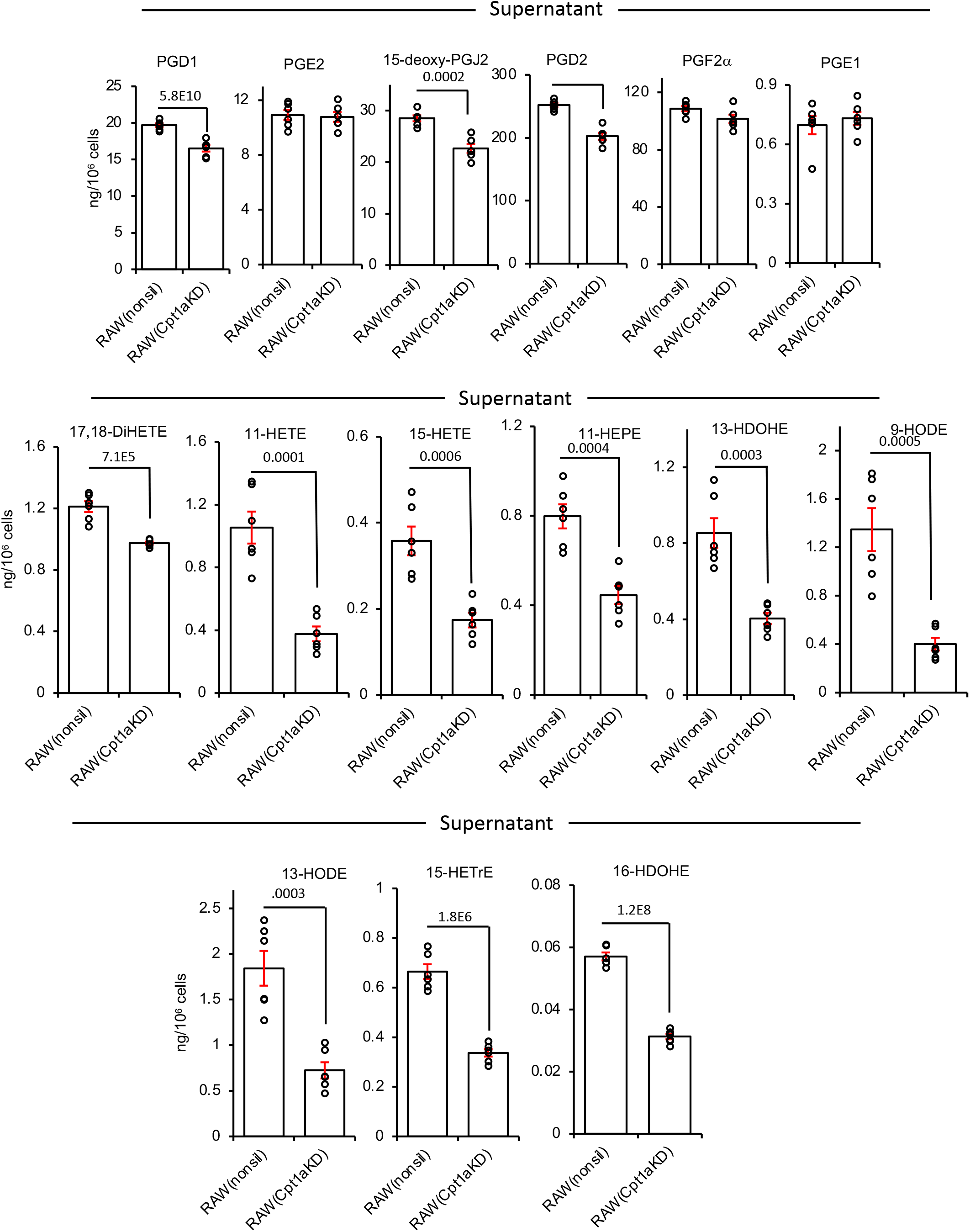
CPT1 genetic knockdown reduces secretion of oxylipins. RAW cells expressing either the non-silencing (RAW^nonsil^) or Cpt1a knockdown siRNA (RAW*Cpt1a^KD^*) were treated with LPS (100 ng/ml) for 24 hr and secretion of PGs, 17,18-diHETE or monohydroxy oxylipins were measured using LC/MS/MS as in Methods. (n = 6, mean +/- SEM, separate wells of cells). Student’s T-test, two-tailed. Where no bar is shown no significant difference was seen.

Since RAW cells do not generate 12-HETE, this metabolite was exogenously added to cultured cells to evaluate mitochondrial oxylipin removal. In basal RAW cells, *Cpt1* knockdown did not prevent metabolism of 12-HETE, instead there was a small increase, perhaps by esterification or peroxisomal metabolism (Figure 5 A). In contrast, 12-HETE removal by LPS-stimulated RAW cells was partially inhibited by *Cpt1* knockdown (Figure 5 A). Furthermore, the triene and diene tetranor metabolites were significantly reduced by *Cpt1* knockdown in LPS-treated but not basal RAW cells (Figure 5 B,C). This indicated that following *Cpt1* knockdown, RAW cells showed reduced mitochondrial metabolism of 12-HETE coupled with lower formation of its tetranor products post-LPS stimulation.

**Figure 5.**
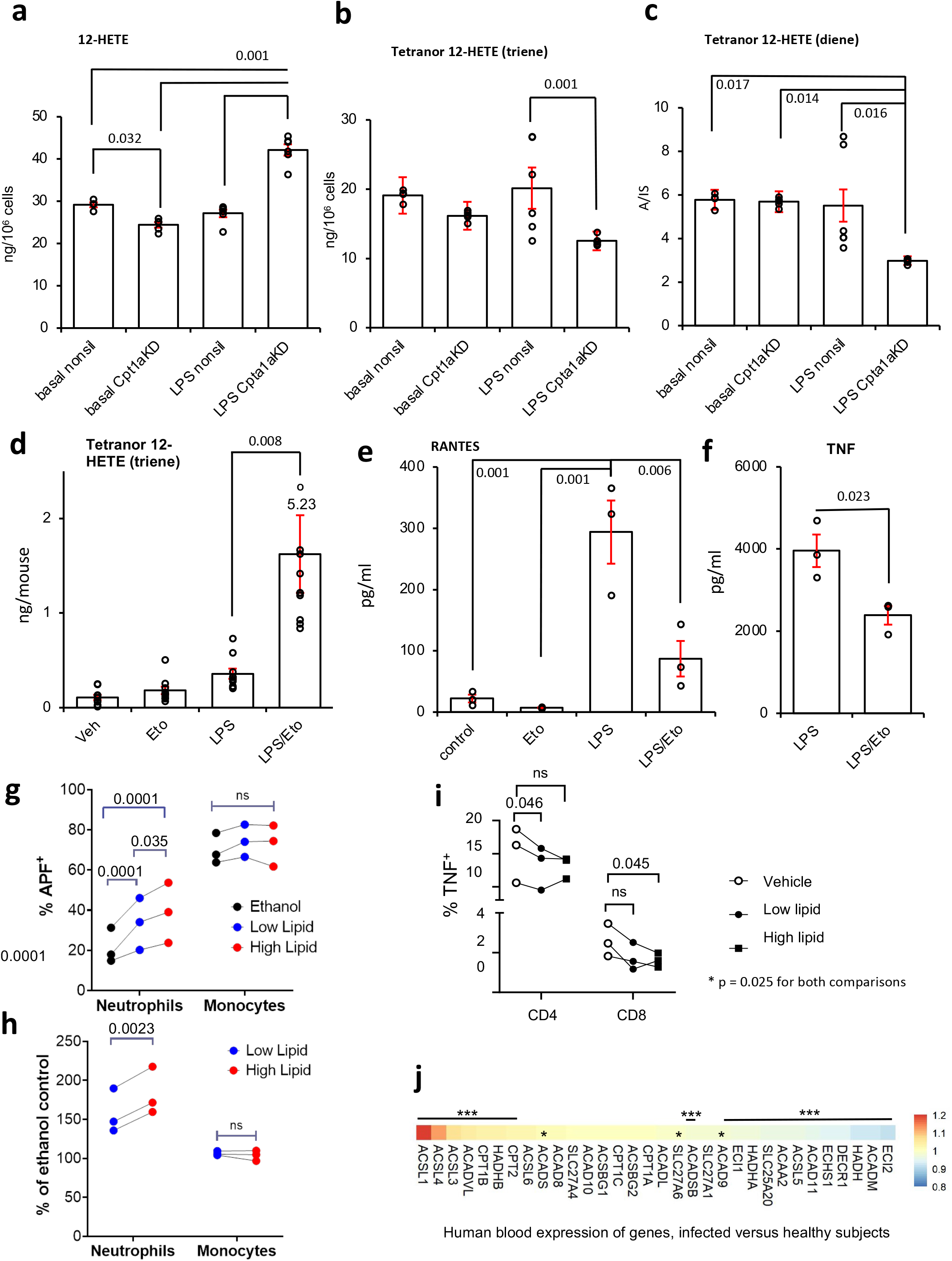
*Cpt1a* genetic knockdown modulates metabolism of 12-HETE and its metabolites by RAW cells, etomoxir reduces metabolism of tetranor triene *in vivo,* and prevents LPS stimulation of RANTES/TNFα generation, oxylipins regulate leukocyte responses, and transcriptomics of human neonatal sepsis proposes multiple genes that support a mitochondrial oxylipin β-oxidation pathway. *Panels A-C. knockdown of Cpt1a alters 12-HETE metabolism.* RAW cells were incubated +/- LPS (100 ng/ml, for 4 hr) with 12-HETE (2.34 µg/ml) added after the first hour (thus added for 3 hr). Supernatants were harvested, then lipids extracted and analyzed for 12-HETE and metabolites using LC/MS/MS as described in Methods (n = 4 (basal nonsil), 5 (basal Cpt1aKD) or 6 (other groups), separate wells of cells, mean +/- SEM). *Panel D. Tetranor triene 12-HETE is increased in vivo during inflammation with CPT1 inhibition.* Wild type mice (female, 7-9 weeks) were injected i.p. with vehicle (PBS), etomoxir (100 μg) and/or LPS (1 μg). After 6 hrs, lavage was harvested and lipids extracted and analyzed using LC/MS/MS as outlined in Methods (n = 10, individual mice). Outliers are shown as red, or with values. *Panels E,F. Etomoxir dampens generation of RANTES and TNF by peritoneal macrophages.* Supernatants from macrophages cultured in vitro for 24 hrs with LPS (100 ng/ml)/etomoxir (25 μM) were tested for RANTES/CCL5 or TNF using ELISAs as described in Methods (n = 3 per group, separate wells of cells, mean +/- SEM). For Panels A-F, * p <0.05, ** p < 0.01, *** p < 0.005 with/without etomoxir, one way ANOVA with Tukey post hoc test was used. *Panels G,H. Oxylipin levels detected in vivo promote neutrophil ROS generation.* Whole blood was incubated with oxylipin mixtures (Supplementary Table 2), and then ROS generation in response to stimulation using opsonized *S. epidermidis was* determined by APF fluorescence as indicated in Methods. Three individual donor samples were separately tested. Monocytes and neutrophils were analyzed using gating strategy as outlined in Methods. Response to high or low oxylipin doses were expressed either as %APF+ cells or % of the vehicle (ethanol) response. Each donor is shown separately. Data was analyzed by repeated measures two-way ANOVA (*G* – pairing p<0.0001, cell type p =0.018, lipid p=0.0002 and interaction p=0.0005; *H*– p pairing p=0.0008, cell type p =0.0094, lipid p =0.0082 and interaction p =0.0024), Sidak’s post hoc tests are indicated – G: Neutrophils Ethanol vs Low Lipid P=0.0001, Neutrophils Ethanol vs High Lipid P<0.0001, Neutrophils Low Lipid vs High Lipid P=0.0350, Monocytes Ethanol vs Low Lipid P=0.7020, Monocytes Ethanol vs High Lipid P=0.8075, Monocytes Low Lipid vs High Lipid P=0.8075 – H: Neutrophils Low Lipid vs High Lipid P=0.0023, Monocytes Low Lipid vs High Lipid P=0.2363). *Panel I. Oxylipin levels detected in vivo modulate CD4+ and CD8+ T cell activation*. Whole blood was incubated with oxylipin mixtures (Supplementary Table 2), and then generation of TNFα was determined in response to activation using CD3/CD28 beads as indicated in Methods. CD8+ and CD4+ T cells were analyzed using the gating strategy outlined in Methods. Data from individual donors is shown. Two-way ANOVA (repeated measures) was used, pairing p =0.0039, interaction p =0.1690, cell type p=0.0007, effect of lipid p =0.0253, Sidak’s post hoc tests are indicated - CD4 - No Lipid vs Low Lipid P=0.0461, CD4 - No Lipid vs High Lipid P=0.7115, CD4 - Low Lipid vs High Lipid P=0.9989, CD8 - No Lipid vs Low Lipid P=0.3742, CD8 - No Lipid vs High Lipid P=0.0449, CD8- Low Lipid vs High Lipid P=0.9952). . *Panel J. Human transcriptomics data shows significant upregulation of several genes including ACSL and CPT isoforms.* Gene expression (log2 expression data) from 35 controls and 26 bacterial confirmed sepsis cases published in [57] were compared for expression of 32 genes putatively involved in mitochondrial import and β-oxidation of oxylipins, based on known metabolic pathways for long chain native FA. * p < 0.05, *** p < 0.005, Students T-test, two-tailed, then adjusted using Benjamini-Hochberg test.

### Inflammation accelerates 12-HETE conversion to triene tetranor 12-HETE via mitochondrial β-oxidation in vivo

*In vivo* inhibition of CPT1 significantly elevated many oxylipins in the peritoneal cavity following LPS challenge. To examine β-oxidation of oxylipins via mitochondria *in vivo*, the formation of tetranor metabolites was next measured. The diene metabolite was absent, however tetranor 12-HETE triene was significantly elevated on CPT1 blockade during inflammatory stimulation (Figure 5 D). This confirms that inflammation *in vivo* not only increases formation and mitochondrial uptake of oxylipins (shown in Figure 1), but also their metabolism to tetranors.

### Oxylipin metabolism by mitochondria is not sustaining oxidative phosphorylation (OxPhos)

Mitochondrial β-oxidation of FA is directly linked with OxPhos. It forms acetyl-CoA, a substrate for the TCA cycle as well as NADH and FADH_2_, substrates for complexes I and II, and also transfers electrons to flavoprotein-ubiquinone oxidoreductase (ETF-QO) [39]. Thus, oxylipins could contribute to OxPhos. However, LPS suppresses mitochondrial respiration, through multiple mechanisms including aconitase inhibition and nitric oxide binding to complex IV, making this unlikely [28]. Here, LPS treatment profoundly decreased cellular respiration consistent with previous data (Supplementary Figure 9 D) [40, 41]. Importantly, LPS did not impact mitochondrial DNA (mtDNA), or cause organelle damage as measured by DNA lesion frequency (Supplementary Figure 9 E,F). Thus, mitochondria are still present in LPS-treated cells, although β-oxidation of FAs is largely uncoupled from OxPhos.

### Blocking CPT1 dampens cytokine/chemokine generation by peritoneal macrophages

CPT1 inhibition doubled the levels of PGs generated by inflammatory activated peritoneal macrophages, in particular the abundant prostacyclin (PGI_2_) metabolite 6-keto-PGF_1_ alpha and PGE_2_ (Figure 1 C). Thus, we next sought to examine whether this impacted PG-mediated autocrine signaling. Several studies have reported that oxylipins such as monohydroxyFAs, PGE_2_ or iloprost (PGI_2_ analog) can suppress cytokine and chemokine generation in monocytes or macrophages, including by EP receptor signaling, or PPARγ activation [42–46], thus we tested the impact of etomoxir on generation of TNF alpha or RANTES in LPS-stimulated peritoneal macrophages. For both proteins, there was a significant decrease in generation when etomoxir was present (Figure 5 E,F). This supports the idea that removing oxylipins can dampen their signaling in macrophages.

### Oxylipin levels detected in vivo regulate leukocyte responses in a concentration dependent manner

Oxylipins are primarily secreted to act on other cell types in a paracrine manner. Next, to address whether macrophage mitochondrial β-oxidation impacts wider oxylipin signaling between other cell types, human phagocyte and T cell responses to these lipids were determined. Neutrophils were chosen since they are known to be sensitive to PGE_2_, via EP receptor signaling [47, 48], while T cells can be regulated by PPARγ, a transcription factor that is regulated by many LOX derived oxylipins [49–55]. Here, reactive oxygen species (ROS) production elicited by *S. epidermidis*, a common pathogen in human peritonitis was measured by aminophenyl fluorescein (APF) fluorescence of whole blood leukocytes [56]. Two concentrations were used that represented amounts of 52 individual oxylipins detected *in vivo,* in the presence or absence of etomoxir (with LPS) (Supplementary Table 1). Here, ROS generation by neutrophils was consistently enhanced by higher or lower oxylipin levels. However, there was a significant concentration-dependent impact in three individual donors (Figure 5 G,H). Thus, the higher amount, consistent with levels detected during CPT1 inhibition, was significantly more potent at activating neutrophils. Monocyte ROS generation was unaffected at either dose (Figure 5 G,H). Next, T cell responses in PBMC were determined by measuring TNF generation following TCR-specific stimulation of CD4+ and CD8+ T cell populations. Here, we found that in general oxylipins had a statistically significant suppressive effect on TNF production in both subsets. Post-test analysis revealed that the lower level of oxylipins alone significantly suppressed TNF in CD4+ cells. In the case of CD8+ cells, only the high level showed significance, while there was a trend at both doses (Figure 5 I). Taken together, these pilot data indicate that modulation of oxylipin levels, within physiological amounts detected in vivo can regulate immune responses of neutrophils and T cells *in vitro*, supporting the concept that oxylipin removal by mitochondria during inflammation could regulate broader cellular innate immune responses.

### Transcriptional analysis of human neonatal bacterial sepsis identifies gene candidates responsible for mitochondrial oxylipin metabolism

The enzymes that import oxylipins into mitochondria are not conclusively known, but it is very likely that those importing long chain FA are involved. These include five acyl-CoA synthetase long-chain family members (*ACSL1,3-6*), as well as *CPT1a, 1b, 1c*, and *CPT2.* The gene products mediate formation of -CoA and then -carnitine derivatives that are required for long chain FA uptake across mitochondrial membranes (Supplementary Figure 10, Supplementary Table 2). This is followed by mitochondrial β-oxidation, which catalyzes sequential removal of 2 carbon fragments to generate acetyl-CoA, and chain shortened metabolites. Genes that encode proteins which metabolize long chain PUFA include *HADHA, HADHB, ECI1, DECR1 and ACADVL* (Supplementary Figure 15, Supplementary Table 2) [22]. A set of 33 genes were compiled including also isoforms with preference for medium or shorter chain FA. Their expression during human infection *in vivo* was tested using a microarray from human neonatal whole blood cells (35 cases, 26 controls). Here, infants suspected of infection had blood cultures conducted, and a diagnosis of bacterial sepsis confirmed [57]. Out of the genes tested, 22 were significantly different with 9 up- and 13 down-regulated in sepsis. Notably, several upregulated genes encode proteins involved in generation of long chain FA-CoAs and their interconversion to acyl-carnitines in mitochondria (Figures 5 J, 6 A). For the most relevant, 7 were significantly upregulated (*ACSL1,3,4, CPT1b, CPT2, ACADVL, HADHB*) while 3 were downregulated (*ACSL5, ECI1, HADHA*) (Figures 5 J, 6 A). This indicates a high degree of regulation during complex bacterial infection in seriously ill humans, suggesting the specific isoforms that maybe involved in oxylipin removal. We also interrogated transcriptomic data from stromal tissues extracted from mice challenged (i.p.) with a cell-free supernatant from a clinical isolate of *S. epidermidis* (SES) [58]. Here, there was overall upregulation, showing a similar trend to the human bacterial dataset (Figure 6 B,C). Although most individual genes were not statistically significant, *Cpt1a* showed around 2-fold induction, which was significant 6 hrs post infection.

**Figure 6.**
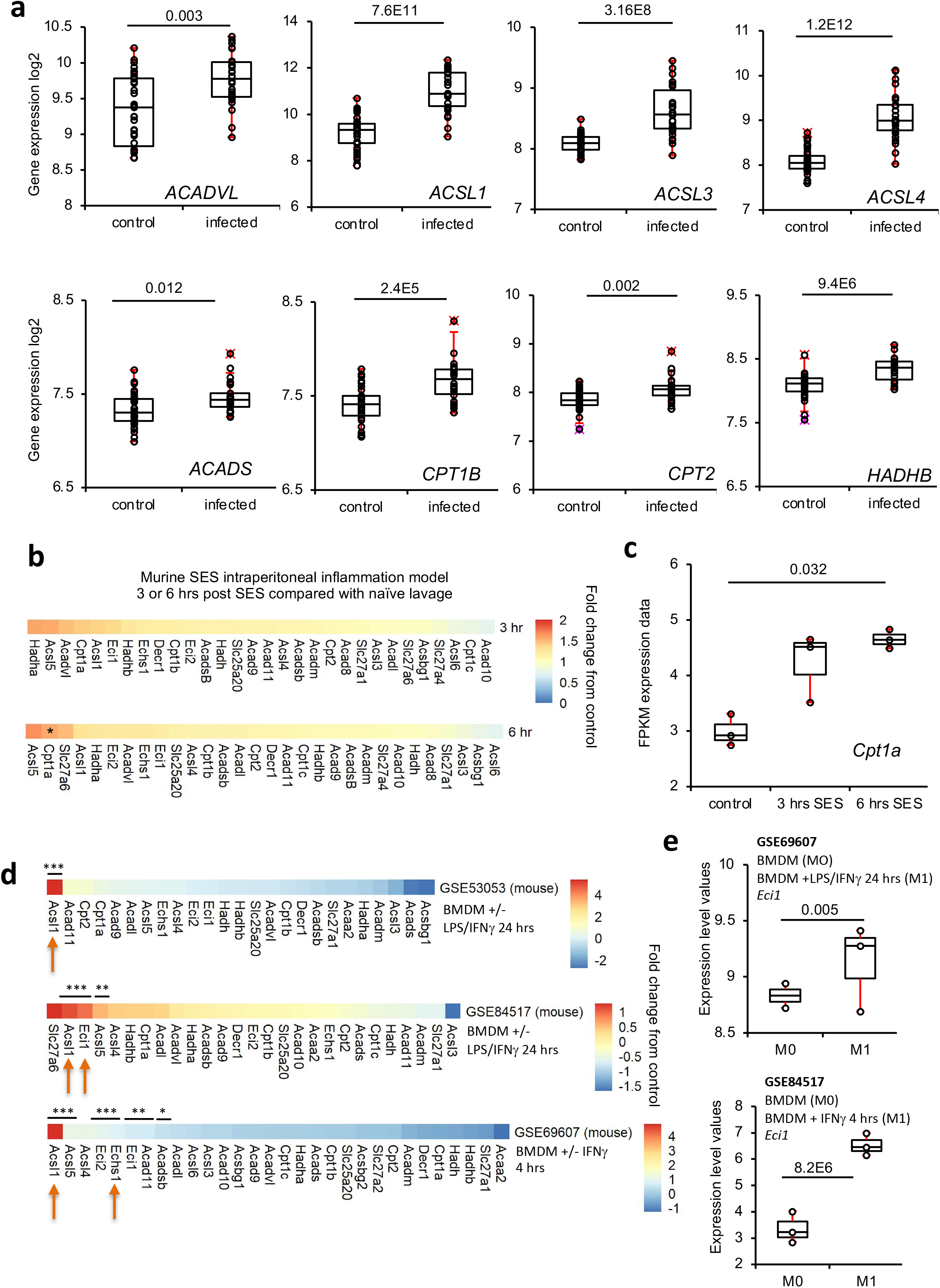
Transcriptomics of the mitochondrial β-oxidation pathway reveals *ACSL1/Acsl1* as a key checkpoint response to LPS inflammation in human and mouse macrophages. *Panel A. Plots for the 8 significantly upregulated genes in the human neonatal dataset are shown.* n = 35 and 26 for infection and controls, respectively. Students T-test, two-tailed, then adjusted using Benjamini-Hochberg test. *Panels B,C. Transcriptomics of mouse peritonitis shows significant upregulation of Cpt1a at 6 hrs post SES*. Gene expression data from peritoneal membranes harvested post SES challenge was analyzed for expression of 32 genes (n = 3 per group) * p < 0.05, Student’s T-test, two-tailed, adjusted using Benjamini-Hochberg test. *Panel C* shows plot for *Cpt1a* expression. *Panels D,E. Transcriptomics reveals Ascl1 and Eci1 to be highly upregulated in response to LPS/IFN*γ *in murine BMDM.* Transcriptomics data obtained from GEO database was analyzed as outlined in Methods for expression of 34 genes selected for potential or known involvement in mitochondrial β-oxidation. Samples were BMDM treated with either LPS/IFNγ or IFNγ alone as indicated. For all genes, the log2fold change was calculated and plotted using Pheatmap in R (Panel D). *Panel E* shows box and whisker plots for normalized expression (using Limma and Oligo Bioconductor packages) of *Eci1* in mouse datasets with n = 3 for all groups, adjusted using Benjamini-Hochberg test (n = 3 for all groups except for GSE53053 M0 (n = 2)). Box shows median, and interquartile ranges (IQR). The ends of the whisker are at 1.5*IQR above the third quartile (Q3) and 1.5*IQR below the first quartile (Q1). If minimum or maximum values are outside this range, then they are shown as outliers.

### A consistent upregulation of ACSL isoforms is revealed across multiple human and murine macrophage datasets, and inhibition of ACSLs dampens 12-HETE metabolism by β-oxidation in RAW cells

Next, data from 3 mouse BMDM or 3 human PBMCs studies were downloaded from GEO [59–64]. Several isoforms of ACSL were found to be upregulated, including *ACSL1,3,4* (human study) or *Acsl1,5* (SES peritonitis and in vitro BMDM experiments). Notably, *ACSL1* expression was upregulated in all studies (Figure 6 D,E, 7 A,B). For BMDM, *Acsl1* was consistently significantly induced on stimulation using either LPS/IFNγ or IFNγ alone (Figure 6D, 7A). Two of the mouse BMDM datasets also indicated significant increases in *Eci1*, encoding 3,2-enoyl-CoA isomerase (Supplementary Figure 15, Figure 6 D,E). In human studies, *ACSL1* was also induced by LPS/IFNγ or IFNγ (Figure 7 B). To confirm the functional relevance of ACSL upregulation, the impact of the pharmacological inhibitor Triascin C on 12-HETE metabolism was tested. 12-HETE removal was strongly inhibited while formation of tetranors was dampened between 60-95 % (Figure 7 C). This is consistent with the requirement for CoA-esters for 12-HETE β-oxidation (mitochondrial or peroxisomal), and confirms the involvement of ACSLs. We note that 12-HETE uptake into phospholipid pools via Land’s cycle esterification also requires ACSL activity. Last, we saw a small non-significant suppression of 12-HETE removal by LPS in RAW cells (Figure 7 C). This is consistent with the process being mainly via Land’s cycle esterification, which was in turn slightly suppressed by LPS (Figure 3 A-C).

**Figure 7.**
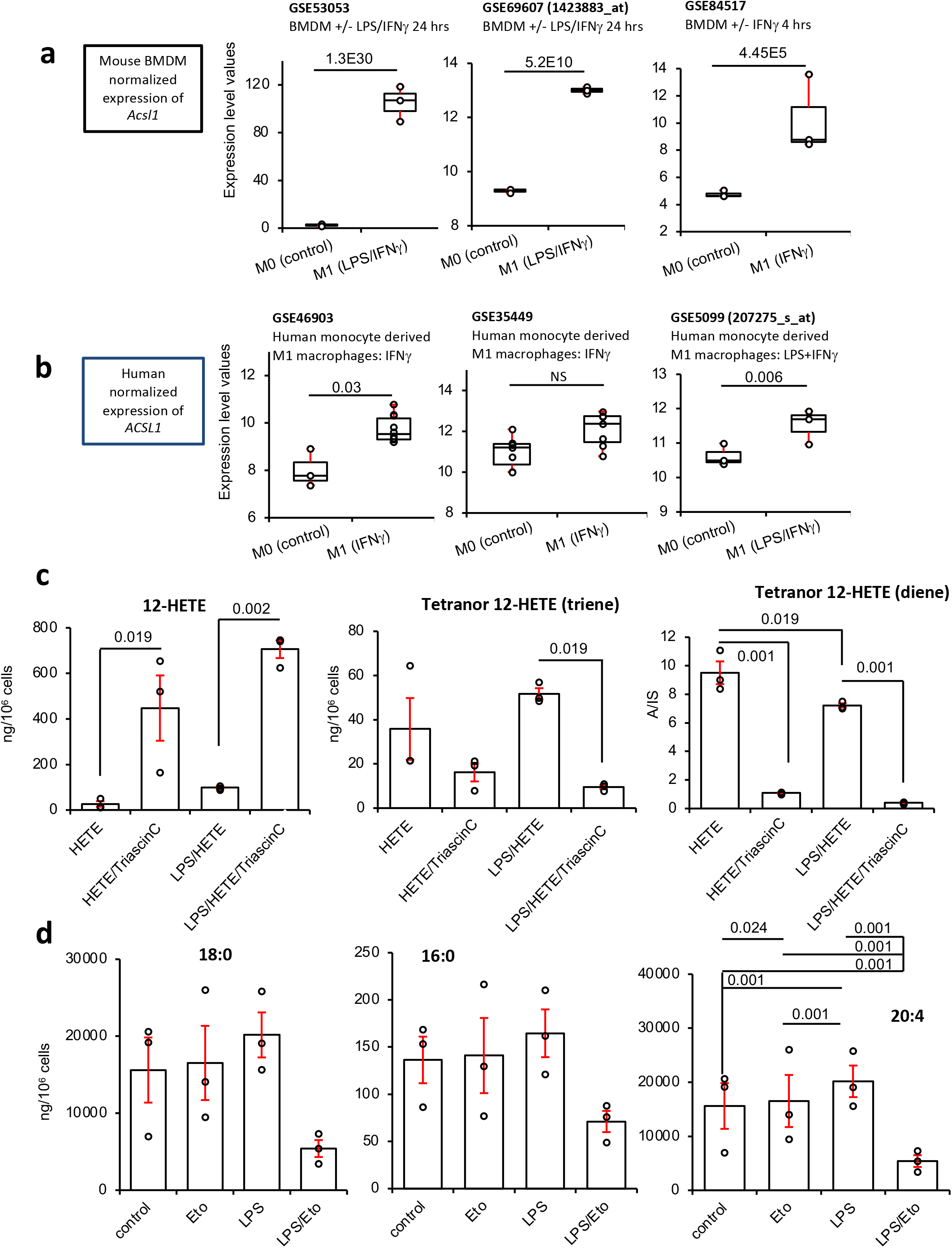
Elevated Acsl1/ACSL1 in mouse and human GEO datasets, triascin C significantly inhibits 12-HETE removal by cells, and prevents generation of tetranor diene or triene HETE metabolites, while etomoxir alters AA levels in RAW cells. *Panel A* Plots for normalized expression (using Limma and Oligo Bioconductor packages) of *Acsl1* in the mouse datasets, with n = 3 for all groups except for GSE53053 M0 (n = 2) adjusted using Benjamini-Hochberg test. *Panel B. ACSL1 is strongly induced in human M1 macrophages.* Human transcriptomics data for *ACSL1* expression was downloaded from GEO and normalized expression level plotted. GSE46903 (n = 3, 10 for M0, M1 respectively), GSE35499 (n = 7), GSE5099 (n = 3). Data were normalized using Limma and Oligo Bioconductor packages as outlined in Methods, then adjusted using Benhamini-Hochberg test. Box shows median, and interquartile ranges (IQR). The ends of the whisker are at 1.5*IQR above the third quartile (Q3) and 1.5*IQR below the first quartile (Q1). If minimum or maximum values are outside this range, then they are shown as outliers. *Panel C. Triascin C alters metabolism of 12-HETE.* RAW cells were cultured for 3 hrs in serum/phenol red free medium with 12(S)-HETE (1.4 g/10^6^ cells), with/without 100 ng/ml LPS with/without 7 μM Triascin C. Cells and supernatant were harvested and 12-HETE and its tetranor metabolites measured using LC/MS/MS (n = 3, mean +/- SEM, separate wells of cells). Significance was tested using ANOVA with Tukey post hoc test. The impact of LPS (either with or without Triascin C) was not significant for any conditions or lipids, except for diene. *Panel D. Etomoxir modulates levels of AA in RAW cells*. RAW cells were cultured for 3 hrs in serum/phenol red free medium with 12(S)-HETE (1.4 μg/10^6^ cells), with/without 100 ng/ml LPS with/without 25 M etomoxir. Cells and supernatant were harvested and 16:0, 18:0 and 20:4 measured using LC/MS/MS (n = 3, mean +/- SEM, separate wells of cells). Significance was tested using ANOVA with Tukey post hoc test.. For 16:0 and 18:0, there were no significant differences found.

### Arachidonate (20:4) levels increase in response to inflammation, and are dampened by CPT1 activity

Pharmacological or genetic inhibition of CPT1/*Cpt1a* dampened PG generation in RAW cells (Figure 3D, 4A, Supplementary Figure 4C). As a potential explanation, we considered whether LPS driven inflammation leads to elevated PUFA biosynthesis, which is then dampened by CPT1 inhibition. Specifically, β-oxidation generates acetyl-CoA, which is converted to malonyl-CoA, used to chain elongate and desaturate essential FA to form PUFA via the gene products of *ELOVL* and *FADS*. To examine this, the impact of LPS/etomoxir on saturated FA (e.g. 16:0, 18:0) and PUFA (e.g. 20:4) levels was tested. No significant effect was seen for saturated FA, although etomoxir tended to decrease them following LPS treatment (Figure 7 D). In contrast, LPS stimulated significant increases in 20:4 levels. Etomoxir had opposing effects on this, dampening AA in resting cells, while enhancing its levels during LPS stimulation (Figure 7 D). Here, LPS most likely increases 20:4 by promoting its phospholipaseA_2_ (PLA_2_)-dependent release from PL membranes. Some 20:4 is then dynamically removed via CPT1, and thus is sensitive to etomoxir. In contrast, the negative impact of etomoxir in resting cells may relate to reduction in supply of malonyl-CoA for elongation. Importantly, the impact of etomoxir on 20:4 was only seen during LPS stimulation, consistent with upregulation of ACSLs and accelerated oxylipin consumption we observed during inflammation. Last, the impact of inflammatory activation on *FADS* or *ELOVL* expression was examined, using the same GEO datasets tested above. However, no consistent trends were seen for any isoforms (data not shown). This indicates that the elongation pathway is not upregulated at the gene expression level by inflammation. In summary, elevation of 20:4 following inflammatory activation most likely arises from activities of phospholipases, and CPT1 then plays a role in its subsequent removal. However, we were unable to elucidate the reason for suppression of PG biosynthesis by inhibition of CPT1, and further studies are required.

## Discussion

Oxylipins are soluble mediators generated during inflammation and infection that link lipid metabolism with immunity. Dynamically controlling their removal is essential to dampen inappropriate inflammation. Here we show that blocking the mitochondrial enzyme carnitine palmitoyltransferase1 (CPT1), effectively doubles secreted levels of endogenously-generated oxylipins *in vivo* during inflammatory challenge. Using exogenous 12-HETE, we propose a paradigm in which macrophage removal of oxylipins and their metabolites involves cycles of secretion and re-internalization of partially-oxidized tetranor intermediates, followed by further mitochondrial metabolism. Extending previous studies, we found high levels of secreted free acid tetranors *in vitro* and *in vivo*, thus regardless of whether formed in peroxisomes or mitochondria, these lipids appear to undergo repeated cycles of acylation/de-acylation (to CoA and carnitine forms, via ACSLs and CPT) facilitating their rapid secretion, re-internalization and further β-oxidation[34, 65]. This demonstrates a complex cellular metabolism for tetranors, and raises questions as to why they are secreted, instead of being simply metabolized to terminal end products. Tetranor compounds can only be formed via β-oxidation (either peroxisomal or mitochondrial). For this to occur, the lipids must be taken up by the cells. Transport of oxylipins into cells is long known to be mediated by a combination of diffusion, and in the case of prostaglandins, prostaglandin transporters (PGT) such as the protein encoded by SLCO2A1 [66]. These proteins have for decades been considered important for supporting cellular inactivation of individual prostaglandin species, and their role in tetranor metabolism could be explored.

Our finding that oxylipin metabolism is strongly stimulated by LPS has implications for the rare genetic disorder, *CPT1 deficiency*, where symptomatic episodes and death are triggered by acute infection[67]. Also, rates of hospitalization and deaths in infants with the *CPT1a* variant p.P479L are higher for homozygotes with respiratory, dental and aural infections[68, 69]. This variant reduces CPT1a activity around 50% [68, 69]. The observation that infection causes severe symptoms indicates that mitochondrial β-oxidation is a protective response to pathogen challenge. This is in line with our findings that CPT1-dependent oxylipin removal is elevated by the bacterial product LPS, and suggests that this maybe an important part of the physiological response to infection, functioning as an “off-switch” for pathologic signaling. Whether oxylipin levels are elevated in patients with CPT1a deficiency is not known, and future studies to examine this are warranted.

Mitochondrial uptake of short/medium chain native FAs is independent of CPT1, however long chain native FA require first activation to acyl-CoA derivatives by acyl-CoA synthetases (ACSL), then conversion to acyl-carnitines, *via* CPT1 (Supplementary Figure 10). Following import into mitochondria via an inner membrane transporter, FA are re-converted to acyl-CoAs by CPT2. We note that so far, evidence for CPT1 being able to import oxylipins relies on the use of etomoxir and *Cpt1a* silencing. Due to the lack of availability of oxylipin-CoAs we are as yet unable to directly test for enzyme activity with isolated mitochondria. Acyl-CoAs are used for mitochondrial β-oxidation, with sequential removal of two C fragments, and generation of acetyl-CoA. Using a human sepsis cohort and multiple human and murine *in vitro* models, we propose a biochemical pathway for oxylipin uptake and mitochondrial metabolism during infection (Supplementary Figure 15). A consistent pattern across all studies was upregulation of *ACSL1/Acsl1*, while we also saw upregulation of *ACSL3,4* (human) and *Acsl5* (mice) (Figure 5J,6A-E,7A,B). Consistent with this, the pan-ACSL inhibitor, Triascin C was highly effective at preventing 12-HETE metabolism to the diene and triene tetranor metabolites in RAW cells (Figure 7C). ACSL1 is already known to shuttle FAs to mitochondria for β-oxidation and lipid synthetic pathways although others may also be involved [70–76]. As yet, the specific ACSLs that acylate oxylipins are not known, although Klett found that all 6 isoforms are active *in vitro* [77]. Given our findings, characterizing which specific ACSLs support the import of oxylipins into mitochondria warrants study. ACSL1 is known to be critical for shuttling FA into mitochondria for β-oxidation in heart, skeletal muscle and adipose tissues [70–74]. Furthermore, in liver, ACSL1 and CPT1 were shown to be physically associated [75, 76]. Thus, ACSL/CPT1 may be a critical checkpoint enabling higher rates of oxylipin metabolism during infection. Indeed, a critical role for ACSL1 in sepsis outcome has been proposed [78]. Regulating import of long chain FAs into the mitochondrial matrix, we found increased *CPT1a or b*, dependent on the model (Figure 6 A-C), along with *CPT2* induction in human sepsis (Figure 6 A). Murine BMDM stimulated with IFN also upregulated *Eci1* (Figure 6 D,E). This encodes DCI, which converts 3-cis or trans-enoyl-CoA to 2-trans-enoyl-CoA during mitochondrial β-oxidation and was previously shown to be induced during Hepatitis C virus infection and required for virus replication [79, 80]. Last, upregulation of the β subunit of the mitochondrial trifunctional protein (encoded by *HADHB*) and very long chain acyl-CoA dehydrogenase (encoded by *ACADVL*) are also fully consistent with their proposed role in oxylipin β-oxidation (Figure 6 A, Supplementary Figure 15).

Macrophages rely primarily on glycolysis for energy when stimulated with LPS/IFN and conversely use FAO to supply OxPhos with substrates, when exposed to immune-modulatory stimuli such as IL-4 [26]. However, recent studies have uncovered a significant role for lipid metabolism in “M1” cells even in the absence of OxPhos, with FA synthetic pathways (FAS) being upregulated by LPS [26]. Our observation that etomoxir partially suppressed 18:0 and 16:0 levels in LPS-stimulated RAW cells is consistent with this (Figure 7 D). This anabolic event is proposed to supply inflammatory activated cells with lipids for processes such as proliferation (sphingolipids, glycerolipids, phospholipids, etc), and generation of oxylipins [26]. However, in the case of PUFA substrates for oxylipin synthesis, fatty acid elongation/desaturation of the essential fatty acids, linoleic and a-linolenic acid by *FADS* and *ELOVL* gene products is the key event required. Although we found significant elevations in 20:4 (Figure 7 D), this was most likely mediated via PLA_2_ activities since there was no impact of etomoxir and *FADS* and *ELOVL* genes were not upregulated by LPS. Here, we focused down on inflammatory activated macrophages that generate oxylipins, where metabolism will favor glycolysis and FAS, and reduced OxPhos. We show for the first time that CPT1 mediates exogenous and endogenously-generated oxylipin β-oxidation by macrophages, that is elevated during inflammation. The impact of blocking CPT1 varied by sub-family of oxylipins, based on either:

*(i) Cell type*. Etomoxir consistently elevated LOX-derived oxylipins and various other mono-hydroxy-oxylipins, while suppressing CYP/sEH metabolites in all macrophages. However, COX-derived PGs increased *in vivo* and in primary macrophages, but were suppressed in RAW cells.

*(ii) How CPT1 was targeted.* In RAW cells, etomoxir or gene silencing suppressed COX-derived PGs and CYP/sEH oxylipins. In contrast, monohydroxy oxylipins were increased by etomoxir but suppressed by CPT1 knockdown in RAW cells. Etomoxir was used at 25 M, well below concentrations found to induce off target effects on adenine nucleotide translocase [36, 37]. Indeed, previous studies using high levels of etomoxir led to FAO being incorrectly proposed as required for alternative activation of M2 macrophages (100-200 μM) [81]. Nowadays, lower concentrations are recommended that block ∼90% of β-oxidation without side-effects (e.g. 10 - 25 μM) [38]. As a second approach, shRNA knockdown stably reduced expression of CPT1a. However, this causes adaptive changes to the metabolic status of macrophage mitochondria, beyond acute inhibition of CPT1. Recent publications implicate CPT1 in maintaining mitochondrial health due to its requirement for importing FAs for maintaining mitochondrial cardiolipin and phospholipid pools [38]. Thus, cells lacking CPT1 do not proliferate normally and may have adapted to a deficiency in OxPhos through alterations in other lipid and energy metabolism pathways. Thus, constitutive knockdown may directly impact the balance between mitochondrial and peroxisomal β-oxidation and the ability to support elongase activities in the cells. Nevertheless, both inhibitory approaches caused significant changes to oxylipin secretion, both for PGs, 12-HETE and its two tetranor metabolites, supporting the hypothesis that mitochondria can be a significant site of oxylipin regulation in macrophages.

### Our data reveal complex modulation of oxylipin removal and formation by CPT1 as follows

*(i) Removal:* Oxylipins and their metabolites can be degraded by two separate β-oxidation pathways, with only mitochondrial enzymes relying on CPT1. However, peroxisomes and mitochondria display metabolic interplay where FA degradation intermediates are transferred for metabolism between the organelles [82]. Thus, stable knockdown of CPT1 may require compensatory increases in peroxisomal β-oxidation. It has been proposed that truncated unsaturated FA metabolites may need to transfer to mitochondria for complete metabolism [82]. Indeed, 12-HETE-triene was suggested as the terminal peroxisomal metabolite for 12-HETE, with further metabolism proposed to require mitochondria [18]. Our data provide supportive evidence for this interplay between organelles, since blockade of CPT1 had a far greater impact on metabolism of tetranor metabolites than exogenous 12-HETE in LPS stimulated cells. Thus, 12-HETE can be metabolized in either organelle, but peroxisomal-generated tetranors are then shuttled to mitochondria, with their oxidation significantly stimulated by inflammation.

*(ii) Generation*: Oxylipins are generated from PUFA, which are themselves synthesized via chain elongation and desaturation of linoleic and α-linolenic acids. This process requires malonyl-CoA, which is formed from acetyl-CoA, a product of which themselves depend at least in part on CPT1 for formation. Thus, while both synthesis and degradation of oxylipins simultaneously occur in macrophages, steady state levels will depend on the cells’ metabolic status, and whether CPT1 is mainly supporting degradation (by β-oxidation) or synthesis (by supplying TCA intermediates) of individual oxylipins. Thus, CPT1 could either increase or decrease levels of oxylipins and their metabolites concurrently, as seen in our study. A detailed flux analysis of this phenomenon is required to further our understanding, since a role for regulation of elongation/desaturation was not clearly seen in our study.

Here, we show that mitochondrial oxylipin removal takes place during inflammation on a background of reduced OxPhos, a well-known response to LPS stimulation by macrophages [28]. However, β-oxidation requires a basal level of OxPhos both to regenerate NAD+ and oxidized flavin cofactors in the electron transferring flavoprotein complex, that reduces ubiquinone. While OxPhos was suppressed, there was a low residual activity remaining in both RAW and peritoneal cells that appeared sufficient to sustain β-oxidation, observed by formation/metabolism of oxylipins and HETE tetranors. Our experiments using exogenous 12-HETE suggest that non-mitochondrial pathways such as peroxisomal oxidation and also esterification is probably responsible for removing most. This agrees with previous studies where exogenous 12-HETE added to mammalian cells was removed by peroxisomes forming a series of metabolites, down to C12:1 [18, 34, 83, 84], and a significant amount is converted to esterified pools[35]. However, these studies did not consider the impact of inflammation and the focus was exclusively on exogenous 12-HETE, not endogenous oxylipins generated by the cells. Here, we extend this a range of oxylipins generated by macrophages, and reveal that during LPS-inflammation, mitochondria (via CPT1) effectively halve secreted amounts of oxylipins en masse, and are a key site for removing peroxisomal metabolites of 12-HETE. We demonstrate that this alters the outcome of leukocyte activation, suggesting that mitochondrial β-oxidation may act as a brake for dampening host inflammatory responses *in vivo* during infection. Further studies are required to test this idea *in vivo*, including characterization of the ability of other leukocytes such as neutrophils and T cells to remove oxylipins via β-oxidation The potential implications for humans with CPT1a deficiency, relating to severely augmented inflammation during infection should be considered and studies initiated to test this idea. Also, whether this phenomenon contributes to infection-associated host inflammation in general, in cases where mitochondrial health is compromised, should be considered.

## Methods

### Mouse peritonitis model

All mouse experiments were performed in accordance with the United Kingdom Home Office Animals (Scientific Procedures) Act of 1986 (P05K6A456) and for peritoneal models under (UK Home Office PPL 30/2938, and PPL P05D6A456). Wild-type female C57BL/6 (strain code 632) mice were purchased for all studies from Charles River Laboratories. Mouse breeders were housed in isolators (barrier) and all experimental mice in individually ventilated cages with 12-h light/dark cycles and controlled temperature (20 – 22 °C) (specific pathogen-free/SPF). Access to water and standard chow was ad libitum, and humidity was atmospheric, and not specifically regulated. Female C57BL/6 mice wild type mice (7-9 weeks) were injected intraperitoneally with etomoxir (100 µg, Sigma Aldrich, Cat. No E1905) and/or LPS (1 µg, E.Coli 0111.B4, Sigma-Aldrich, Cat. No L3024) or vehicle (PBS) per mouse. After 6 hr, mice were euthanized using CO_2_, then peritoneally lavaged using 5 ml ice cold PBS (with 5 mM EDTA). Eicosanoid internal standard mix was added to the lavage solution (7.5 μl per 5 ml lavage solution, Supplementary Data File 2). Cells were separated by centrifugation and resuspended in 1 ml PBS. Supernatant and cells were frozen at −80°C until lipid extraction using solid phase as described below.

### Isolation and treatment of resident peritoneal macrophages

Wild type mice (female, 9-13 weeks) were lavaged with PBS supplemented with 5 mM EDTA, and peritoneal cells from 4 - 5 mice pooled together. A small amount was labelled with F4/80 (BD Horizon, BV421 Rat Anti-Mouse F4/80, Clone T45-2342, Cat.: 565411) and CD11b-allophycocyanin antibodies (BD Pharmingen, Clone M1/70, PE Rat Anti-Mouse, Cat 557397), both 0.2 µg/ml final concentration, to determine macrophage numbers. Cells were first blocked for non-specific antigen binding (incubating for 20 min at RT with 2 µg/ml CD16/CD32 Rat anti-Mouse, unlabeled, Clone: 2.4G2, BD, 553142) then antibodies were added without washing off the blocking solution, and cells were incubated for further 20 min at RT. Cells were then centrifuged for 5 min at 350g, 4C, supernatant discarded and cells resuspended in 150 µl PBS and % macrophages determined using flow cytometry. Peritoneal cells were seeded at 10^6^ macrophages/well of a 6-well plate and cultured in DMEM (with 10 % heat inactivated FCS and pen/strep), for 2 hours at 37 °C, 5 % CO_2,_ 99 % humidity. Nonadherent cells were removed by washing with media, and the remaining cells cultured in serum free, phenol red free, RPMI, with/without LPS (100 ng/ml) and/or etomoxir (25 μM). After 24 h, cells and supernatants were harvested separately, with supernatant first centrifuged (350g, 5 min, 4 °C), and cells scraped quickly into 1 ml RPMI. Supernatant and cells were then snap-frozen in liquid nitrogen and stored at −80 °C, prior to lipid extraction and analysis. For analysis of TNFα and CCL5/RANTES, supernatants were analysed using commercial ELISA assay kits (R & D Systems; Mouse TNFα DuoSet ELISA, Cat. No DY410-05; Mouse CCL5/RANTES DuoSet ELISA, Cat. No DY478-05) according to the manufacturer’s protocol.

### Generation and culture of RAW264.7 cell lines stably overexpressing Alox15, or expressing shRNA targeting Cpt1a

*Alox15* overexpressing RAW264.7 cells were generated by retroviral transduction using the pMX-IP retroviral vector, which is based on the Moloney Murine Leukemia Virus (MMLV) [85]. Briefly, mRNA was extracted from C57BL/6 peritoneal cells, and *Alox15* was PCR amplified with the following primer sequences:

- Alox15-Fw TCCTGCAGGCCTCGAGCCACC<u>ATG</u>GGTGTCTACCGCATCC
- Alox15-Rv: CGCGCCGGCCCTCGAGTCATATGGCCACGCTGTTTT

The In-fusion cloning kit (Takara Bio, 102518) was utilized to insert *Alox15* into the *Xho I* site of the pMXs MMLV retroviral plasmid. Retroviral particles were generated after transfection of HEK293T cells with the viral plasmid. RAW264.7 cells were spin-infected with packaged retrovirus (*Alox15* or empty vector control pMXs-IP). Positive infections were selected by resistance to 3 μM puromycin, screened for positive *Alox15* expression by PCR and 12-HETE generation using LC/MS/MS (Supplementary Figure 11 A). Separately, RAW264.7 cells were infected with lentiviral vectors to generate *Cpt1a* knockdown or non-silencing control cell lines, as described [86]. Alterations from the published method were different lentiviral plasmids (pSMART mEF1a/TurboGFP (Cpt1a targeting shRNA [V3SVMM03-13316872] and non-targeting control) - Dharmacon). Briefly, lentiviral particles were created by transfecting HEK293T cells (lentiviral plasmid plus helper constructs (pCMV-Δ8.91 (Gag/Pol, Tat and Rev) and pMD2.G (VSV-G coat)) using the Effectene transfection reagent (Qiagen). Lentivirus was concentrated 48 h after transfection from 0.45 µm filtered supernatants layered on top of 1/6^th^ volume of 0.584 M sucrose using a Beckman Coulter Ultracentrifuge (100,000 x g, 90 min, 4 °C). Lentiviral particles concentrated from 20 ml of supernatant were added to a ∼10 % confluent RAW264.7 cells in a T-175 flask for 3 days. After another 4 days of subsequent cell expansion positively infected cells were sorted using GFP expression on a BD FACS Aria cell sorter (100 μm nozzle size) (Supplementary Figure 9 C), using Attune NxT software (V2). Knockdown was validated by real-time PCR. Mature *Cpt1a* mRNA target sequence: CAGTGGTATTTGAAGCTAA, the sequence of the non-targeting control was not disclosed by Dharmacon but is validated to have minimal targeting to known mouse genes. To confirm knockdown, RNA was extracted using the PureLink RNA extraction kit, cDNA was created using the high capacity reverse transcription kit (Life Technologies). Real-time PCR was performed using the Taqman assay (Applied Biosystems) on a Stratagene MX3000P real-time PCR instrument. Data was analyzed using the ΔΔCt method, normalizing Cpt1a (probe spans exons 13-14: Mm01231183_m1) gene expression to the housekeeping genes Hprt (probe spans exons 2-3: Mm03024075_m1) and Actb (probe spans boundary of exon 3: Mm02619580_g1) (Supplementary Figure 9 A). RAW cells were cultured in DMEM with 10 % FCS and pen/strep. Cells were passaged at ∼80 % confluency using PBS with 5 mM EDTA and 27.7 mM lidocaine for ∼5 min at 37 ^0^C. Cells were cultured with/without 100 ng/ml LPS, with/without 25 μM etomoxir for 24 h in serum free RPMI medium. Then cells and supernatants were harvested separately and stored at −80^0^C until lipid extraction and analysis using LC/MS/MS.

#### Bone marrow derived macrophages (BMDM) isolation and differentiation

Bone marrow cells were harvested from femurs and tibias of mice (8-10 weeks old, female C57BL/6) using syringe flushing. After centrifugation and washing twice with PBS cells were plated at 2 x 10^5^/ml in DMEM (10% FCS, +/- 5 % horse serum, pen/strep, L-glutamine, +/- HEPES). MCSF was added at final concentration 20 ng/ml. On day 3, cells were refed using media and MCSF. On day 7-8, cells were washed with PBS, then lifted with lidocaine/EDTA or ACCUTASE™ (Sigma-Aldrich), plated at 1 - 2 x 10^6^ per well of a 6-well plate and rested for 24 h. Cells were washed and then refed with DMEM (as above, with MCSF) and cytokines as follows: M0: medium only, M1: LPS (100 ng/ml)/IFNγ (20 ng/ml), M2: IL-4 (20 ng/ml). For some experiments, FCS was omitted. Etomoxir (25 µM) was added in some experiments. After 24 hrs supernatant was recovered and frozen at −80 °C. Cells were washed and then recovered in 1 mL of PBS using rapid scraping on ice, and the cell suspension immediately quenched in liquid N_2_ prior to storage at −80 °C.

Samples were analyzed for phenotype confirmation (cytokine production, mRNA gene expression by real time PCR) and lipid composition, as described. CCL5, TNFα and IL-6 in supernatants were measured using ELISAs (Mouse TNFα; Mouse CCL5; Mouse IL-6, R&D Systems) according to the manufacturer’s protocols. Confirmation of phenotype is shown in Supplementary Figure 11 B,C. Total RNA was recovered using the RNeasy Mini Kit (QIAGEN, 74104) according to the manufacturer’s instructions. RNA concentration and purity were determined using a Nanodrop spectrophotometer. cDNA was prepared from 1 µg of RNA. RNA was reverse-transcribed using superscript III reverse transcriptase (Invitrogen) in a total volume of 20 µL for 1 h at 50 °C, and the reaction was terminated at 70 °C for 15 min. Power SYBR Green PCR Master Mix (Thermo Fisher scientific) was used for real-time PCR. The Master Mix (10 μL) was added to dH2O (7.5 μL), and 1 μl primer mix (0.5 μg/mL each primer) to make a gene specific master mix; cDNA (1.5 μL) from each sample was added to the reaction mix (17.5 μL) in a 96-well plate well. The plate was inserted into a QuantStudio 3 Real-Time PCR system and the PCR amplification started as per instrument instruction: 50°C – 2 min; 95°C - 10 min; 95°C - 15 s; 60 °C - 1 min; Repeat 3 - 4 for 40 cycles; 95°C - 15 s; 4 °C end. Primers used for real-time PCR were designed to be intron-spanning, and to have a melting temperature 55 - 65 °C. The intron-spanning aspect ensured that only mRNA and not genomic DNA were amplified. All primers and oligonucleotides were obtained (Sigma, Gillingham UK) as salt free purified lyophilized DNA. The sequences of the real time PCR primers used are shown in Supplementary Table 3.

#### Metabolism of exogenous 12-HETE and 12-HETE-d8 by RAW cells

RAW cells (2 x 10^6^/ml media) were supplemented with 9 µM 12(S)-HETE, 12(S)-HETE-d8, 12(R)-HETE tetranor 12(S)-HETE (from Cayman Chemical) for up to 12 hrs in serum free RPMI, with 100 ng/ml LPS, with/without 25 µM etomoxir. Amounts supplemented were (per 10^6^ cells): 12h timecourse 12(S)-HETE: 1.44 μg. 3 hr fixed time supplementation experiments: 12(S)-HETE/12(R)-HETE, 12(S)-HETE d8: 1.5 μg, tetranor 12(S)-HETE triene: 1.2 μg. Following incubation, supernatant was harvested, centrifuged to remove any dead/dying cells and then snap frozen. Cells were gently rinsed with PBS to remove any dead/dying cells. Following this, the cells were scraped in PBS and snap frozen. Samples were stored at −80 °C. Lipids were extracted as described below, using solid phase C18 columns, resuspended in small volumes of methanol and stored at −80 °C until analysis using LC/MS/MS. For experiments testing the impact of Triascin C, cells and supernatant were together scraped and harvested before snap freezing, then processing as outlined below for lipid extraction and analysis.

#### Lipid extraction

Internal standard mixture (see eicosanoid assay for details, Supplementary Data File 2) was added to each sample before extraction. Oxylipins were extracted from cells by adding 2.5 ml of solvent mixture (1M acetic acid/2-isopropanol/hexane (2:20:30, v/v/v) to 1 ml of sample, followed by vortexing for 60 s, and then adding 2.5 ml of hexane. The mixture was vortexed again for 60 s, centrifuged at 500 g, 4 °C for 5 min, and lipids recovered in the upper organic solvent layer. The lipids were reextracted by adding another 2.5 ml of hexane, followed again by vortexing and centrifugation. The lipids were again recovered in the upper organic solvent layer. The combined hexane layers were dried by vacuum and resuspended in methanol. Oxylipins were extracted from supernatants using solid phase extraction (SPE) using C18 columns (Sep-Pak C18, Waters). Columns were activated using 12 ml methanol, followed by 6 ml acidified water (0.4 % glacial acetic acid in HPLC water) followed by loading the sample (all samples were in 85 % aqueous, 15 % methanol, pH 3.0), and washing first with 10 ml acidified water (0.4 % glacial acetic acid in HPLC water), then with 6 ml of hexane. The columns were left to dry for 15 - 20 min followed by eluting the lipids with 8 ml methyl formate. The lipids were vacuum dried, resuspended in methanol and stored at −80 °C.

### LC-MS/MS targeted analysis of oxylipins

Targeted lipidomics analysis of 93 oxylipins was performed on a Nexera liquid chromatography system (Nexera X2, Shimadzu) coupled to a 6500 QTrap mass spectrometer (AB Sciex) as described, with minor adaptations [87]. Briefly, liquid chromatography was performed at 45 °C using a Zorbax Eclipse Plus C18 (Agilent Technologies) reversed phase column (150 × 2.1 mm, 1.8 μm) at a flow rate of 0.5 mL/min over 22.5 min. Mobile phase A was (95 % HPLC water/5 % mobile phase B; v/v and 0.1 % acetic acid) and mobile phase B was acetonitrile/methanol (800 ml + 150 ml; and 0.1 % acetic acid). The following linear gradient for mobile phase B was applied: 30 % for 1 min, 30 - 35 % from 1 to 4 min, 35 – 67.5 % from 4 to 12.5 min, 67.5 – 100 % from 12.5 to 17.5 min and held at 100 % for 3.5 min, followed by 1.5 min at initial condition for column re-equilibration. Injection volume was 5 µL. Lipids were analyzed in scheduled multiple reaction monitoring (sMRM) mode with a detection window of 55 s for each analyte. Ionization was performed using electrospray ionization in the negative ion mode with the following MS parameters: temperature 475 °C, N_2_ gas, GS1 60 psi, GS2 60 psi, curtain gas 35 psi, ESI voltage −4.5 kV. Cycle time was 0.4 s. Peak areas for lipids were integrated and normalized for deuterated internal standards and quantification was performed using external calibration curves with authentic standards for all investigated oxylipins. Data was acquired using Analyst (V1.6) integration and quantification was performed using MultiQuant software (version 3.0.2, AB Sciex Framingham, MA, U.S.A.). A full list of the internal standards and oxylipins is provided in Supplementary Data File 2 along with calibration range, MRM transitions and corresponding MS parameters (collision energy, declustering potential). See Supplementary Data File 1 for list of abbreviations and formal names for lipids measured. In this assay, identification is based on retention time comparison with authentic standards and the presence of a diagnostic precursor and product ion pair, however stereochemistry is not established and in the case of lower abundance lipids, is often unclear. For that reason, chirality has been removed. Limit of quantitation (LOQ) for each lipid was set at signal:noise 5:1, and the primary standard amount (ng on column) closest to this value for each lipid is shown in Supplementary Data File 2, column N. All chromatograms from tissue/cell samples were manually checked for peak quality and excluded from analysis if they fell below limit of 5:1 (signal:noise, LOQ) with a minimum of 6 points across each peak. Where lipids fell below LOQ, a zero value was recorded, but replaced with 50%LOQ for statistical analysis. Example chromatograms for all lipids analyzed are provided in Supplementary Figure 12.

Targeted lipidomics analysis of 12(S)-HETE, 12(R)-HETE, 12(S)-HETE-d8 or tetranor 12(S)-HETE metabolites was performed on a Nexera LC coupled to a 4000 QTrap (AB Sciex). Briefly, liquid chromatography was performed at 40 °C using a Spherisorb ODS2 C18 column (4.6 x 150 mm, 5 µm, Waters) at a flow rate of 1 ml/min over 30 min. Mobile phase A was (75 % HPLC water/ 25 % acetonitrile; v/v and 0.1 % acetic acid) and mobile phase B was 60 % methanol/ 40 % acetonitrile; v/v and 0.1 % acetic acid. The following gradient for mobile phase B was applied: 50-90% B over 20 mins, then held at 90% B for 5 mins, followed by re-equilibration to 50% B from 25-30 mins. Injection volume was 10 µL. Ionization was performed using electrospray ionization in the negative ion mode with the following MS parameters: temperature 650 °C, N_2_ gas, GS1 60 psi, GS2 30 psi, curtain gas 35 psi, ESI voltage −4.5 kV. Cycle time was 1.46 s. Peak areas for lipids were integrated and quantification was performed using external calibration curve with 15-HETE-d8 as standard for all investigated compounds. As no primary standard is available for the tetranor diene, data is expressed as analyte/internal standard signal ratio (A/IS). Data integration and quantification was performed using MultiQuant software (version 3.0.2, AB Sciex Framingham, MA, U.S.A.). MRM transitions and corresponding MS parameters (collision energy and declustering potential) for each analyzed lipid is provided in Supplementary Table 4.

### Mitochondrial stress test

Oxygen consumption rate (OCR) was measured using Seahorse XF^e^96 analyzer (Agilent, with Wave 2.3/4 software) and the Seahorse XFe96 FluxPak (102416-100, Agilent) according to the manufacturer’s protocols. Briefly, RAW/RAW*Alox15* cells were seeded at 2.5 x 10^5^ cells/well and peritoneal macrophages were seeded at 5 x 10^5^ cells/well (and incubated with LPS (100 ng/ml)). Mitochondrial function was assessed using inhibitors (all from Sigma) injected in the following order: oligomycin (1µM), FCCP (Carbonyl cyanide 4-(trifluoromethoxy)phenylhydrazone, 2 µM or 4 µM final concentration for peritoneal macrophages or RAW cells respectively), rotenone (1 µM), antimycin A (10 μM for peritoneal macrophages; 1 µM for RAW/RAW*Alox15* cells). OCR is described as pmol/min (per well).

### Heatmap method

For generation of heatmaps, amounts (ng/10^6^ cells) of each lipid were averaged, then log10 was applied to the mean value. Heatmaps were generated using Pheatmap (V1) in R (3.6.3). The Euclidean metric was chosen to establish the treatments’ relationships depicted as clusters[88]. The clusters are aggregated following the “shortest distance” rule. Lipids are color-coded by putative enzymatic origin (Supplementary Data File 1).

### Determination of reactive oxygen species generation by peripheral blood cells

Human blood from healthy workplace volunteers was obtained with ethical approval from School of Medicine (SMREC 16/02) with informed consent. Volunteers were male or female, healthy, between the ages of 25-55, with no genotype information. There was no participant compensation. Whole blood (5 ml) was drawn using a butterfly syringe into 50 μl heparin sodium; 12μl of this was then placed in a microcentrifuge tube and 1 ml cold PBS added. Following centrifugation (350 x g, 5 min, 4 °C) the cell pellet was recovered and resuspended, then stained with anti-CD15 BV510 (1:50 in 0.5 ml PBS, W6D3, Becton Dickinson, 563141) for 30 min on ice. Cells were centrifuged (350 x g, 5 min, 4 °C) and resuspended in DMEM media (2 mM glutamine) containing amino-phenyl fluorescein (APF) (1:1000, cell concentration = cells from 5 μl blood per ml of APF media). *S. epidermidis* bacteria (clinical isolate) [56] were killed by incubation at 65 °C for 20 min, then washed by centrifugation and resuspended in PBS three times (12,000 x g, 5 min). Bacteria were counted by flow cytometry using an Attune NxT; bacteria were excluded from other particles by size (Forward-scatter vs Side-scatter). Bacteria were first added to blood cells at varying doses, to determine optimal reactivity per donor isolate (ranging from 1.37 x 10^5^ −2.75 x 10^6^ bacteria per ml of APF media containing cells from 5 μl blood). For this, cells+bacteria were incubated for 20 min at 37 °C, then tubes placed on ice. Cells were washed three times with ACK red blood cell lysing buffer (350 x g, 5 min, 4 °C) then analyzed by flow cytometry. Cell gating strategies are shown in Supplementary Figure 11 D-F. This included forward-vs side-scatter profile (FSC vs SSC) that distinguishes neutrophils and monocytes from lymphocytes. Neutrophils were further separated from eosinophils and monocytes via their high expression of CD15. The bacterial dose was selected based on a neutrophil APF^+^ response of between 10 - 40 % APF^+ve^ cells. Once this was optimized, the experiment was repeated in triplicate for each donor, and using three individual donors, using a fresh aliquot of blood. The additional step following APF addition, was that lipids or ethanol vehicle (1:100) were added and samples incubated at 37 °C for 10 min before the bacteria addition step. Lipids used and final concentrations are provided in Supplementary Table 1. Calculations assumed a 100 μl peritoneal volume *in vivo* (Supplementary Table 1) [89]. Next, bacteria were added, cells were incubated for 20 min at 37 °C, then tubes placed on ice, washed, and analyzed using flow cytometry for APF^+ve^ neutrophils or monocytes. Means for each donor were calculated separately and compared using two-way ANOVA with Sidak’s post-test, where p < 0.05 was considered significant.

### T-cell activation assay

All procedures were performed at 4 °C, centrifugation steps were 350 x g for 5 min, and reagents and equipment were from Theromofisher and antibodies from Biolegend unless stated otherwise. Whole blood was acquired with heparin sodium (Sigma - 20 USP units per ml) and gently layered onto 2/3 volume of Histopaque 1077 in a centrifuge tube. Blood was centrifuged at room temperature at 400 x g for 15 min, (low acceleration and deceleration), the peripheral blood mononuclear cell (PBMC) layer and half the serum and histopaque volume were removed to a separate tube. Six times volume of cold PBS was added, and cells centrifuged. Supernatant was removed completely, and cells were resuspended in serum free RPMI media (+ L-glutamine) for counting, RPMI was added to achieve 5 million per mL (centrifugation and resuspension required – acts as an additional wash). PBMCs (50 μL – 250,000 cells) were added to separate wells of a 96-well U-bottom plate and incubated at 37 °C for 20 min. Lipid mixes (25 μL of 4x concentrated high, low or ethanol control in media) were added and incubated for 15 min at 37 °C. Anti-CD3/CD28 Dynabeads (25 μL of 4x concentrated in media = 0.64 beads per PBMC) or control were then added and incubated for 2 h at 37 °C. Golgi Plug and Golgi Stop (BD biosciences, 5 μL of 20x concentrated in media) were added for the last 4 h of stimulation at 37 °C (6 h total). Ice cold wash solution (PBS + 5 mM EDTA + 5% BSA) was added to each well and cells mixed and transferred to a 96-well v-bottom plate. The plate was centrifuged, and supernatant removed. Cells were resuspended in 25 μL block solution (wash plus 1 in 40 Human TruStain FcX – Biolegend) and incubated for 5 min. Surface stain antibody cocktail was added – 25 μL wash containing 0.5 μL Zombie Live/Dead Aqua, 0.25 μL anti-CD8 BV-711 (RPA-T8) (BioLegend, cat # 301044, used at 1:100 dilution), 1 μL anti-CD4 BV605 (OKT4) (BioLegend, cat # 317438, used at 1:25 dilution), 0.5 μL anti-CD3 Percp (SK7) (BioLegend, cat # 344814, used at 1:50 dilution). The plate was incubated at room temperature for 20 min. Wash solution (150 μL) was added and cells centrifuged, the supernatant was removed and cells washed again with wash solution. Cells were resuspended in 100 μL Fix/Perm solution (BD Biosciences) and incubated for 30 min. Wash solution (100 μL) was added and cells centrifuged, the supernatant was removed, and cells washed again with wash solution. Wash solution (150 μL) was added, and cells were stored overnight. The plate was centrifuged, and supernatant removed, cells were resuspended in internal antibody mix – 50 μL perm-wash (BD biosciences) with 1 μL anti-TNF APC-Vio770 (cA2) (Miltenyi Biotec, cat # 130-120-169, used at 1:50 dilution). The plate was incubated for 30 min. Perm-wash solution (150 μL) was added and cells centrifuged, the supernatant was removed and cells washed again twice with perm-wash solution. Cells were resuspended in 0.5 mL wash solution and acquired on a flow cytometer. Anti-mouse IgG-κ compensation particles were used for single stain compensation controls (BD Biosciences, cat # 552843, 1 drop of beads per 100ul staining buffer. 50ul used per compensation control tube). Live, single cells were gated, and debris removed before analysis of CD3 positive events.

### Assessment of mitochondrial DNA in LPS treated cells

RAW cells were seeded in 6 well plates at 1×10^6^ cells/well and grown in DMEM with 10% FCS, 2mM glutamine and 100 U/ml of penicillin/streptomycin for 24 hr. Next, medium was aspirated, and LPS (100 ng/ml) or chloroquine (40 nM) added, and cells incubated for 24 hr. Total DNA was extracted from cells with PureLink Genomic DNA kit (Fisher Scientific, cat. No. 10053293) according to the manufacturer’s instructions and quantified using a Nanodrop (Thermo Scientific). Assessment of mtDNA copy numbers, relative to genomic DNA (gDNA), was done as previously described [90]. Briefly, quantitative PCR (qPCR) analysis was performed using PerfeCta^TM^ SYBR Green FastMix^TM^, low ROX^TM^ (Quanta Biosciences) and primers specific for mouse mtDNA (mouse mtDNA forward: 5’-CCCCAGCCATAACACAGTATCAAAC-3’, mouse mtDNA reverse: 5’-GCCCAAAGAATCAGAACAGATGC-3’) or the genomic 18S ribosome gene (mouse 18S forward: 5’-AAACGGCTACCACATCCAAG-3’, mouse 18S reverse: CAATTACAGGGCCTCGAAAG-3’) (Merck KGaA). Primers specific for mtDNA gives rise to a 201bp product, while primers specific for the 18S ribosomal gene gives a product that is 112 bp. qPCR was performed in a Stratagene MX3000P machine (Stratagene California, San Diego, CA, USA) using 96-well Non-Skirted, White PCR Plates (ABgene, UK) and MicroAmp Optical Caps (Applied Biosystems, UK). Each reaction was performed on 25 ng diluted total DNA in a volume of 20 μL, under the following conditions: 95°C for 2 minutes; 95°C for 15 seconds, 60°C for 1 minute, repeated 40 cycles; 95 °C for 1 minute, 55 °C for 30 s, 95°C for 30 s. mtDNA/gDNA was generated by normalizing the quantified mtDNA values to the quantified 18S DNA values. To calculate mtDNA damage based on lesion frequency, the DNA of the cells was isolated as described above and assessment of mtDNA damage performed using qPCR as previously described [90]. A common reverse primer (mouse mtDNA reverse: 5’-GCCCAAAGAATCAGAACAGATGC-3’) in combination with primers called either forward *short PCR primer* (5’-GCAAATCCATATTCATCCTTCTCAAC-3’) or forward *long PCR primer* (5’-CCCAGCTACTACCATCATTCAAGTAG-3’). The reverse primer with the short forward primer gives rise to a 16.2 kb product, or with the long forward primer gives rise to a 80 bp product. The lesion frequency of the mtDNA is calculated using the equation below [90]. Chloroquine (CQ) at 40 M was used as positive control (Supplementary Table 5)

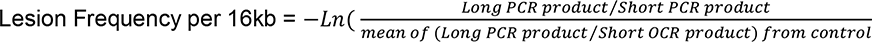

### Transcriptional analysis of GEO datasets

A set of genes relevant to mitochondrial β-oxidation were collated, and then examined for differential gene expression (Supplementary Table 2). Published transcriptome data (NCBI GEO DataSets Accession numbers GSE53053 (https://www.ncbi.nlm.nih.gov/geo/query/acc.cgi?acc=GSE53053); GSE69607 (https://www.ncbi.nlm.nih.gov/geo/query/acc.cgi?acc=GSE69607); GSE84517 (https://www.ncbi.nlm.nih.gov/geo/query/acc.cgi?acc=GSE84517); GSE46903 (https://www.ncbi.nlm.nih.gov/geo/query/acc.cgi?acc=GSE46903); GSE35449 (https://www.ncbi.nlm.nih.gov/geo/query/acc.cgi?acc=GSE35449) and GSE5099 (https://www.ncbi.nlm.nih.gov/geo/query/acc.cgi?acc=GSE5099) regarding different macrophage phenotypes (M0 or M1 from both murine and human origin) generated using similar experimental conditions to those in the current study, were downloaded. All datasets were processed in R and using packages from Bioconductor. Affymetrix microarray data (GSE69607, GSE5099) were normalized and processed using Limma (3.40.6) and Oligo Bioconductor (1.48.0) packages [91, 92]. Illumina beadchip microarray data (GSE46903, GSE35449) were normalized and processed using Limma and lumi (2.36.0) Bioconductor packages [91, 93]. Paired-end reads from Illumina sequencing (GSE53053, GSE84517) were trimmed with Trim Galore (0.5.0 and assessed for quality using FastQC (0.11.8), using default parameters [94, 95]. Reads were mapped to the reference sequence (human GRCh38 or mouse GRCm38) reference genome using STAR (2.7.0) and counts were assigned to transcripts using featureCounts (2.0.0) with the Ensembl gene build GTF (GRCh38.96 or GRCm38.84)[96, 97]. Both the reference genome and GTF were downloaded from the Ensembl FTP site (http://www.ensembl.org/info/data/ftp/index.html/). Differential gene expression analyses used the DESeq2 (1.24.0) package [98]. Genes were discarded from the analysis differential expression if they failed to be significant (significance: adj.pval < 0.05, Benjamini-Hochberg correction for multiple testing).

### Analysis of human microarray data

Data from a published study was obtained, from GSE25504 (https://www.ncbi.nlm.nih.gov/geo/query/acc.cgi?acc=GSE25504) [57]. Here, whole blood was analyzed using Illumina Human Whole-Genome Expression BeadChip HT12v3 microarrays comprising 48,802 features (human gene probes) from neonates with proven microbiological evidence of infection from a usually sterile body site (35 controls, 26 bacterial confirmed sepsis cases). These patients also demonstrated significantly elevated expression markers which form part of the published classifier for sepsis [57]. Log2 expression data for the set of relevant genes (Supplementary Table 2) was compared, Students t-test, significance: adj.pval < 0.05, Benjamini-Hochberg correction for multiple testing.

### Analysis of mouse in vivo bacterial peritonitis gene expression

The SES model was carried out and lavage harvested at 3 or 6 hrs as described [58], data is available as: E-MTAB-10739 (https://www.ebi.ac.uk/biosamples/samples/SAMEA9418371). 8-12 weeks old mixed gender wild type C57Bl/6 mice were from Charles River. Parietal peritoneal tissue was extracted at 3 and 6 h post SES administration, and collected using aseptic techniques. Two sections of lining were immediately snap frozen in liquid N_2_ and stored at −80 °C prior to total RNA extraction. For this, peritoneal membrane sections (80 mg) were dissociated in 1 ml buffer RLT (QIAGEN) supplemented with β-mercaptoethanol (1:100 v:v) using a handheld electric homogenizer (Benchmark Scientific). Lysate was diluted 1:3 in distilled water and digested in 0.2 mg/ml proteinase-K (Invitrogen; 25530049) for 10 min at 55 °C. Lysate was cleared and RNA precipitated in 70 % ethanol. Total RNA was extracted using the RNeasy minikit (QIAGEN; 24136) in accordance with the manufacturer’s instructions. RNA was eluted in 50 μl RNase-free water and quantified using a nanodrop 2000. The integrity and quality of RNA preparations was assessed using an Agilent 2100 bioanalyzer. Samples with an RNA integrity number (RIN) exceeding 8 were used for library preparation (2-4 μg input). Cytoplasmic, mitochondrial, and ribosomal RNA was depleted using the Ribominus transcriptome isolation kit (Ambion; K155001). Libraries were prepared using the RNA-seq kit v2 (Life technologies; 4475936) and sequencing on an ion torrent (Thermo Fisher). Raw data was mapped using Torrent Suite^TM^ (STAR and Bowie2 (2.3.4.3) aligners) using mm10 as the reference genome. Library quality was assessed using FastQC (0.11.8). Differential gene expression analysis was completed with DEseq2 (Bioconductor, 1.24.0). For the specific genes tested (Supplementary Table 2), Students t-test was used, significance: adj.pval < 0.05, Benjamini-Hochberg correction for multiple testing.

#### Analysis of 12-HETE incorporation into PLs by RAW264 cells

For MS precursor scanning, 1×10^6^ RAW 264 cells were seeded in a 6-well plate in DMEM supplemented with 10% FBS. The next day, cells were washed twice with PBS and 2 ml serum free RPMI (no phenol red) was added to each well. Cells were treated for 3 hours with 100 ng/ml LPS, with and without 2.8 μg 12(S)-HETE/10^6^cells. Following 3-hour incubation, cells were scraped into their media and immediately transferred to 2.5ml extraction solvent (hexane:isopropanol) and extracted as described in Methods. For subsequent experiments to quantify 12-HETE incorporated into PLs, 2×10^6^ cells were seeded into 6-well plates in DMEM supplemented with 10% FBS. The next day, cells were washed twice with PBS and 1 ml serum free RPMI (no phenol red) was added to each well. Cells were treated for 3 hours with and without 100 ng/ml LPS and with and without 1.4 μg 12(S)-HETE/10^6^cells. Following a 3-hour incubation, cells were scraped directly into their culture media and immediately transferred to 2.5ml extraction solvent and extracted using the hexane:isopropanol method, as described in the methods section. Lipids were analysed on a Sciex 6500 QTrap, with chromatography as follows: Luna 3μm C18 150x 2mm column (Phenomenex, Torrance, CA) with a gradient of 50–100% B over 10 min followed by 30 min at 100 % B (A, methanol:acetonitrile:water, 1 mM ammonium acetate, 60:20:20; B, methanol, 1 mM ammonium acetate) with a flow rate of 200 μl/min. Source and MS conditions were as follows: CUR 35, IS −4500, TEM 500, GS1 40, GS2 30, DP -50, CE -38, CXP -11. First a precursor scan (PREC 319.2) was performed in negative mode to determine a list of precursor PE ions present in the cell extract, scanning a mass range of 600 - 950 Da at a scan rate of 1000 Da/s. Following this, 10 ions of interest were identified and EPI scans were performed for each precursor ion to obtain MS/MS spectra. Source and chromatography conditions were identical to above. Product ions were scanned at 1000 Da/sec with a mass range of 100-820 Da, using a dynamic fill time in the trap. Once candidate lipids were identified, extracts were analysed in MRM mode, monitoring the precursor ion to the 12-HETE internal product ion (*m/z* 179.1) (Supplementary Table 6). Samples were spiked with internal standards 15:0-18:1(d7) PE (10ng) and 15:0-18:1(d7) PC (300ng), and analytes quantified against a calibration curve using 18:0a/12-HETE-PE and 18:0a/12-HETE-PC as primary standards. MRMs were as follows: *m/z* [M-H]^-^ 738.6-179.1, 754.6-179.1, 764.6-179.1, 766.6-179.1, 780.6-179.1, 782.6-179.1, 808.7-179.1, 810.7-179.1 Chromatography and source conditions were the same as above. Peak inclusion criteria were those that had at least 7 points across the peak and exceeding a signal:noise ratio of 5:1.

#### LC-MS/MS of free fatty acids (FFA)

FFA were extracted from cell culture cells and supernatant (1 ml) using the hexane:isopropanol method described above. An internal standard mix was added (10 µl) to samples prior to extraction (Supplementary Table 7). FFA were derivatized in sample extracts according to Han et al (2015) with modifications[99]. Dried extracts were reconstituted in methanol (100 µL), then 50 uL 3NPH (200 mM 3-nitrophenyl-hydrazinein, 50/50 Methanol/H_2_O) and 50uL EDC/PYR (120 mM 1-Ethyl-3-(3-dimethylaminopropyl)carbodiimide and 6 % pyridine, 50/50 Methanol/H_2_O) were added. Samples were vortexed then incubated for 30 min at 40 °C. Excess derivatization reagents were quenched by addition of 0.5 % formic acid (100 µL; 75/25 Methanol/H_2_O) and incubation at 40 °C for 30 min. Samples were aliquoted into HPLC vials for LC/MS/MS in MRM mode on a Nexera liquid chromatography system (Shimadzu) coupled to a QTRAP 4000 mass spectrometer with an ESI source (Sciex). Liquid chromatography was performed using a C18 reverse phase column (Kinetex Polar C18 Column; 100Å, 100 × 2.1 mm, 2.6 µm particle size, Phenomenex). Two mobile phases were used: 100 % H_2_O + 0.1 % formic acid (mobile phase A) and 100 % methanol + 0.1 % formic acid (mobile phase B). Gradient was 15 % B 0-2.47 min, increasing to 55 % B at 11.12 min; then to 100 % B at 20 min; held at 100 % B up to 32 min; then 10 min re-equilibration at 15 % B. The flow rate was at 0.2 ml/min and injection volume 2µL. Column temperature, 35 °C. MRM transitions, and MS settings (declustering potential and collision energy) for individual FFA are shown in Supplementary Table 7. FFA were detected in negative ion mode and the method was scheduled with fixed retention times and 40 sec acquisition windows. Analytes were quantified based on the ratio of response to deuterated internal standards in comparison to externally generated standard curves for primary standards using MultiQuant software, V3.0.2.

#### Statistics

For lipidomics experiments comparing more than two conditions, one way ANOVA with Tukey post hoc test is used (https://astatsa.com/OneWay_Anova_with_TukeyHSD/_get_data/). Although all conditions are compared, our study is focused on the impact of CPT1 inhibition/silencing, so our text and figures generally show those comparisons. Where only two conditions are compared, Student’s T test was used. For the complete information on all comparisons, ANOVA and T test results are also provided in our supplementary data file (Supplementary Data.xls). Where no significant difference was seen, no comparison bar is shown. Where lipids were not detected in some samples of a dataset, their values were replaced with corresponding 50%LOQ values prior to conducting statistical analysis. These were calculated using the LOQ for the corresponding primary standards, taking into account sample volumes injected on column. Figures show zero values, and all 50%LOQ values are provided (in red text) in supplementary data, where used, but the zero value was plotted on figures. If a lipid was not detected in any sample, it was not included in the dataset, and the full list of lipids measured in all samples is provided in Supplementary Data File 2. All p values are provided in Source Data. For statistical analysis, Prizm (7-9) or Excel (2013, 2016 and office 365 versions) were used.

## Data Availability

The raw numbers for charts and graphs are available in the Source Data file whenever possible. All data are included in the Article and its Supplementary Information, or are available from the authors upon reasonable requests.

## Acknowledgements.

Funding from Wellcome Trust (094143/Z/10/Z) and European Research Foundation (LipidArrays) is gratefully acknowledged (VBO). VBO is a Royal Society Wolfson Research Merit Award Holder and acknowledges funding for LIPID MAPS from Wellcome Trust (203014/Z/16/Z). Ser Cymru Project Sepsis grant funded by WG/EU-ERDF (PG, VBO). PRT is funded by a Wellcome Trust Investigator Award (107964/Z/15/Z) and the UK Dementia Research Institute. MAC is funded by BBSRC Discovery Fellowship (BB/T009543/1). VDU acknowledges The Nathan Shock Center P30 AG05088. Kidney Research UK (RP-024-20160304, SAJ), Versus Arthritis (Reference 20770 awarded to SAJ, VOD). MAC is supported by BBSRC Discovery Fellowship (BB/T009543/1). We gratefully acknowledge expert input from Alan Brash, Vanderbilt University, relating to lipid structural elucidation, James Burston for supporting analysis of mitochondrial function, Widad Dantoft for provision of the human sepsis dataset, William Watkins for assistance with statistical analysis, and our colleague Javier Uceda Fernandez (deceased), Cardiff University for the murine transcriptomics dataset.

## Author Contributions statement

MM, KK, LD, VJT, PRSR, GAB, CH, PK, MR, JML, MAC, MA, MG, DAW conducted experiments, MM, KK, CH, LD, JML, PG, VDU, DAW, SAJ, PRT designed experiments, RA, SD RCM, MM, KK, VJT, PRSR, GAB, CH, PK, MR, DAW, JML, LD analysed data, NHS, SM implemented and supported data analysis, VOD drafted the manuscript and all authors edited the manuscript.

## Competing Interests Statement

Dr Paul Kennedy is an employee of Cayman Chemical. All other authors declare no competing interests.

**Figure legends:** Source data are provided as a Source Data file for all figures.

---

**Supplementary Figure 1.**
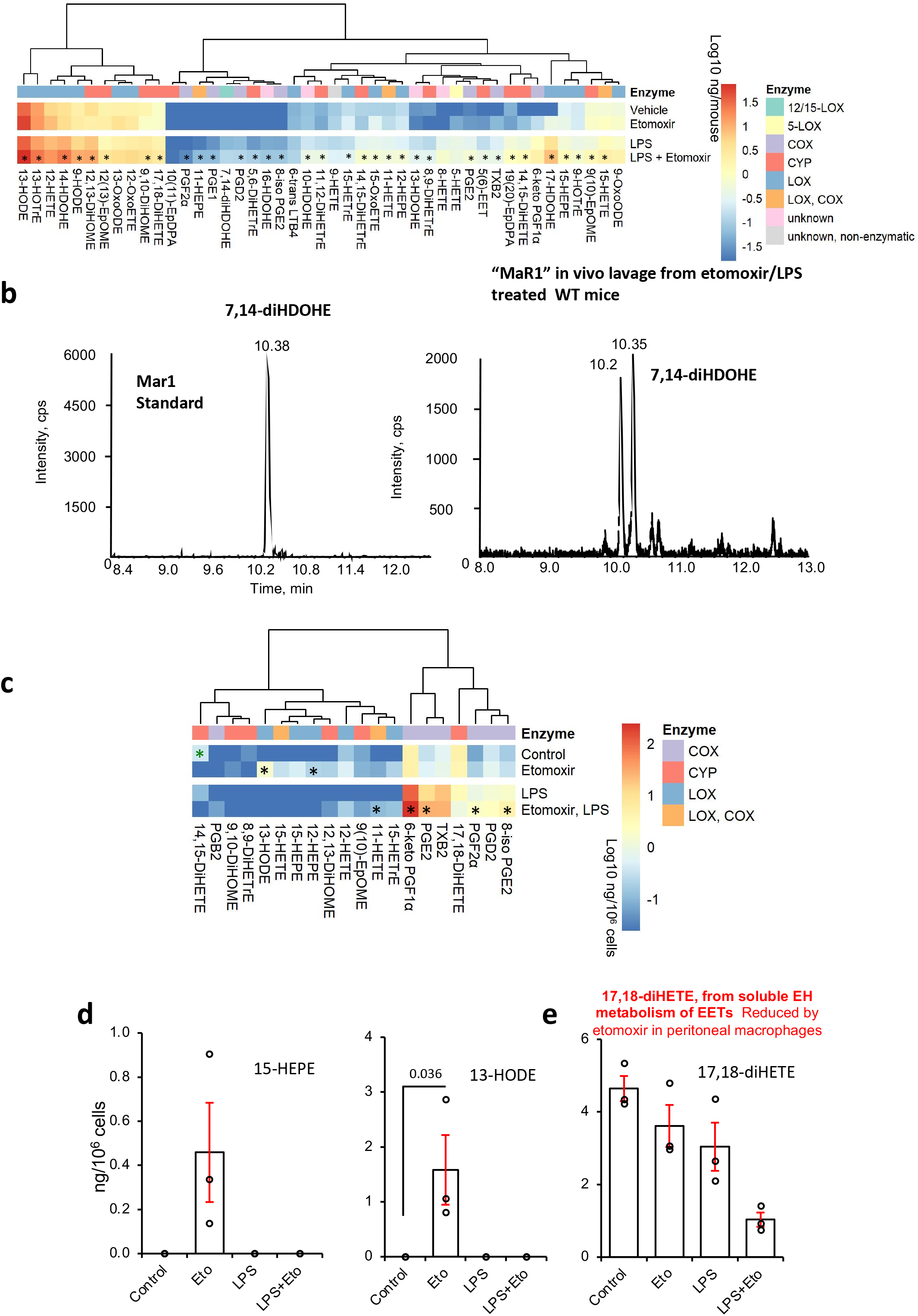
Inhibition of CPT1 increases oxylipin levels significantly during inflammation *in vivo,* LC/MS/MS of “Mar1” in mouse peritonitis in response to LPS and the impact of CPT1 inhibition on oxylipins in naïve peritoneal macrophages. *Panel A. Inhibition of CPT1 modulates oxylipin metabolism in vivo.* Wild type mice (female, 7-9 weeks) were injected i.p. with vehicle (PBS), etomoxir (100 μg) or LPS (1 μg). After 6 hrs, lavage was harvested and lipids extracted using SPE then analyzed using LC/MS/MS as outlined in Methods. A heatmap was generated using Pheatmap as described in Methods, with data as log10 ng/total mouse lavage for all oxylipins (n = 10), data is mean +/- SEM, * p < 0.05, one way ANOVA with Tukey post hoc test, stats are shown for effect of etomoxir only, where significant. For full ANOVA analysis see Supplementary Data.xls. Tree shows hierarchical clustering. *Panel B. LC/MS/MS of lipid extract from peritoneal lavage (mouse treated with etomoxir and LPS)*. A lipid with same retention time as Mar1 was detected in lavage from mice treated with etomoxir and LPS *in vivo*, labelled herein as 7,14-diHDOHE. Standard is shown in left panel, mouse lipid in right panel*. Panel C. Heatmap shows that many lipids are elevated when CPT1 is inhibited in vitro*. Peritoneal macrophages were isolated as described in Methods, then cultured in serum-free medium in the presence of etomoxir (25 μM) with/without LPS (100 ng/ml). After 24 hrs, supernatant was harvested and lipids extracted using SPE, then analyzed using LC/MS/MS. A heatmap was generated using Pheatmap as described in Methods, with data as log10 ng/10^6^ for all oxylipins (n = 3, data is mean +/- SEM, * p < 0.01, one way ANOVA with Tukey post hoc test, stats are shown for effect of etomoxir only, where significant, black: significant elevation with etomoxir, compared to vehicle or LPS alone, green: significant reduction with etomoxir, compared to vehicle or LPS alone). Tree shows hierarchical clustering. *Panels D,E. The impact of LPS and etomoxir on generation of selected oxylipins, measured by LC/MS/MS, in wild type naïve peritoneal macrophages*. Supernatant from peritoneal macrophages was analyzed for oxylipins following culture of macrophages as outlined in methods, for 24 hrs with LPS (100 ng/ml) +/- etomoxir (25 μM). (n = 3, data is mean +/- SEM, separate wells of cells). * p < 0.05, one way ANOVA with Tukey post hoc test, stats are shown for effect of etomoxir only, where significant. Where no stars are shown, no significant difference was seen.

**Supplementary Figure 2.**
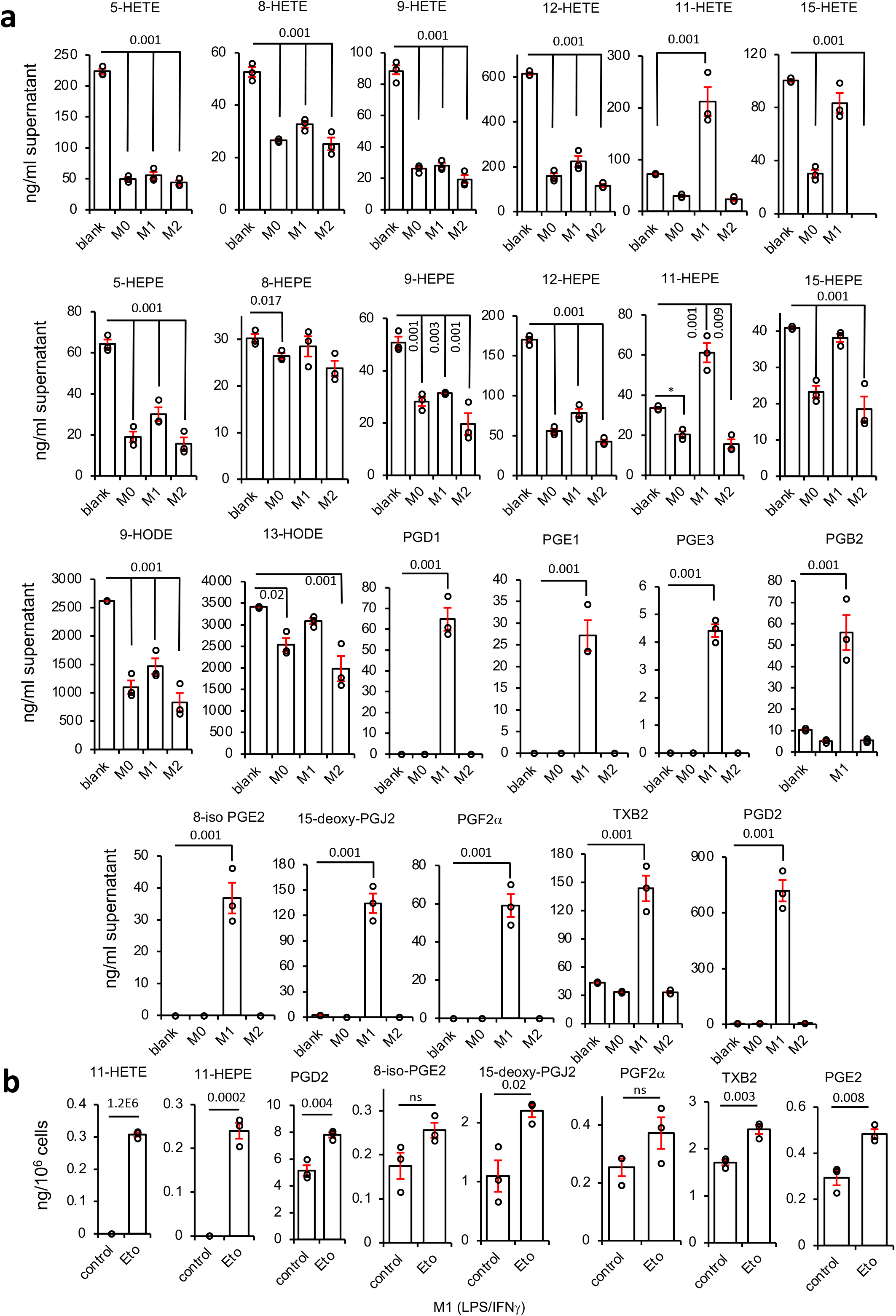
BMDM-derived M0, M1 and M2 cells consume diverse oxylipins from serum, while M1 cells consume prostaglandins via CPT1. *Panel A. Serum derived mono-hydroxy oxylipins are metabolized by M0, M1 and M2 cells, while M1 cells generate prostaglandins that are low abundance in serum.* BMDM-derived M0 (M-CSF), M1 (LPS/IFNg), and M2 (IL-4) cells were derived during a 24 hr culture in medium containing 10 % FCS, as described in Methods, then supernatant extracted and analyzed for oxylipins using LC/MS/MS as described in Methods. (n = 3, separate wells of cells). Data is mean +/- SEM, one way ANOVA with Tukey post hoc test, stats are shown for comparison with blank, where significant. Where no stars are shown, no significant difference was seen. *Panel B. M1 cell secretion of prostaglandins is increased by CPT1 inhibition*. BMDM-derived M1 cells were treated in the absence (control) or presence (etomoxir) of etomoxir (25 µM) for 24hr. Here, FCS-free medium was used to avoid contaminating oxylipins from serum (n = 3, mean +/- SEM. Separate wells of cells, with/without etomoxir, Student’s T test, two-tailed).

**Supplementary Figure 3.**
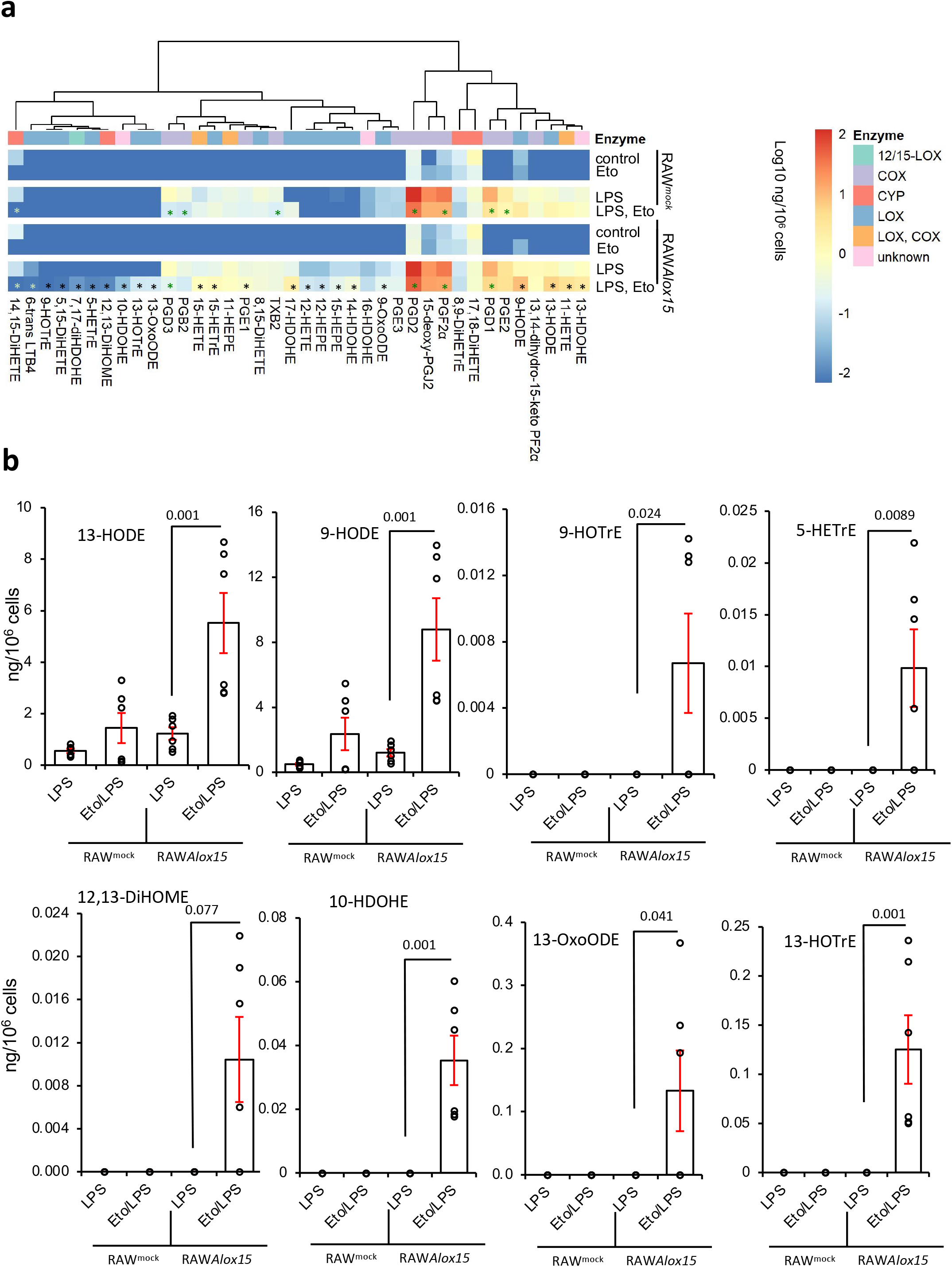
Inhibition of CPT1 increases secretion of 12/15-LOX-derived eicosanoids from RAW cells overexpressing *Alox15.* *Panel A.* A heatmap was generated using Pheatmap as described in Methods. Raw^mock^ and Raw*Alox12* cells were cultured with etomoxir (25 mM) and/or LPS (100 ng/ml) for 24 hr before harvest and SPE extraction of supernatant for oxylipin analysis using LC/MS/MS. Data are shown as log10 ng/10^6^ values for all oxylipins. black: significant elevation compared to vehicle or LPS alone, green: significant reduction compared to vehicle or LPS alone, p<0.05, one way ANOVA with Tukey post hoc test was carried out on the LPS treated samples as a group, stats are shown for effect of etomoxir only, where significant). Tree shows hierarchical clustering. *Panel B. Selected lipids shown in more detail.* (n = 6, separate wells of cells). Data is mean +/- SEM, one way ANOVA with Tukey post hoc test was carried out on the LPS treated samples as a group, stats are shown for effect of etomoxir only, where significant. Where no stars are shown, no significant difference was seen

**Supplementary Figure 4.**
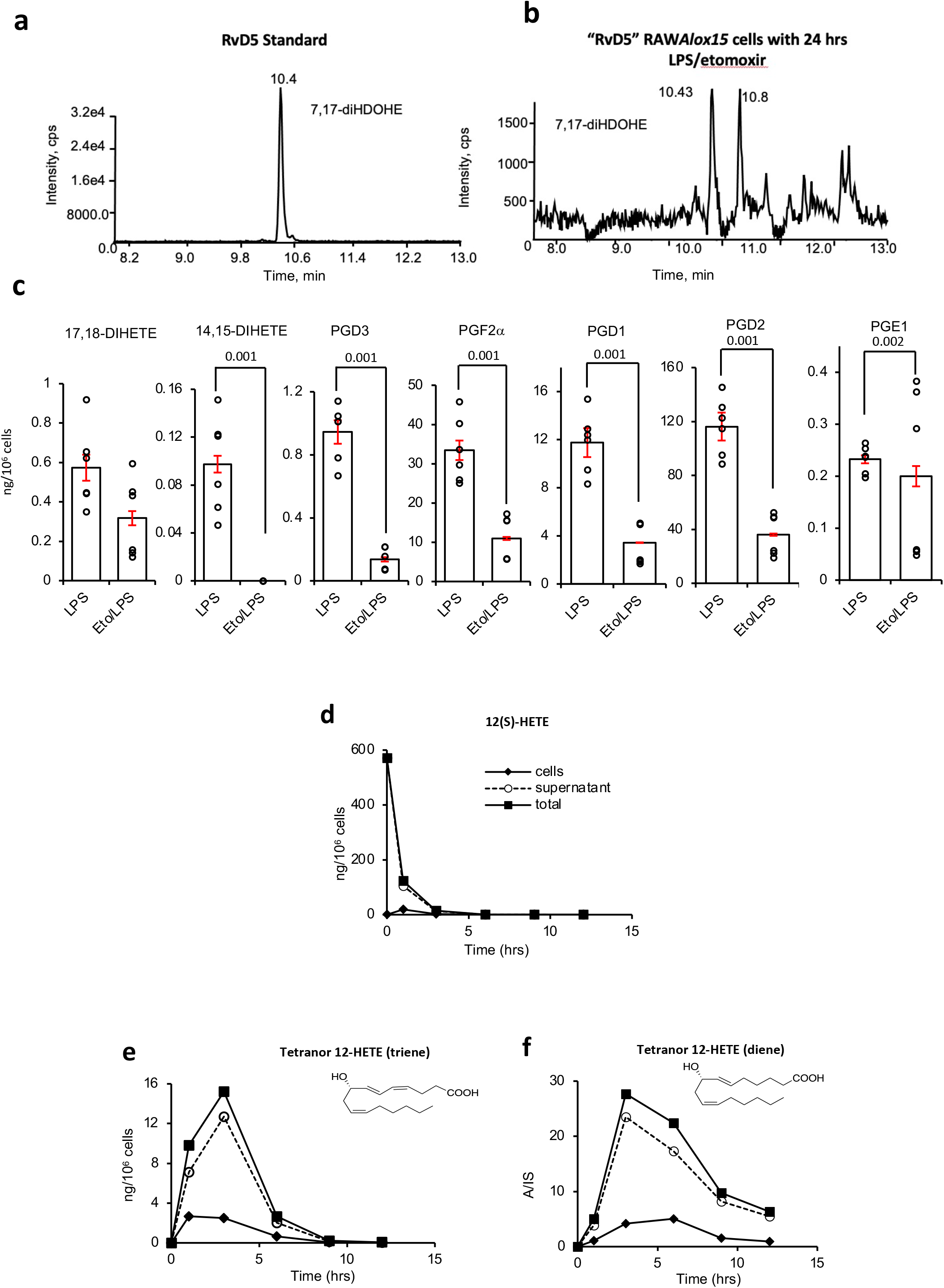
Representative chromatograms for 7,17-diHDOHE (“RvD5”) in RAWAlox15 cells, impact of etomoxir on 17,18-di-HETE and PGs in RAW cells, and timecourse of 12-HETE metabolism in RAW cells. *Panel A,B. Supernatant from RAWAlox15 cells treated with LPS/etomoxir contains isomers of 7,17-di-HDOHE*. Cells were cultured with LPS/etomoxir for 24 hrs as in Methods, then supernatant was analyzed for the presence of RvD5 using LC/MS/MS/MS (Panel B). Authentic standard is shown for comparison (Panel A). *Panel C. Etomoxir reduces generation of 17,18-diHETE and PGs from RAW cells*. RAW cells were cultured for 24 hr with/without etomoxir (dose) and LPS (dose) before supernatant was harvested and analyzed using LC/MS/MS as in Methods. (n = 6, mean +/- SEM, separate wells of cells), one way ANOVA with Tukey post hoc test was carried out on the full dataset of RAW^mock^ and RAW*Alox15* but data on RAW^mock^ cells is shown since these lipids are generated by COX or CYP (see Supplementary data). Stats are shown for effect of etomoxir only, where significant. Where no stars are shown, no significant difference was seen. *Panels D-F. 12-HETE is rapidly consumed and converted to two tetranor 12-HETE metabolites by RAW cells*. RAW cells were cultured for up to 12 hr in serum free medium with 1.4 mg 12(S)-HETE added per 10^6^ cells. At varying timepoints, samples were harvested and lipids extracted and analyzed using LC/MS/MS as described in Methods.

**Supplementary Figure 5.**
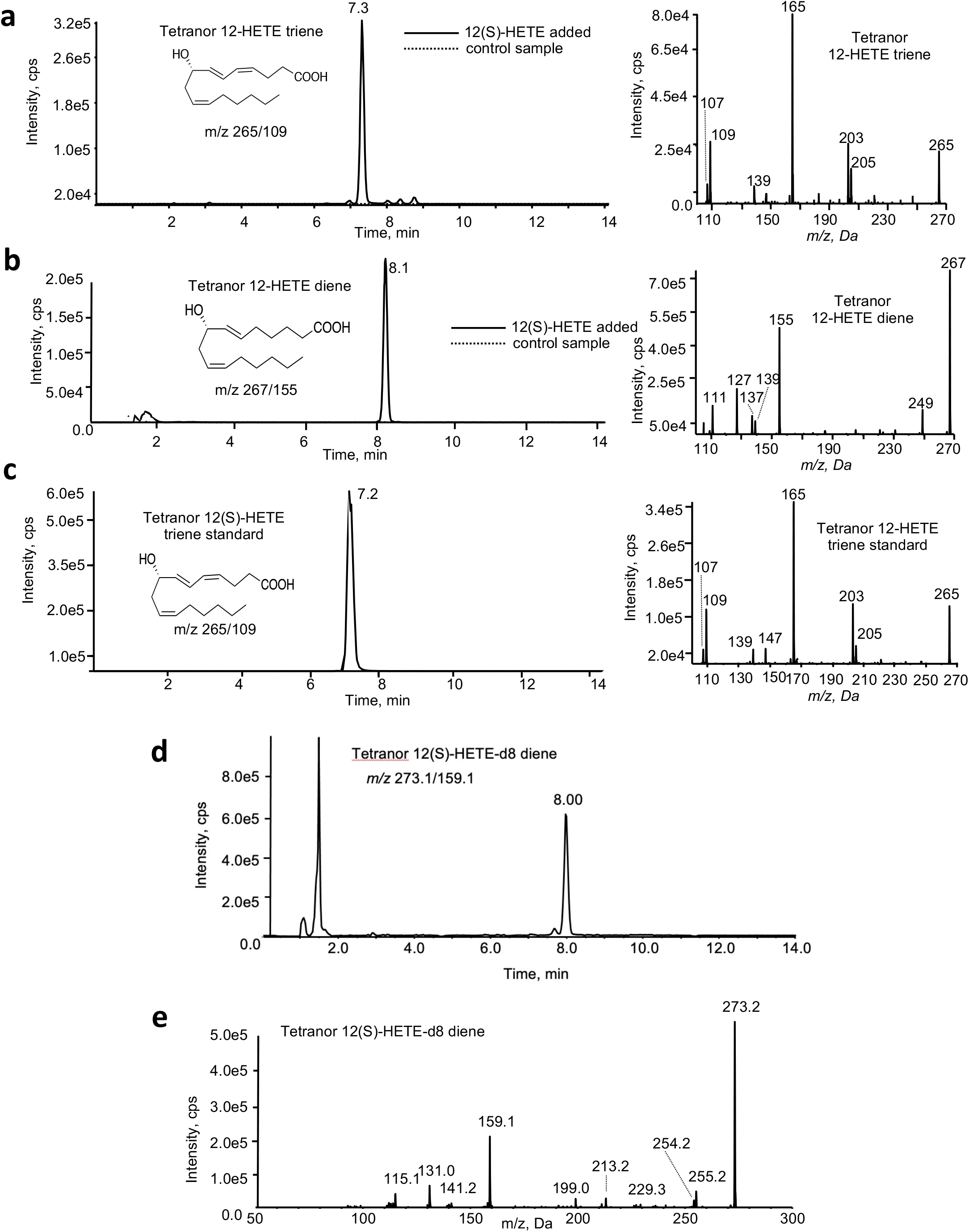
Mass spectrometry confirmation of formation of two tetranor 12-HETE metabolites in RAW cells. 12(S)-HETE or 12-HETE-d8 (1.5 mg) were incubated with RAW cells for 3 hrs and then supernatants harvested for LC/MS/MS. *Panels A,B. Formation of tetranor diene and triene 12-HETEs.* A representative sample is shown for each, including chromatogram and MS/MS spectrum, with control (dotted line) and 12(S)-HETE-supplemented (solid line) samples overlaid. *Panel C. MS/MS of the tetranor triene 12-HETE standard*. *Panels D,E. Formation of the 12(S)-HETE-d8 tetranor diene, following supplementation of 12(S)-HETE-d8 to RAW cells (1.5 mg/10^6^ cells for 3 hrs),* showing a representative chromatogram (D) and MS/MS spectrum (E).

**Supplementary Figure 6.**
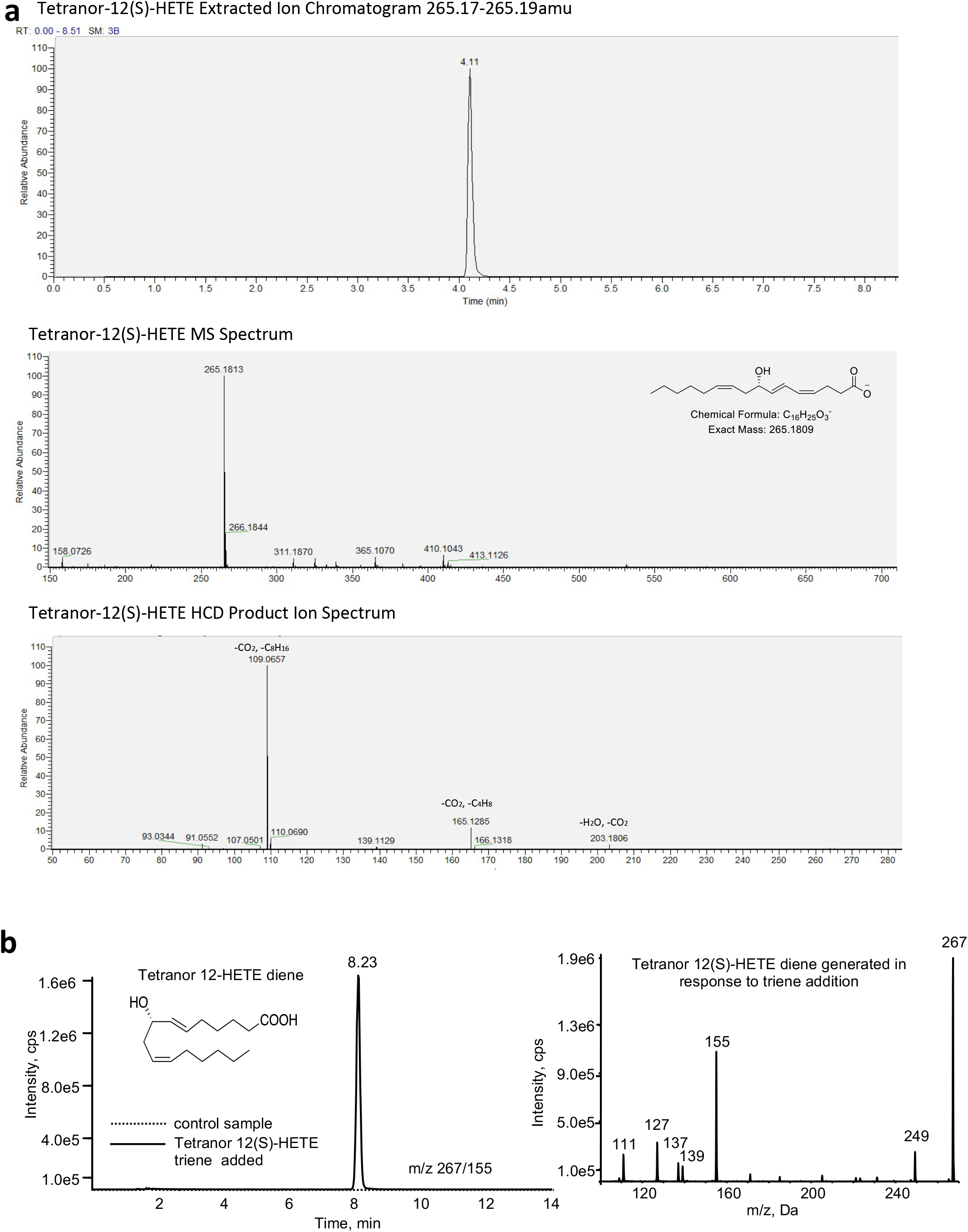
High resolution MS/MS of the 12-HETE tetranor triene standard and tetranor triene 12(S)-HETE is converted into the diene isomer in macrophages. *Panel A.* Extracted ion chromatogram, high resolution MS spectrum, and MS/MS spectrum of the tetranor-12(S)-HETE triene standard. *Panel B. Addition of tetranor triene 12(S)-HETE leads to formation and secretion of the diene isomer*. 1.2 mg triene standard was incubated per 10^6^ RAW cells for 3 hr, then supernatants harvested and analyzed using LC/MS/MS.

**Supplementary Figure 7.**
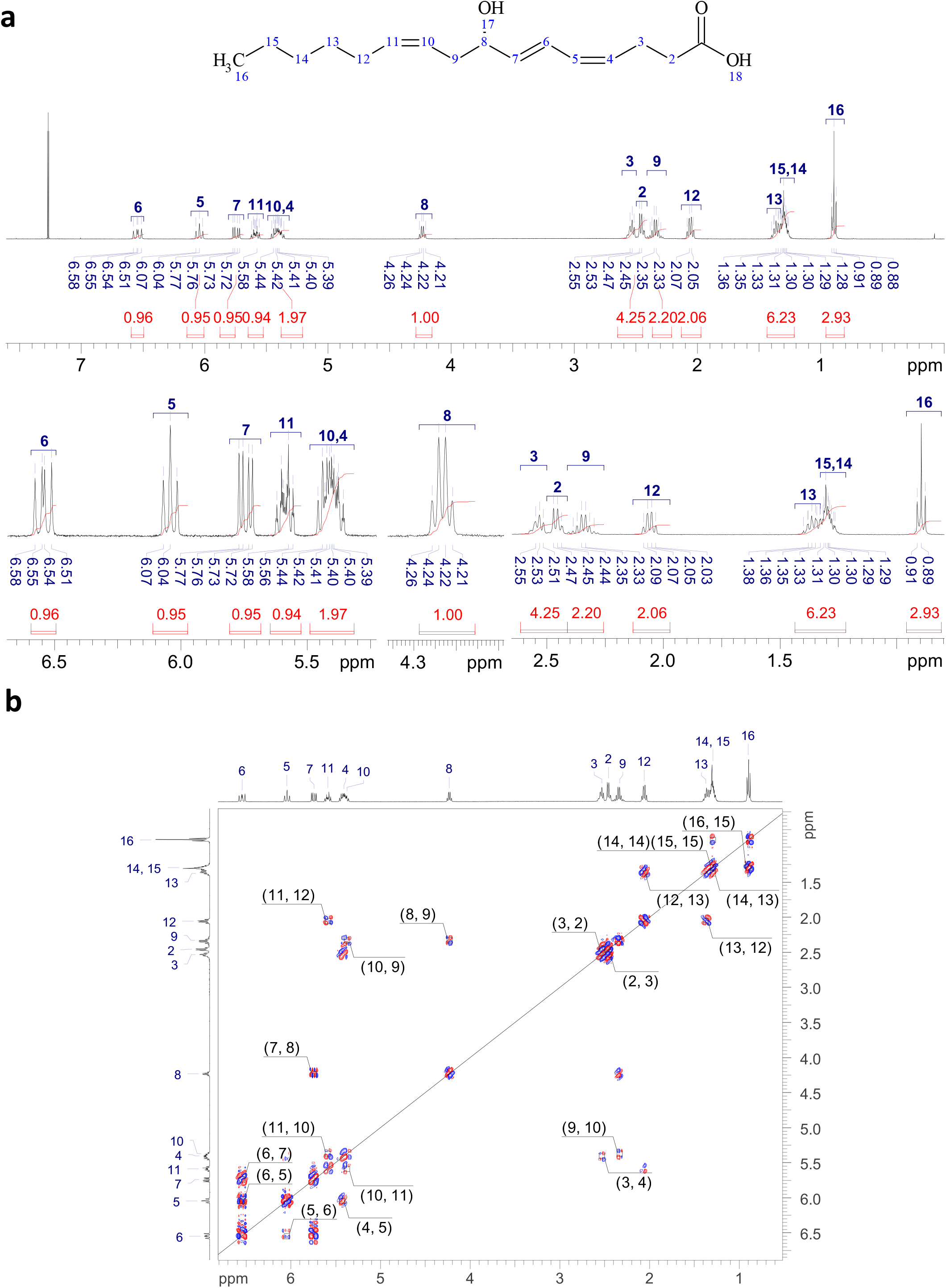
^1^H and ^1^H-^1^H COSY NMR spectra of the tetranor-12(S)-HETE standard. *^1^H (Panel A) and ^1^H-^1^H COSY (Panel B) NMR* demonstrates that the H11 vinyl proton at 5.60 ppm is connected to the methylene H12 protons at 2.05 ppm, which itself is connected back to the alkyl protons at 1.33 ppm. This identifies the 5.60 ppm signal as H11, which correlates with one of the protons that overlap at 5.4 ppm (area of two protons). This 5.4 ppm signal must then be the H10 proton as well as the H4 vinyl proton (which correlates with the H5 vinyl proton and H3 methylene protons).

**Supplementary Figure 8.**
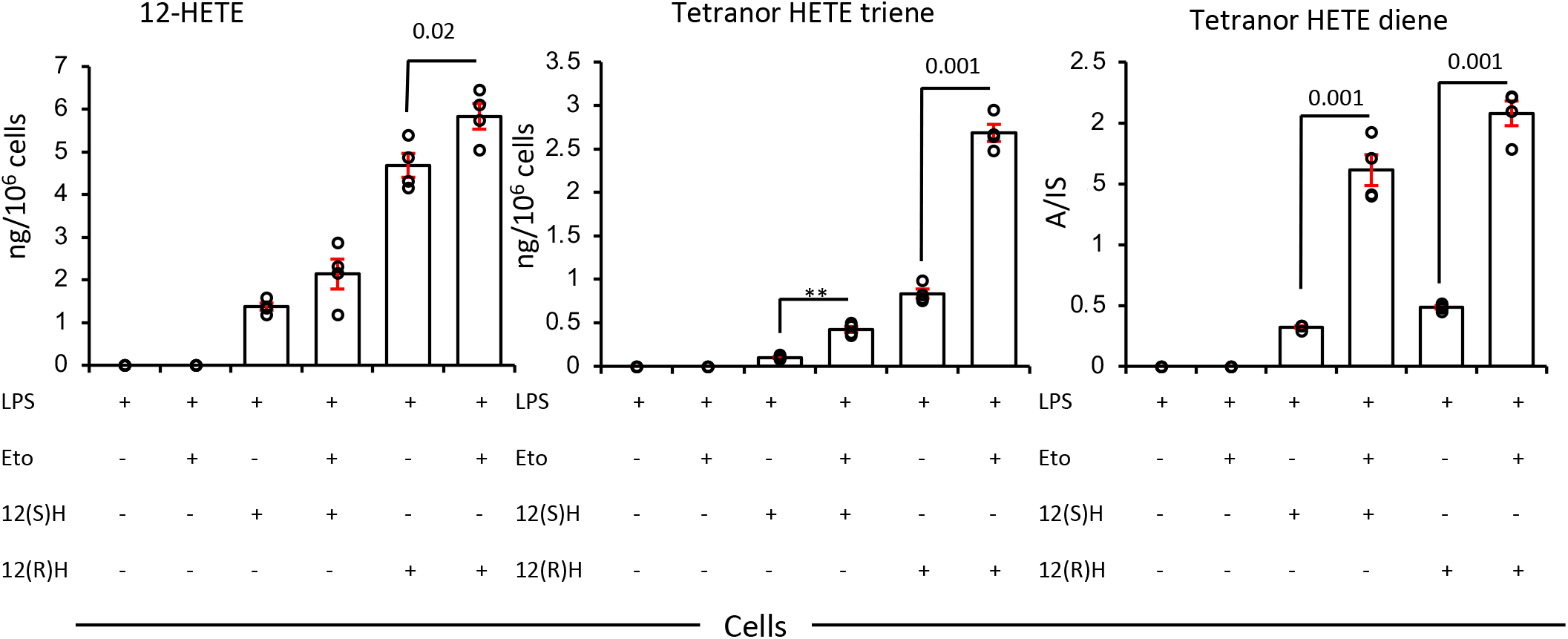
CPT1 inhibition prevents metabolism of 12(S) or 12(R)HETE and their tetranor triene and diene metabolites. RAW cells were supplemented with 1.5 mg 12(S) or 12(R)-HETE/10^6^ cells for 3 hrs with/without etomoxir (25 mM), then cell pellets were analyzed for levels of 12-HETE and its triene and diene tetranor products using LC/MS/MS (n = 4, mean +/- SEM, separate wells of cells). For all panels, comparisons are with/without etomoxir, one way ANOVA with Tukey post hoc test, stats are shown for effect of etomoxir only, where significant.

**Supplementary Figure 9.**
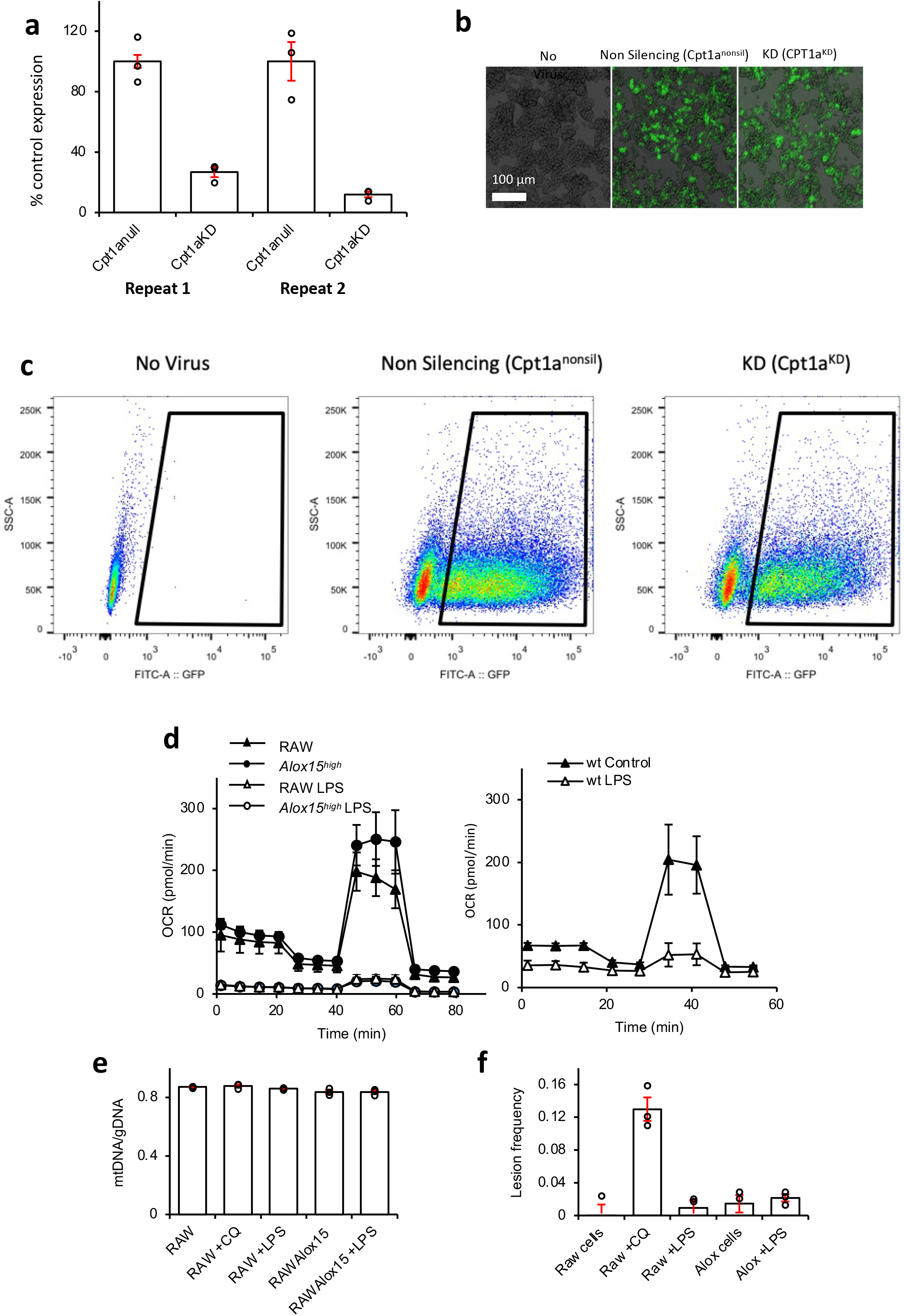
Confirmation of *Cpt1a* knockdown in RAW cells and OxPhos is suppressed in LPS-stimulated macrophages. *Panel A. Cpt1a expression is suppressed in Cpt1a^KD^ cells versus Cpt1a^null^.* Bar graph showing the effect of Cpt1a knockdown shRNA on Cpt1a gene expression in RAW264 macrophages, measured by real-time PCR. (n = 3 per group, separate wells of cells, mean +/- SEM), and two repeats are shown. *Panel B. Images of RAW264 cells infected with viruses* (purified by 2.5 μg/ml puromycin) taken on a 10x objective lens using an EVOS microscope (Life Technologies). Image shows transmitted brightfield overlaid with Green fluorescent GFP. Imaging was performed on one occasion as confirmation of infection. *Panel C. Representative images showing GFP expression and sort gates for RAW264 cells infected with viruses using the BD FACS Aria II flow cytometer*. *Panel D. LPS treatment of RAW or naïve peritoneal macrophages reduces mitochondrial OxPhos*. RAW, RAW*Alox15*, or naïve peritoneal macrophages from wild type or *Alox15^-/-^* mice were treated with LPS (100 ng/ml) for 24 hrs. Mitochondrial function was assessed as described in Methods for peritoneal macrophages (wild type mice) or cell lines (RAW, or *Alox15*^high^) (n = 5, 4 or 5 respectively, mean +/- SEM). *Panels E,F. LPS doesn’t alter mitochondrial DNA or lesion frequency in RAW macrophages.* RAW and RAW*Alox15* cells were mock treated or treated with CQ (40 mM) or LPS (100 ng/ml) for 24 hr. Subsequently their DNA was isolated and used to assess the mtDNA to gDNA ratio (E) and the frequency of lesions in mtDNA (F) using qPCR, one-way ANOVA with Tukey correction, no data were significantly different from RAW samples, (except RAW+CQ, as expected for lesion frequency only, acting as a positive control). Three independent experiments were conducted and a single value for each sample generated in each experiment. These were then averaged (mean +/- SD).

**Supplementary Figure 10.**
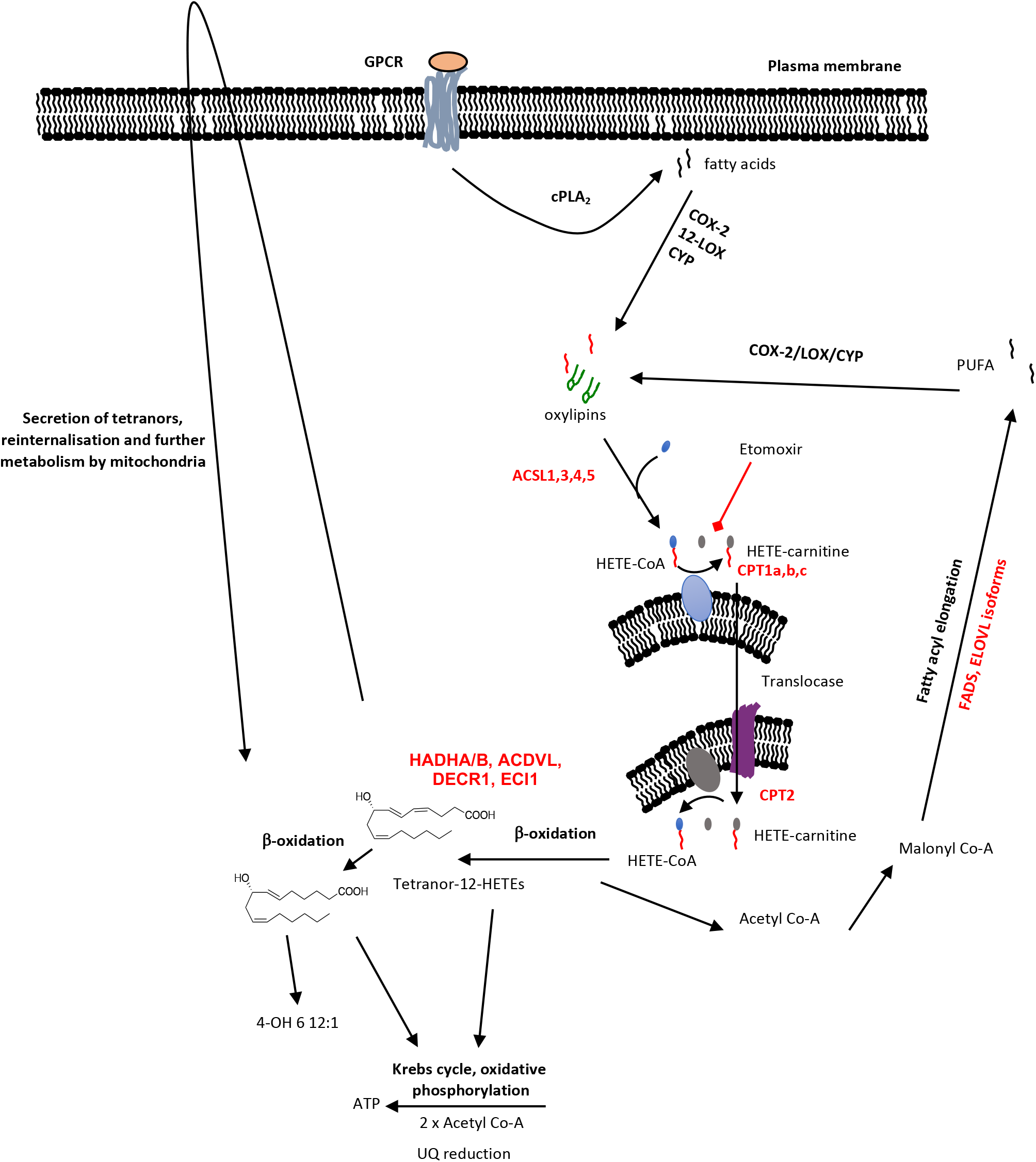
Graphic showing formation and mitochondrial metabolism of oxylipins. *Schematic for pathway for mitochondrial metabolism of oxylipins.*

**Supplementary Figure 11.**
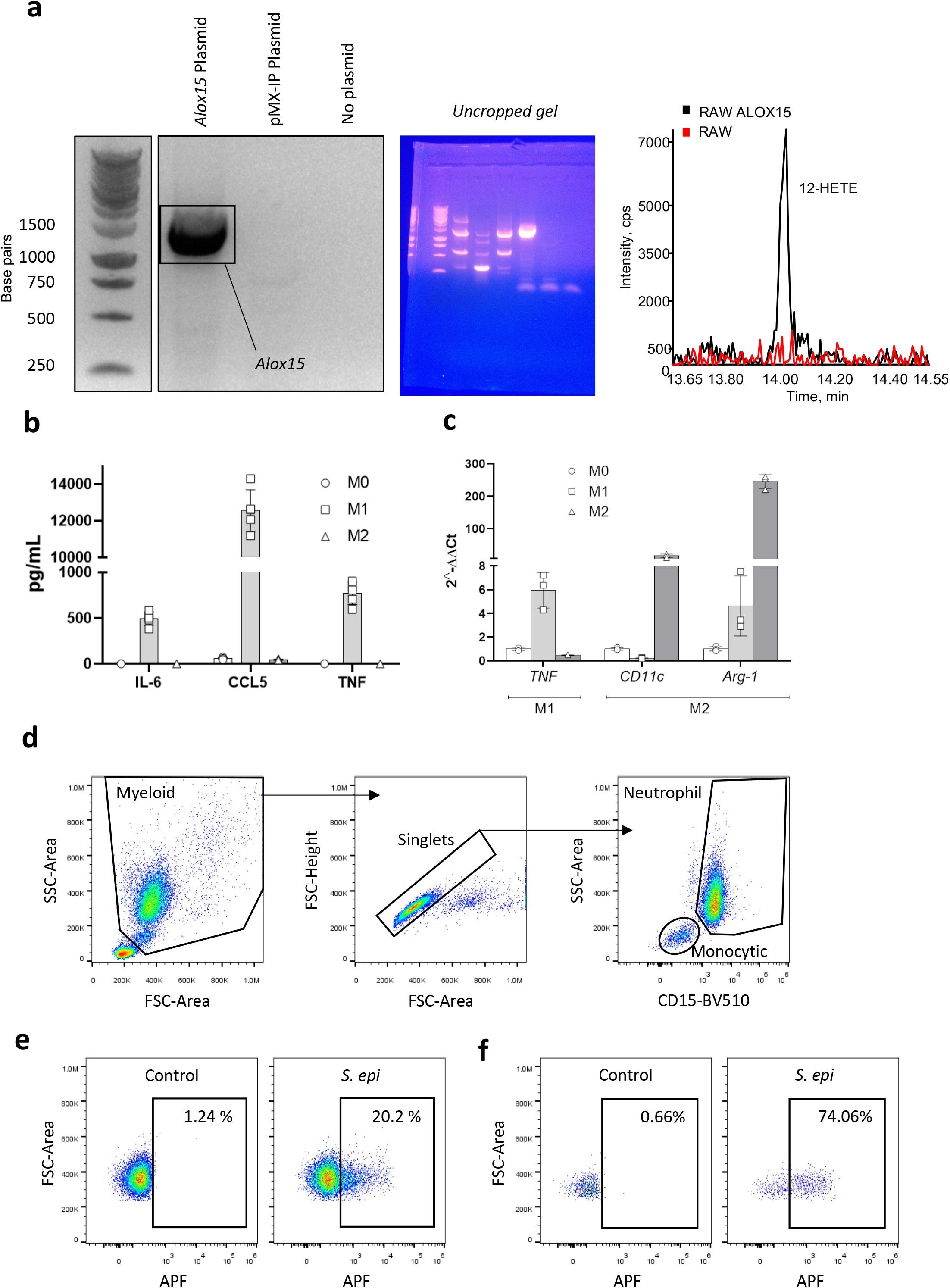
Expression of Alox15 in overexpressing RAW cells, confirmation of phenotype in bone marrow derived macrophages, and gating strategy for neutrophils and monocytes in vitro. *Panel A. Expression of Alox15 in RAW cells.* RNA was isolated from the RAW*Alox15^-/-^* and pMX-IP control cell lines, then converted to cDNA and PCR was performed using primers to amplify *Alox15*. Amplified PCR products were run on a 10% agarose gel and showed that *Alox15* was amplified from the *Alox15* plasmid, but not from the pMX-IP cDNA (plasmid) control, or the no DNA (no plasmid) control, thereby confirming expression as expected. Chromatogram shows generation of 12-HETE in RAWAlox15 cells only, but not mock transfected controls. The full gel is shown (right panel). Unlabelled markers in the left panel (lack of space) are at 10K, 8K, 6K, 5K, 5K, 3K, 2.5K, 2K base pairs. The gel was run once to confirm expression prior to confirmation using the enzyme activity assay. *Panel B*. *M1 macrophages secrete typical cytokines*. Cytokine production of typical M1 phenotype markers (IL-65, CCL5 and TNF) were determined by ELISA analyses of supernatants post differentiation as described in Methods (n = 5, mean +/- SD). *Panel C. Confirmation of classical and alternative activation in BMDM*. RNA was isolated from activated BMDM, converted to cDNA and then analyzed by real time PCR as described in Methods section. Probes for typical phenotype markers are indicated inset, n = 3, mean +/- SD. *Panels D-F. APF flow cytometric gating strategy.* Representative flow cytometry plots showing the gating strategy for neutrophils and monocytic cells, with a distinctive forward-vs side-scatter profile (FSC vs SSC) that distinguishes them from lymphocytes (Panel D). Neutrophils were further distinguished from eosinophils and monocytes via high expression of CD15. Representative gating for APF^+ve^ neutrophils (Panel E) and monocytes (Panel F) is depicted in the presence or absence of *Staphylococcus epidermidis* (*S. epi*).

**Supplementary Figure 12.**
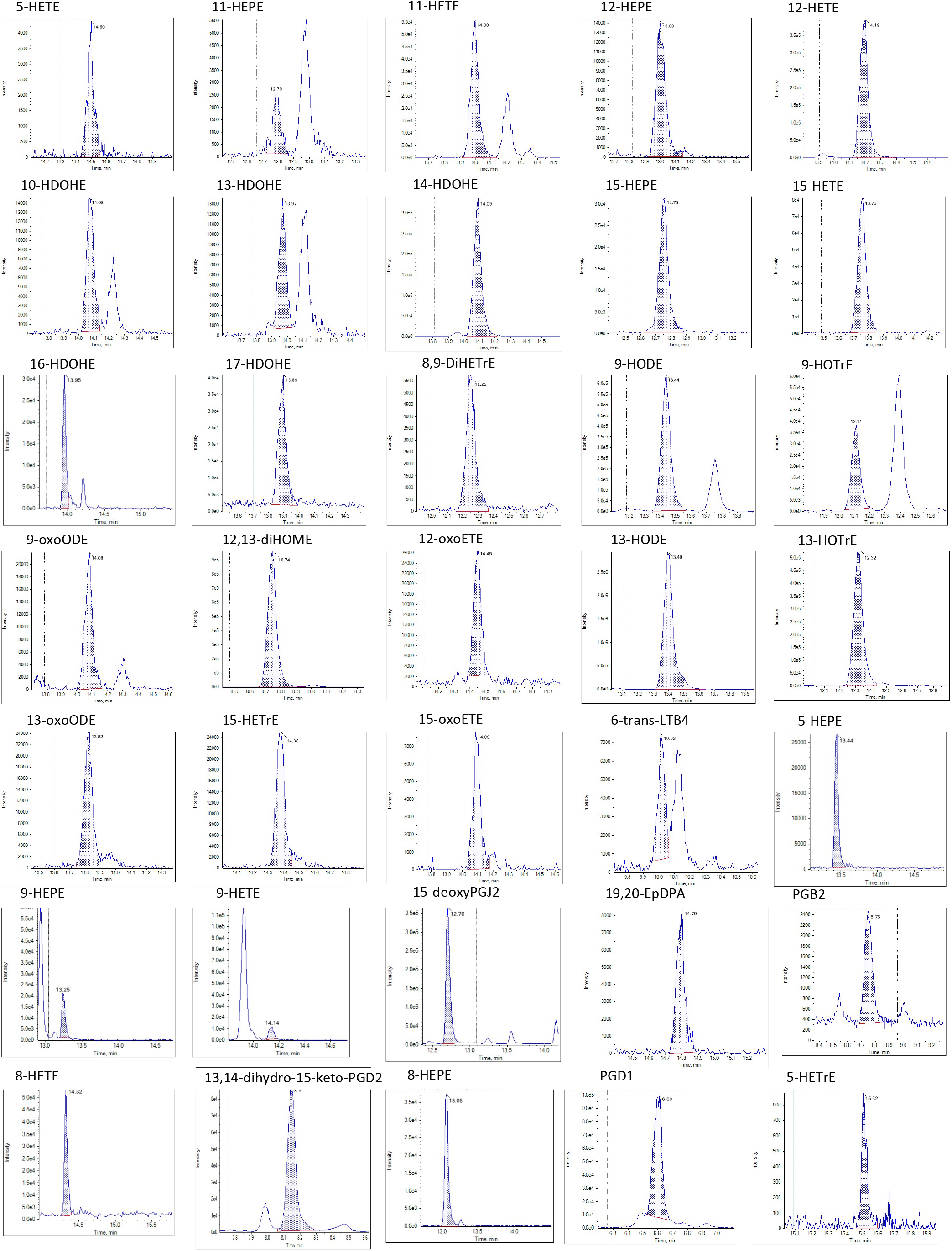
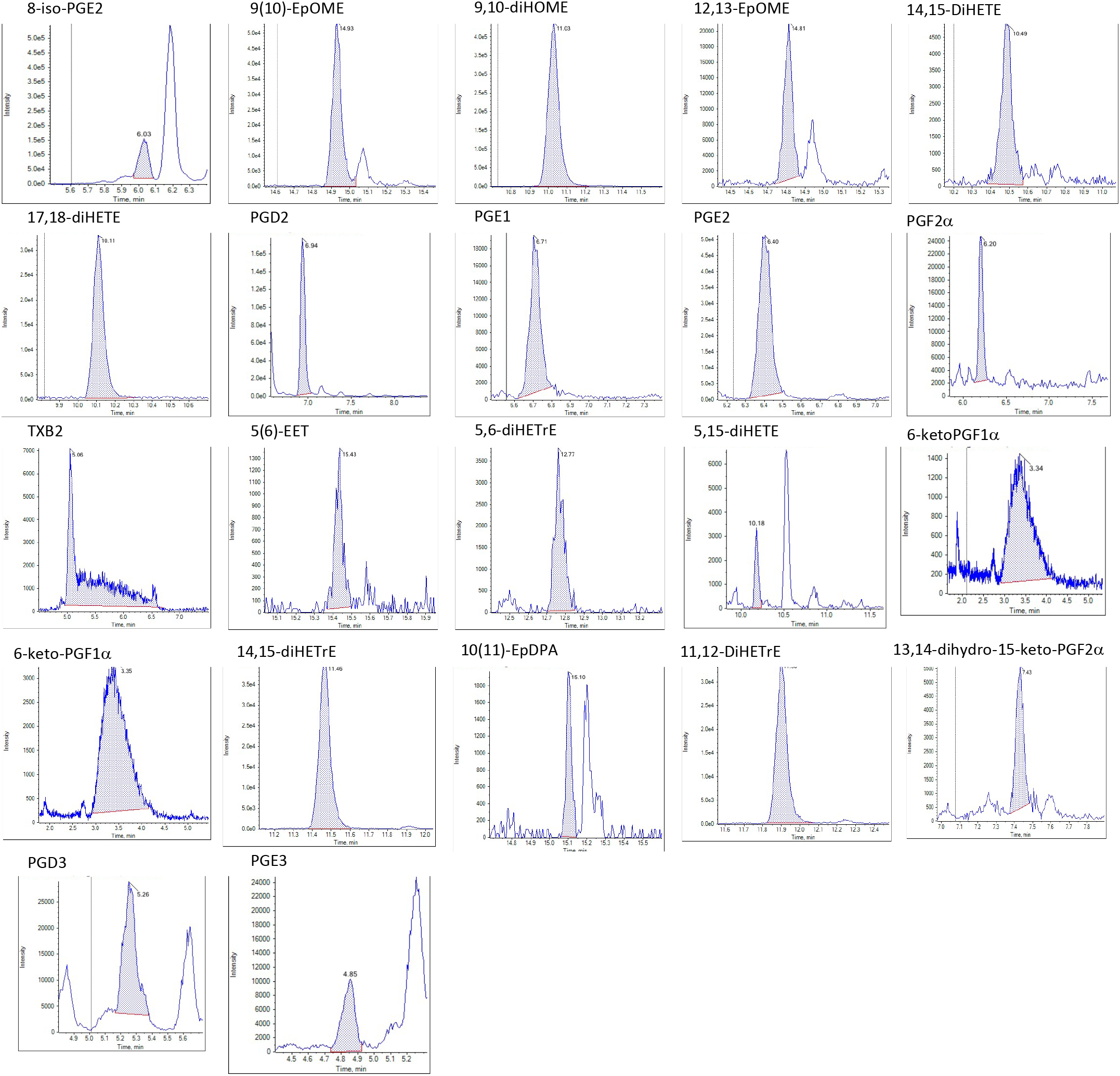
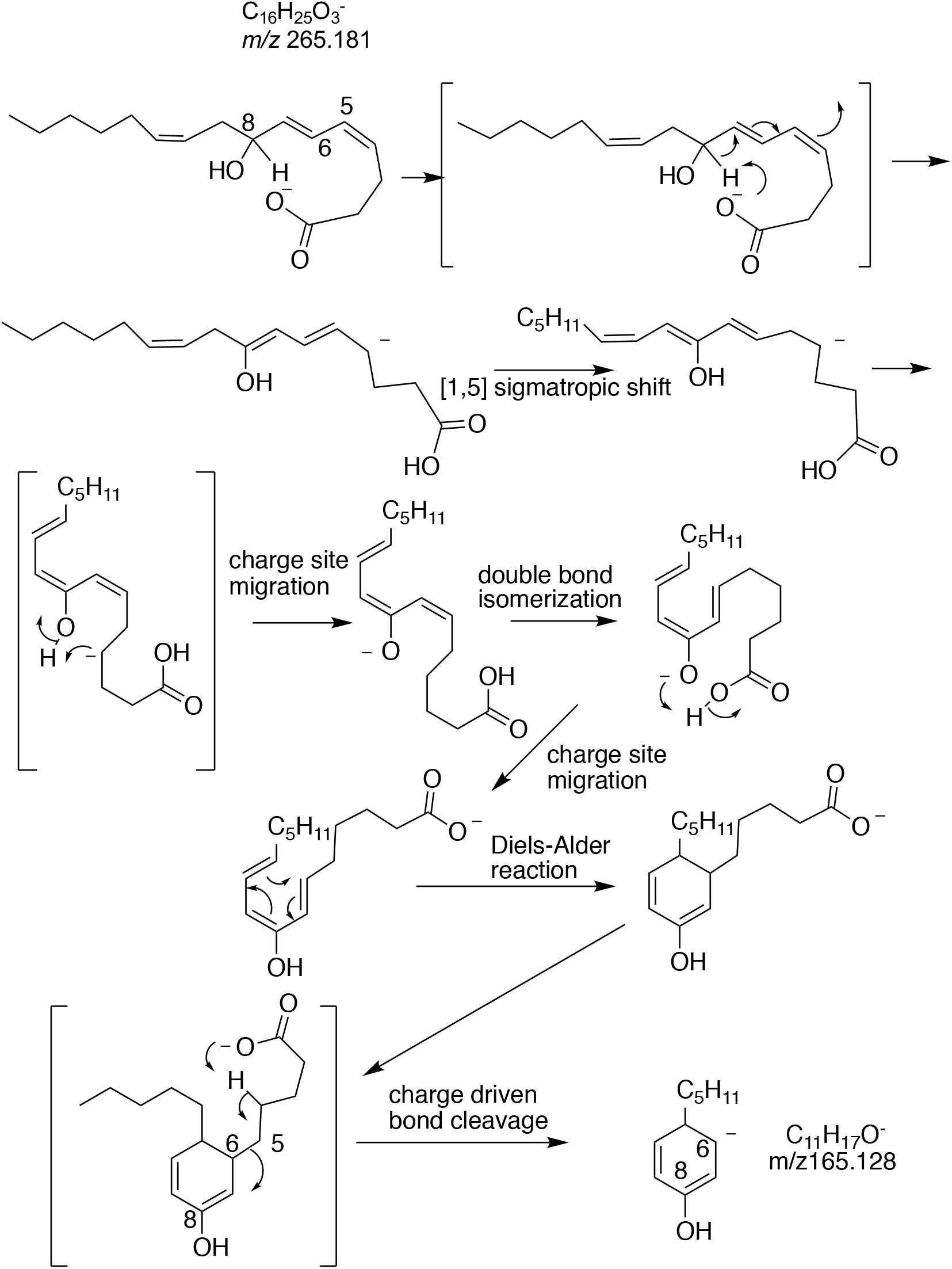
Representative chromatograms from oxylipin analysis showing typical peaks detected. **Supplementary Scheme 1. Proposed MS/MS for 8S-hydroxy-4Z,6E,10Z-hexadecatrienoic acid (tetranor triene).**

**Supplementary Figure 13.**
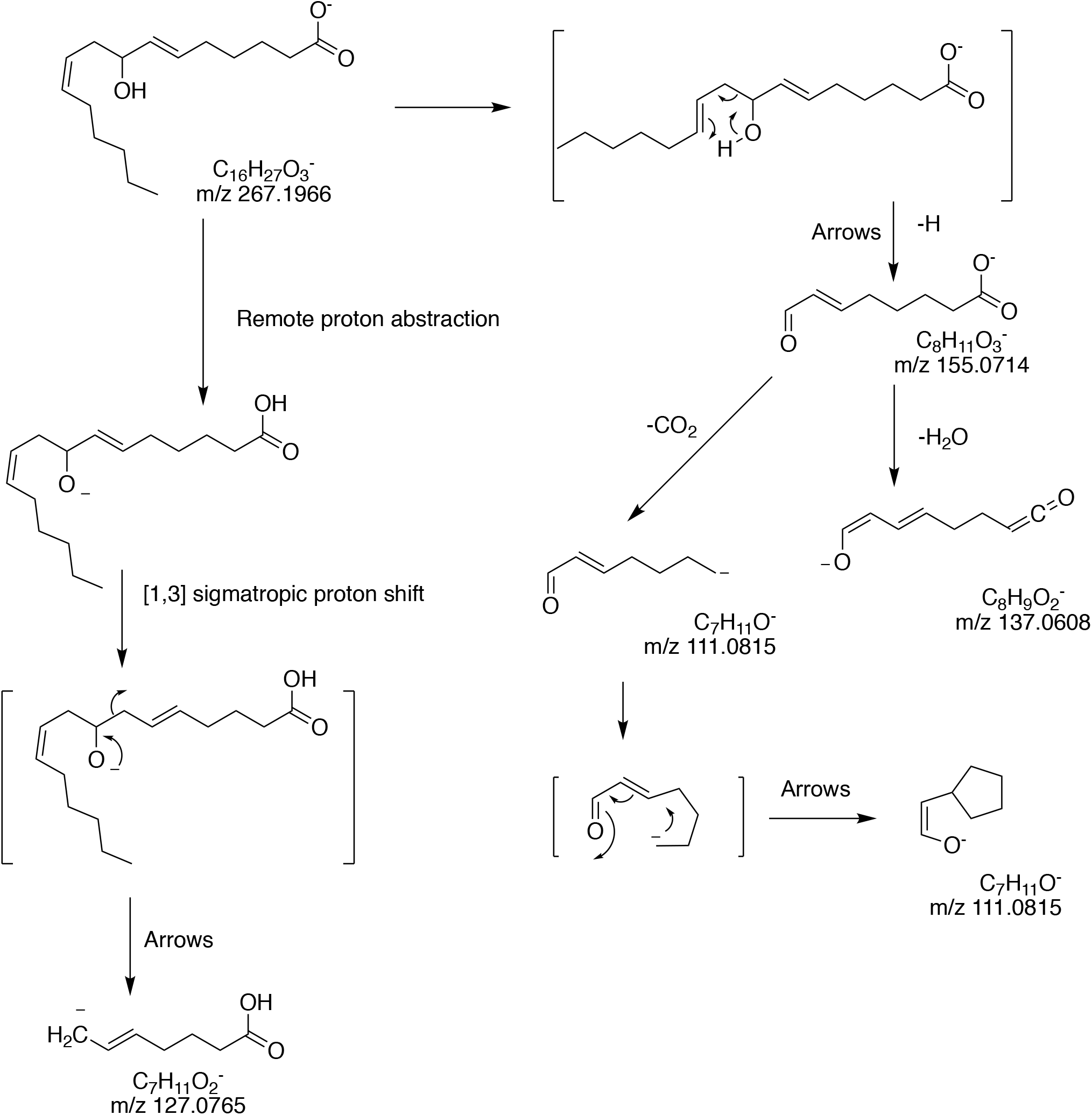
Supplementary Scheme 2. Proposed MS/MS for 8S-hydroxy-6E,10Z-hexadecadienoic acid (tetranor diene). The term “Arrows” is used to denote each of the steps where little one-sided arrows suggest electrons might move

**Supplementary Figure 14.**
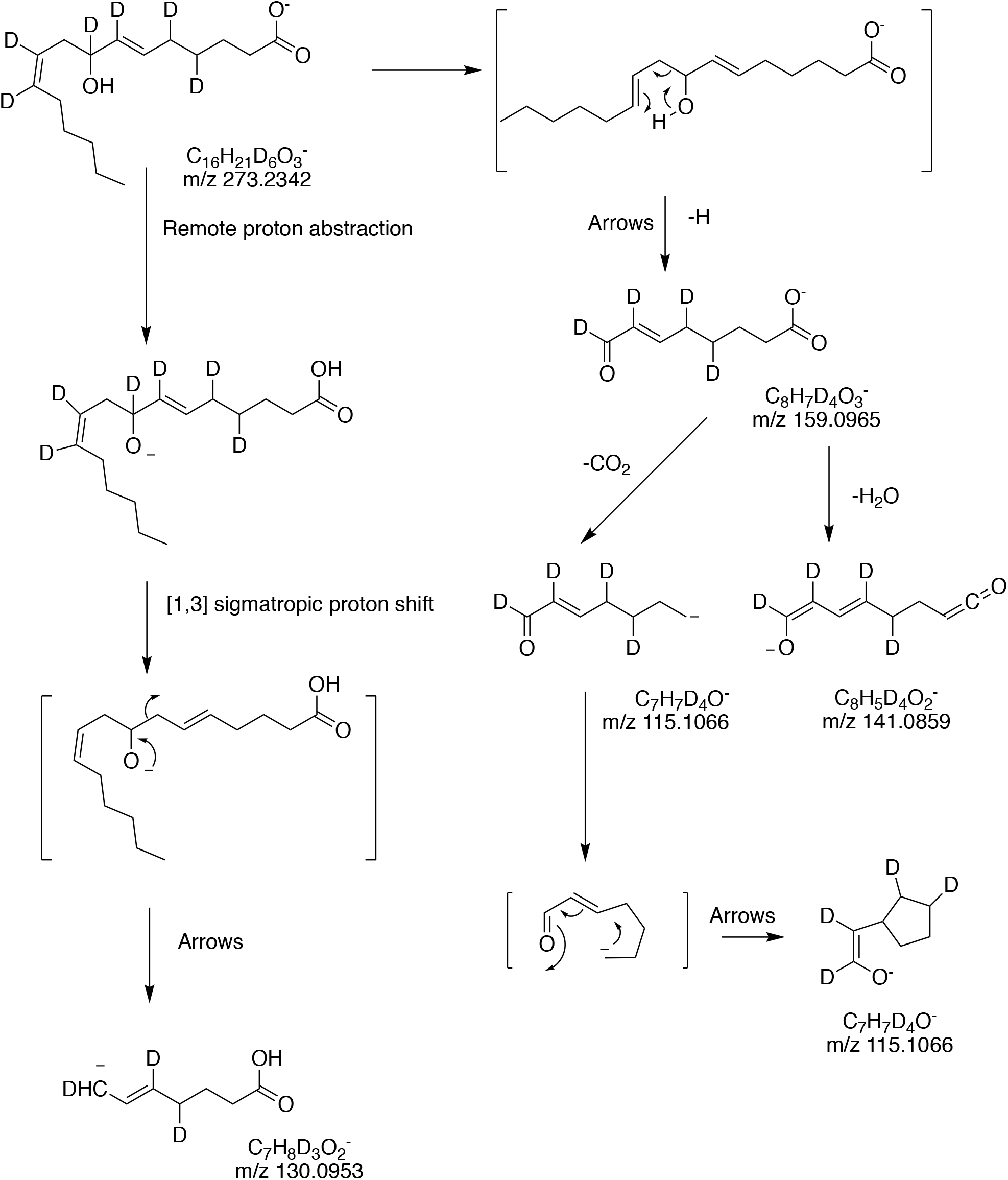
Supplementary Scheme 3. Proposed MS/MS for D_6_-8S-hydroxy-6E,10Z-hexadecadienoic acid (tetranor diene).

**Supplementary Figure 15.**
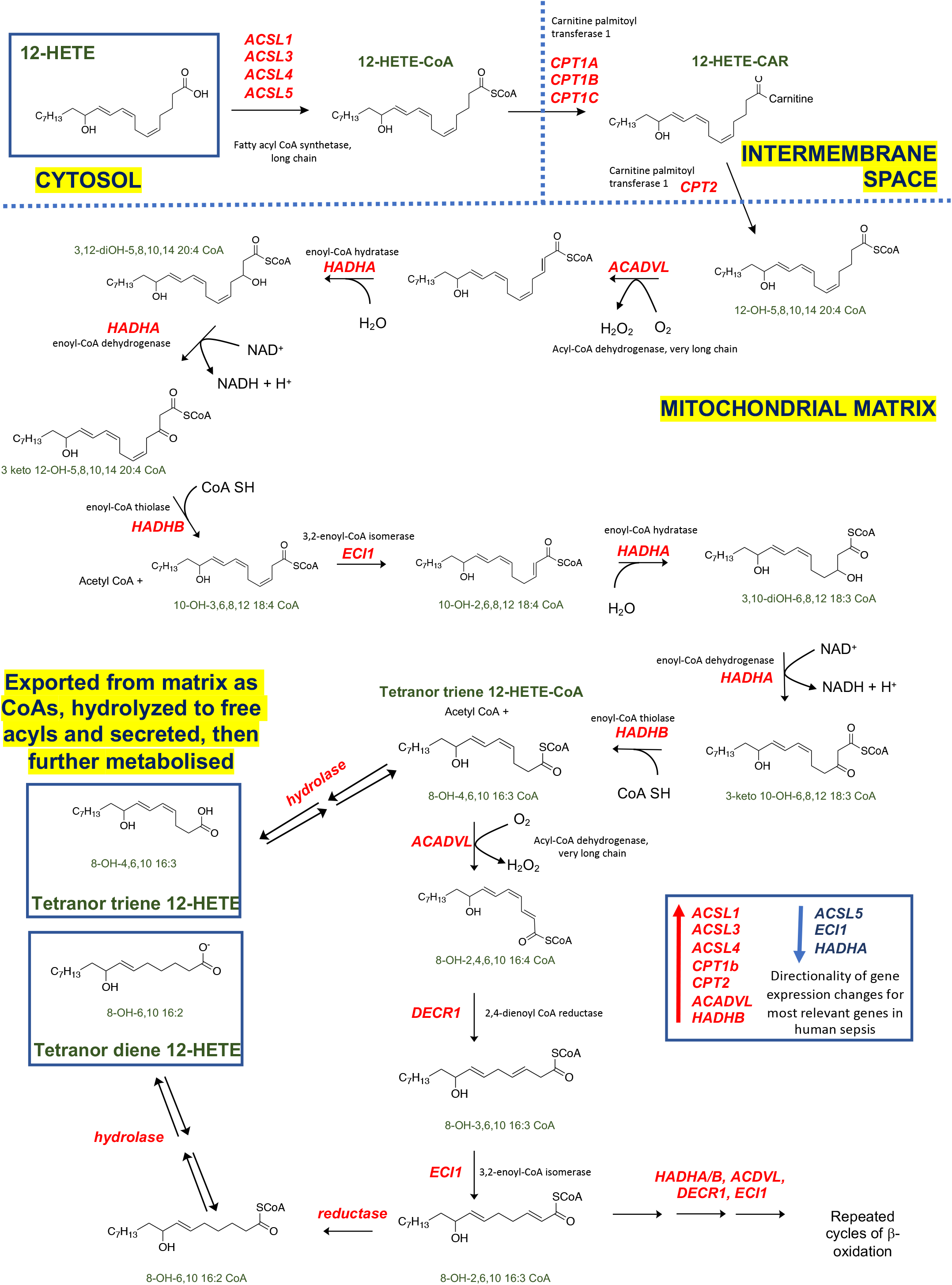
Proposed biochemical mechanism of β-oxidation for 12-HETE.

**Supplementary Table 1.**

|  | ng/mouse |  |
| --- | --- | --- |
| median | low lipid | high lipid |
| 5-HETE | 0.20796525 | 0.4159305 |
| 8-HETE | 0.223796 | 0.447592 |
| 9-HETE | 0.08038325 | 0.1607665 |
| 11-HETE | 0.27483475 | 0.5496695 |
| 12-HETE | 7.3727075 | 14.745415 |
| 15-HETE | 1.03487875 | 2.0697575 |
| 20-HETE | 0.3686335 | 0.737267 |
| 11-HEPE | 0.02583275 | 0.0516655 |
| 12-HEPE | 0.27923875 | 0.5584775 |
| 15-HEPE | 0.52596975 | 1.0519395 |
| 4-HDOHE | 0.02050975 | 0.0410195 |
| 8-HDOHE | 0.050181 | 0.100362 |
| 10-HDOHE | 0.1987765 | 0.397553 |
| 11-HDOHE | 0.059643 | 0.119286 |
| 13-HDOHE | 0.1355335 | 0.271067 |
| 14-HDOHE | 10.580795 | 21.16159 |
| 16-HDOHE | 0.037497 | 0.074994 |
| 17-HDOHE | 3.375567 | 6.751134 |
| 20-HDOHE | 0.02456675 | 0.0491335 |
| 9-HODE | 4.63153975 | 9.2630795 |
| 13-HODE | 31.3015975 | 62.603195 |
| 9-HOTrE | 0.37260025 | 0.7452005 |
| 13-HOTrE | 11.3827 | 22.7654 |
| 15-HETrE | 0.138236 | 0.276472 |
| 9-OxoODE | 0.676628 | 1.353256 |
| 13-OxoODE | 1.7746715 | 3.549343 |
| 12-OxoETE | 1.64698925 | 3.2939785 |
| 15-OxoETE | 0.2720885 | 0.544177 |
| 9,10-DIHOME | 2.03603825 | 4.0720765 |
| 12,13-DIHOME | 5.28231 | 10.56462 |
| 5,6-DIHETrE | 0.05350925 | 0.1070185 |
| 8,9-DIHETrE | 0.169896 | 0.339792 |
| 11,12-DIHETrE | 0.2299905 | 0.459981 |
| 14,15-DIHETrE | 0.3937595 | 0.787519 |
| 8,15-DIHETE | 0.027859 | 0.055718 |
| 14,15-DIHETE | 0.6634135 | 1.326827 |
| 17,18-DIHETE | 1.1895955 | 2.379191 |
| 6-trans LTB4 | 0.0572235 | 0.114447 |
| Mar-01 | 0.03122825 | 0.0624565 |
| 9(10)-EpOME | 0.52882825 | 1.0576565 |
| 12(13)-EpOME | 1.18597275 | 2.3719455 |
| 5(6)-EET | 0.17895925 | 0.3579185 |
| 10(11)-EpDPA | 0.00502125 | 0.0100425 |
| 19(20)-EpDPA | 0.5217775 | 1.043555 |
| PGD2 | 0.053451 | 0.106902 |
| PGE1 | 0.02898425 | 0.0579685 |
| PGE2 | 0.29952575 | 0.5990515 |
| 13,14-dihydro-15-keto PGD2 | 0.0087255 | 0.017451 |
| 8-iso PGE2 | 0.039122 | 0.078244 |
| PGF2α | 0.02432725 | 0.0486545 |
| 6-keto PGF1α | 1.00629425 | 2.0125885 |
| TXB2 | 0.188358 | 0.376716 |

**Supplementary Table 2.**
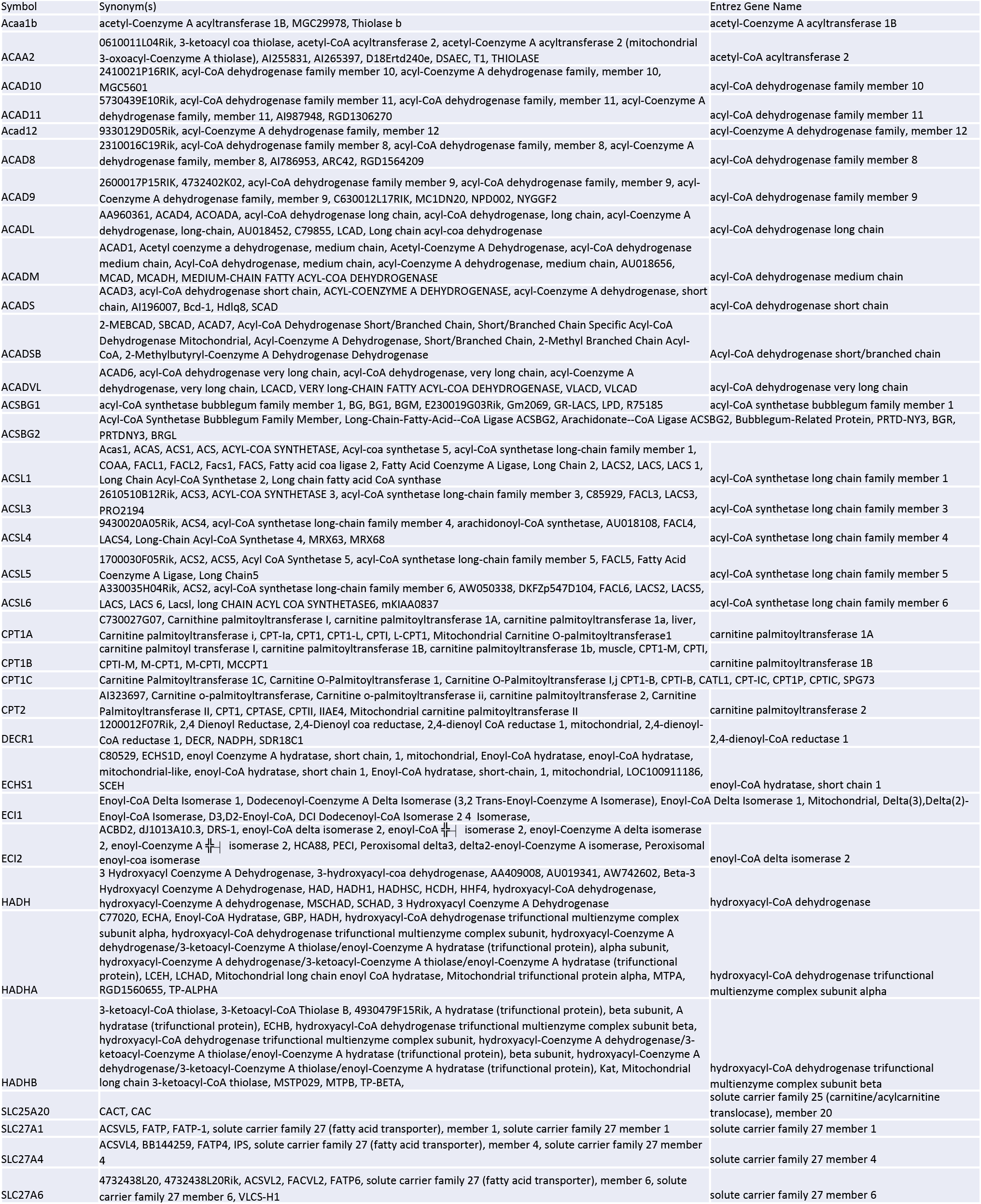
Putative genes relevant to micothondrial uptake and β-oxidation.

**Supplementary Table 3.**
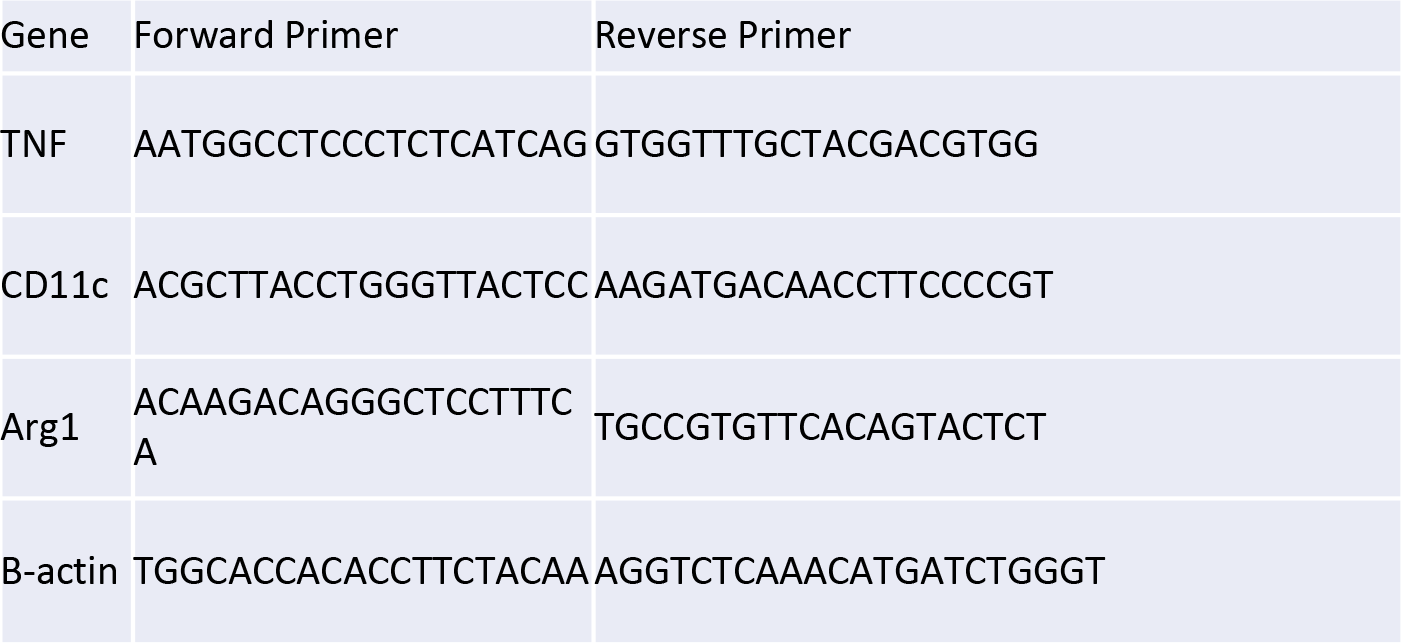
Primers used for macrophage polarization analysis.

**Supplementary Table 4.**
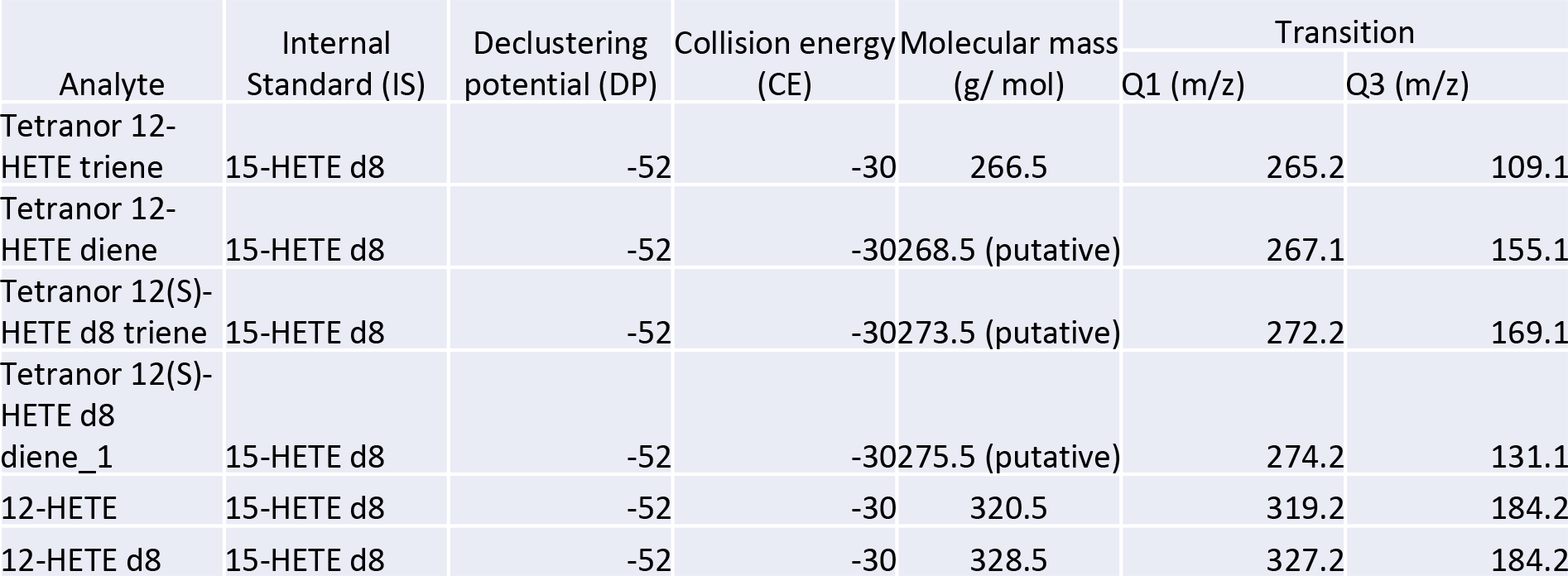
MRM and instrument parameters for HETE metabolites.

| Analyte | Internal Standard (IS) | Declustering potential (DP) | Collision energy (CE) | Molecular mass (g/ mol) | Transition |  |
| --- | --- | --- | --- | --- | --- | --- |
|  |  |  |  |  | Q1 (m/z) | Q3 (m/z) |
| Tetranor 12-HETE triene | 15-HETE d8 | -52 | -30 | 266.5 | 265.2 | 109.1 |
| Tetranor 12-HETE diene | 15-HETE d8 | -52 | -30 | 268.5 (putative) | 267.1 | 155.1 |
| Tetranor 12(S)-HETE d8 triene | 15-HETE d8 | -52 | -30 | 273.5 (putative) | 272.2 | 169.1 |
| Tetranor 12(S)-HETE d8 diene_1 | 15-HETE d8 | -52 | -30 | 275.5 (putative) | 274.2 | 131.1 |
| 12-HETE | 15-HETE d8 | -52 | -30 | 320.5 | 319.2 | 184.2 |
| 12-HETE d8 | 15-HETE d8 | -52 | -30 | 328.5 | 327.2 | 184.2 |

**Supplementary Table 5.**
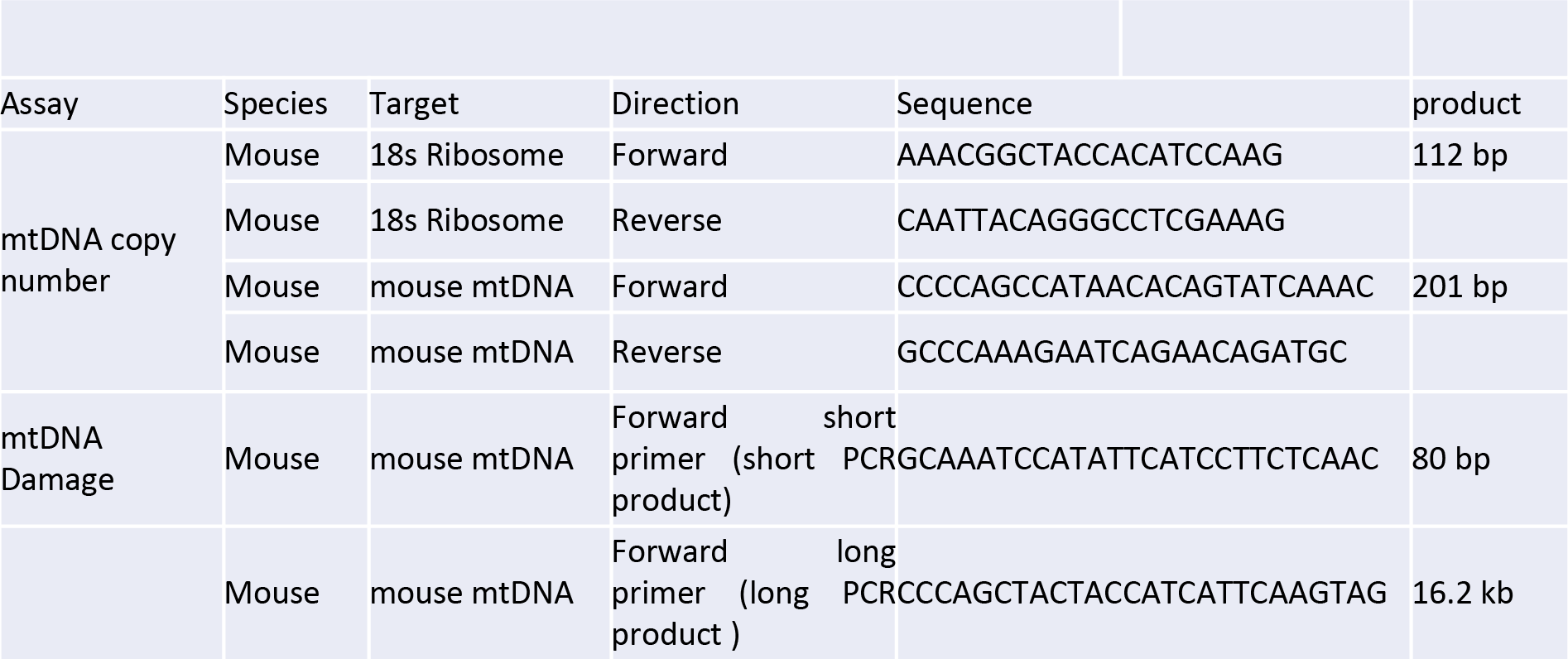
mtDNA primers for copy number assessment.

**Supplementary Table 6.** MRM and instrument parameters for oxidized phospholipids.

| MRM | Assignment | Retention Time (mins) |
| --- | --- | --- |
| 738.6/179.1 | PE P-16:0_12-HETE | 12.40 |
| 754.6/179.1 | PE 16:0_12-HETE | 11.80 |
| 764.6/179.1 | PE P-18:1_12-HETE | 12.63 |
| 766.6/179.1 | PE P-18:0_12-HETE | 13.96 |
| 780.6/179.1 | PE 18:1_12-HETE | 12.05 |
| 780.6/179.1 | PC 16:1_12-HETE | 10.51 |
| 782.6/179.1 | PE 18:0_12-HETE | 13.31 |
| 782.6/179.1 | PE 16:0_12-HETE | 11.72 |
| 808.7/179.1 | PE 18:1_12-HETE | 11.95 |
| 810.7/179.1 | PE 18:0_12-HETE | 13.27 |

**Supplementary Table 7.**
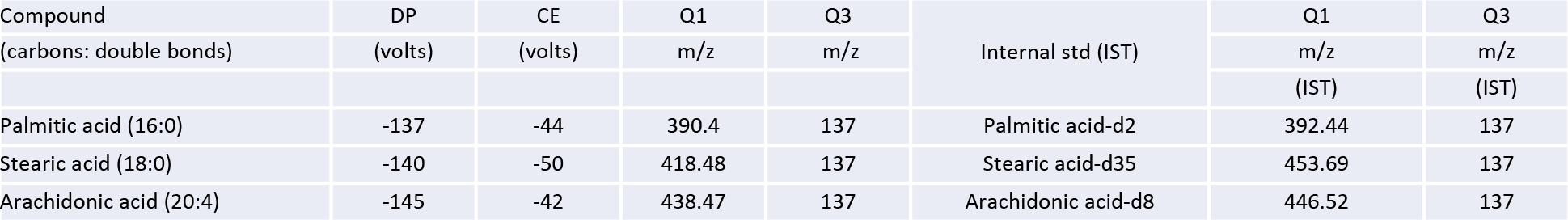
MRM transitions and MS parameters for FFA and the deuterated internal standards utilised (same MS settings)

| Compound | DP | CE | Q1 | Q3 | Internal std (IST) | Q1 | Q3 |
| --- | --- | --- | --- | --- | --- | --- | --- |
| (carbons: double bonds) | (volts) | (volts) | m/z | m/z |  | m/z | m/z |
|  |  |  |  |  |  | (IST) | (IST) |
| Palmitic acid (16:0) | -137 | -44 | 390.4 | 137 | Palmitic acid-d2 | 392.44 | 137 |
| Stearic acid (18:0) | -140 | -50 | 418.48 | 137 | Stearic acid-d35 | 453.69 | 137 |
| Arachidonic acid (20:4) | -145 | -42 | 438.47 | 137 | Arachidonic acid-d8 | 446.52 | 137 |

